# Improved nylon polymerization using amide diads

**DOI:** 10.1101/2025.02.28.640842

**Authors:** Liangyu Qian, Isaiah T. Dishner, Dana L. Carper, Vilmos Kertesz, Nicholas T. Zolnierczuk, Nduka D. Ogbonna, Nikki A. Thiele, John F. Cahill, Jeffrey C. Foster, Joshua K. Michener

## Abstract

Nylons, a major class of synthetic polyamides, are widely used due to their excellent mechanical strength, thermal stability, and chemical resistance. Conventional nylon production relies on the polymerization of lactams or stoichiometric nylon salts. However, applying these approaches to unconventional precursors such as bio-derived glutaric acid produces polymers with low molecular weight and limited applications. To address these challenges, we demonstrated that chemically-synthesized nylon diads enable the production of higher-molecular-weight polyamides compared with traditional salts. We then identified a biosynthetic approach using amide synthetases to convert unprotected bifunctional substrates into nylon-relevant diads. Using a cofactor regeneration system, enzymatic diad synthesis was scaled to produce sufficient material for laboratory-scale characterization and solid-state polymerization. Amide synthetases demonstrated broad substrate scope, catalyzing the regioselective assembly of diverse nylon-relevant diacids, diamines, and ω-amino acids. This strategy offers a novel route to synthesize challenging nylon monomers and advances production of bioderived nylons.

## Introduction

Polyamides are polymers linked by amide groups and synthesized by the reaction of diamines with dicarboxylic acids, by the condensation of ω-amino acids, or by ring-opening polymerization of cyclic lactams. Nylons, a prominent category of synthetic thermoplastic polyamides, were produced at an estimated scale of 9 million tons in 2020 (Global Industry Analysts, 2020). Applications of nylons are widespread due to their exceptional mechanical strength, thermal stability and chemical resistance, making them indispensable in industries such as textiles, automotive manufacturing, and additive manufacturing. The continued demand for high-performance materials has driven research efforts aimed at improving the synthesis and functional properties of nylons, with an emphasis on achieving higher molecular weight and expanding the scope of usable precursors.^1–4^

Commercial nylons are typically produced by polymerizing either a single ω-amino acid (e.g., nylon-6, also known as PA6) or a stoichiometric nylon salt formed from a diamine and a diacid (e.g., nylon-66, or PA66). Efforts to improve nylon sustainability have led to interest in incorporating bioderived components such as glutarate and cadaverine.^5–13^ However, polymerization of nylon salts containing glutarate often yields polymers with low molecular weight, which has hindered their broader exploration and application.^5,14^ Despite the limited studies on glutarate-based nylons, PA55 composed of cadaverine and glutarate has been reported to possess desirable properties, including a high melting temperature and excellent thermal stability.^14^ PA65 remains largely unexplored but shows a distinct hydrogen bonding pattern and moderate crystallinity.^5,15^ More flexible synthetic routes that can readily incorporate non-conventional monomers could expand the scope of potential polyamides.

As an alternative synthesis route, polymerization of the PA66 diad, a product of enzymatic PA66 hydrolysis, was shown to yield a polymer with a higher molecular weight than the corresponding nylon salt.^16^ However, it was unclear whether other nylon diads would similarly outperform their respective salts. Furthermore, while oligomeric products can be formed from polyamide hydrolysis, direct diad synthesis is challenging. Peptide synthesis via solid-phase methods is well-established, but it involves expensive protection and deprotection steps to control chain length, sequence and reactivity, as well as extensive use of organic solvents.^17^ Alternatively, enzymatic synthesis could potentially offer a more efficient and environmentally friendly route for generating these intermediates. A biosynthetic approach would be particularly useful if combined with a pathway for biosynthesis of nylon-relevant monomers, since production of zwitterionic diads would avoid the cost of separately synthesizing, neutralizing, and purifying diacids and diamines.^18^

Biology provides several possible routes for amide bond formation. Ribosomes can synthesize amide bonds between natural α-amino acids. However, while ribosomes can generate polyamides with high molecular weight and exquisite sequence control, expanding the monomer scope beyond canonical α-amino acids is challenging.^19–25^ Additionally, non-ribosomal peptide synthetases are multimodular enzymes that offer greater flexibility in monomer scope and bond formation but are difficult to express and engineer.^26–28^ Finally, amide synthetases can catalyze amide bond formation in small molecules by using ATP to adenylate a carboxylic acid, which is then condensed with an amine.^29,30^ These enzymes natively function in the biosynthesis of complex secondary metabolites such as antibiotics and siderophores and have been mainly repurposed for the production of pharmaceuticals.^31–35^ They are relatively small (400-500 amino acids), easy to express and purify, and active in heterologous hosts.

For example, the amide synthetase McbA is known for its broad substrate specificity.^36^ However, McbA has been shown to have low activity with aliphatic substrates, suggesting that it may not accept nylon-relevant monomers.^31^ Similarly, the amide synthetase SfaB accepts a broad range of short-chain fatty acids and catalyzes amidation or thioesterification with various amine or thiol nucleophiles, but reportedly does not tolerate dicarboxylic acids such as those used in nylons.^37^ Therefore, the application of natural or engineered amide synthetases for the synthesis of nylon building blocks remains largely unexplored.

In this study, we showed that chemically-synthesized nylon diads can be used to generate a range of polyamides with higher molecular weights compared to polymerization of traditional salts. Motivated by these findings, we then demonstrated a simplified biosynthetic strategy using amide synthetases to produce nylon diad precursors from unprotected bifunctional substrates. By coupling amide bond formation with an ATP-regeneration system, we optimized and scaled up this biocatalytic process and established it as a viable method for synthesizing nylon diads. Furthermore, we showed that amide synthetases exhibit a broad substrate scope and can catalyze the regioselective assembly of a diverse range of nylon-relevant diacids, diamines, and ω-amino acids. This approach provides a facile route for the polymerization of challenging monomers and expands the potential for tailoring nylon properties through enzymatically derived precursors.

## Results and Discussion

### Diad polymerization and characterization

To extend the previous results showing the advantages of diad-based precursors in PA66 synthesis,^16^ we tested whether additional diads, particularly those incorporating glutarate, could also increase the molecular weight of the resulting polymer compared to the corresponding nylon salt (Figure 1A). These diads were prepared from combinations of cadaverine (**C**) or hexamethylenediamine (**M**) as the diamine, and succinic acid (**S**), glutaric acid (**G**), or adipic acid (**A**) as the carboxylic acid (See Figure S1 for full list of substrate codes). We previously synthesized the **MA** diad by coupling *N*-Boc-1,6-hexamethylenediamine with monomethyl adipate using EDC as the coupling agent.^16^ Although we initially attempted to synthesize the diads directly from unprotected diamines using the procedure reported by Madhavachary *et al*.,^38^ we ultimately found this method overly cumbersome as it required complex chromatographic separation to completely remove the difunctionalized byproduct. To this end, **CS, MS, CG, MG, CA**, and **MA** (Figure 1B) were synthesized through conventional methods to provide diads in sufficient quantity for polymerization (see Supplemental Methods and Figures S2-S37 for details). These diads were polymerized by the solid-state polymerization (SSP) protocol described in our previous work.^16^ Number average and weight average molecular weights (*M*_n_ and *M*_w_, respectively) of the resulting polymers were measured by size exclusion chromatography (SEC), and *M*_w_ values were compared to polyamides prepared from traditional nylon salts (Figure 1C). These data showed diads containing adipic acid or glutaric acid moieties produced polyamides with significantly higher *M*_w_ than those synthesized from their equivalent nylon salts. Molecular weight data was not recorded for PA55 synthesized from the **CG** diad because the resulting polymers were insoluble in HFIP. We hypothesized the difference in *M*_w_ could be attributed to multiple factors, most notably by improving the stoichiometric balance of the diamine and diacid components, minimizing evaporation of volatile hexamethylenediamine, and starting the polymerization with 50% of the potential amide bonds preformed.

**Figure 1:**
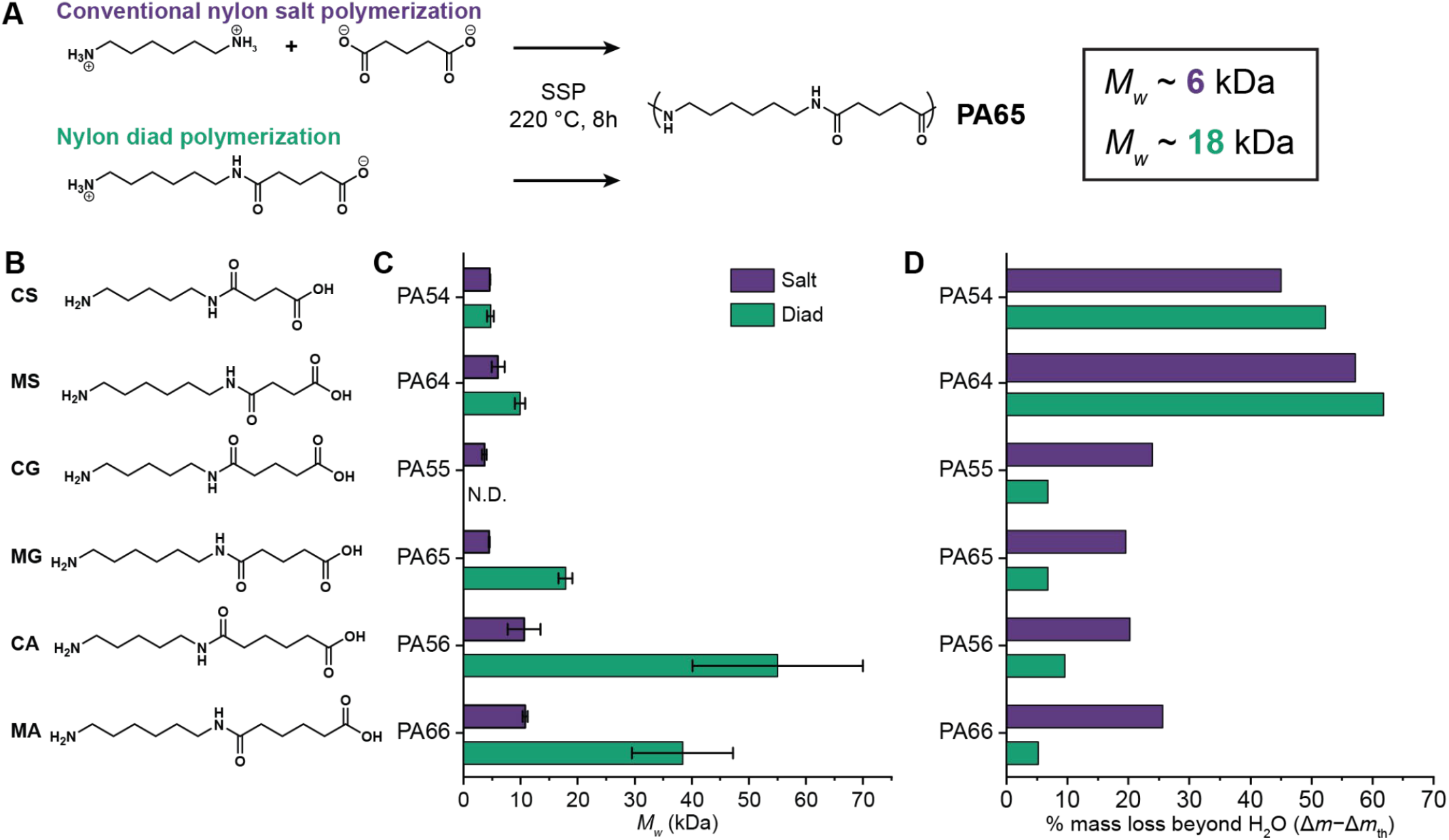
Nylon diads yield polyamides with high molecular weight. A) Comparison of PA65 synthesized by solid state polymerization (SSP) from conventional nylon salt or equivalent nylon diad. B) Chemical structures of nylon diads. **CS**: cadaverine-succinic acid diad; **MS**: hexamethylenediamine - succinic acid diad; **CA**: cadaverine - adipic acid diad; **CG**: cadaverine - glutaric acid diad; **MG**: hexamethylenediamine - glutaric acid diad; **CA**: cadaverine - adipic acid diad; **MA**: hexamethylenediamine - adipic acid diad. C) Comparison of weight-averaged molecular weights (*M*_w_) of polyamides prepared from diads and their corresponding nylon salts through solid-state polymerization. *M*_w_ data is not reported for **CG** because the polymers synthesized from this diad were insoluble in HFIP. Sample size is *N* = 3. Error bars are shown as the mean ± standard deviation. D) Excess mass loss as determined from isothermal TGA studies. Excess mass loss is calculated as the difference between the experimentally determined mass loss and the expected mass loss assuming 100% monomer conversion.

To test these hypotheses, we used isothermal thermogravimetric analysis (TGA) to monitor the change in sample mass under simulated SSP conditions. Diad and nylon salt samples were initially conditioned at 100 °C for 1 h to remove any residual solvent and/or moisture, and subsequently polymerized by heating to 220 °C for 8 h (Figures S40-S45). As shown in Figure 1D, most nylon salts lose a considerably larger portion of their initial mass than their diad counterparts over the course of the polymerization, even when accounting for differences in expected mass loss caused by the evolution of water. Moreover, the majority of this mass loss occurs at relatively low monomer conversion (i.e. within approximately the first 30 min). These results imply that evaporation of diamine and/or diacid components of the nylon salts creates a stoichiometric imbalance which limits maximum *X*_n_ in accordance with Carother’s equation.^39^ Exceptionally high mass losses were observed during the polymerization of succinic acid-containing PA64 and PA54 samples regardless of whether the diad or nylon salt starting materials were used (Figures S44 and S45). This mass loss was ultimately attributed to competing imidization pathways which can create volatile byproducts when imidization occurs adjacent to a polymer chain end. Imide formation was clearly visible from Fourier-transform infrared (FT-IR) spectroscopy of the PA64 and PA54 polymers, which showed an imide C=O stretch at ∼1770 cm^-1^ (Figure S46). Thermal imidization also explains the relatively low MW of PA64 and PA54 samples since imidization imbalances the ratio of carboxylic acid to amine chain ends and also acts as a form of ring-chain equilibrium which further limits *X*_n_.40

To compare the physical properties of the polyamides prepared from diads with those prepared from the corresponding nylon salt, we used thermogravimetric analysis (TGA) and differential scanning calorimetry (DSC) to measure the degradation temperatures (*T*_d_), glass transition temperatures (*T*_g_), melting temperatures (*T*_m_), and enthalpies of melting (ΔH_m_) (Figures S47-S52 and Table S2). No considerable difference was observed in *T*_d_ for PA66, PA56, PA65, PA55, or PA54 prepared from either nylon salts or diads. However, a somewhat lower *T*_d_ was observed for PA64 synthesized from **MS** compared to its corresponding salt (Figure S51A). Additionally, PA64 prepared from **MS** showed an apparent two step decomposition profile, whereas the salt-derived polyamide decomposed in a single step. Despite this difference in thermal stability, both PA64 samples display nearly identical thermal properties, consistent with the hypothesis that the two samples are chemically identical. Likewise, similar agreement in *T*_g_, *T*_m_, and ΔH_m_ for salt-derived and diad-derived polyamides were observed for PA66, PA65, and PA54. In combination, we found that diads are preferred feedstocks for nylon polymerization, since they yield polyamides that are chemically identical but have higher molecular weights compared to the corresponding salt-derived polyamides.

### Amide synthetases can synthesize nylon diads

Motivated by findings that diads are promising precursors for nylon polymerization, we sought to investigate more efficient routes for diad synthesis. To overcome the need for protecting groups in chemical diad synthesis, we explored a biosynthetic route using amide synthetases to couple unprotected substrates (Figure 2B). To evaluate the activities of amide synthetases on nylon building blocks, five enzymes (DdaG, SfaB, SuCphA1, McbA and PylC) reported to catalyze isopeptide bond formation were heterologously expressed in *E*.*coli* BL21(DE3). Among these, four enzymes (DdaG, SfaB, SuCphA1 and McbA) were successfully purified as soluble hexahistidine-tagged proteins (Figures S53-S54), while His-tagged PylC was insoluble. To enhance PylC solubility, a SUMO tag was added to the C-terminal of the hexahistidine tag, resulting in a soluble His-SUMO-tagged PylC (Figure S55).^41^ We first confirmed the reported activities of DdaG to ligate 2,3-diaminopropionate with fumarate,^35^ SfaB to ligate 5-chlorovaleric acid with cadaverine,^37^ and SuCphA1 to ligate aspartate with arginine.^42^ However, the His-SUMO-tagged PylC failed to exhibit its native activity of ligating lysine and ornithine.^43^ Attempts to cleave the His-SUMO tag from the recombinant PylC were unsuccessful. Therefore, we next conducted *in vitro* biochemical assays using the soluble enzymes (DdaG, SfaB, SuCphA1, and McbA) with nylon constituents (adipic acid, hexamethylenediamine, succinic acid, cadaverine, glutaric acid) to assess and quantify the production of the respective diads (**CS, MS, CG, MG, CA** and **MA**) by comparison to chemically-synthesized standards using direct injection immediate droplet-on-demand/open port sampling interface-mass spectrometry (IDOT/OPSI-MS) (Figures 2C and S56).

**Figure 2:**
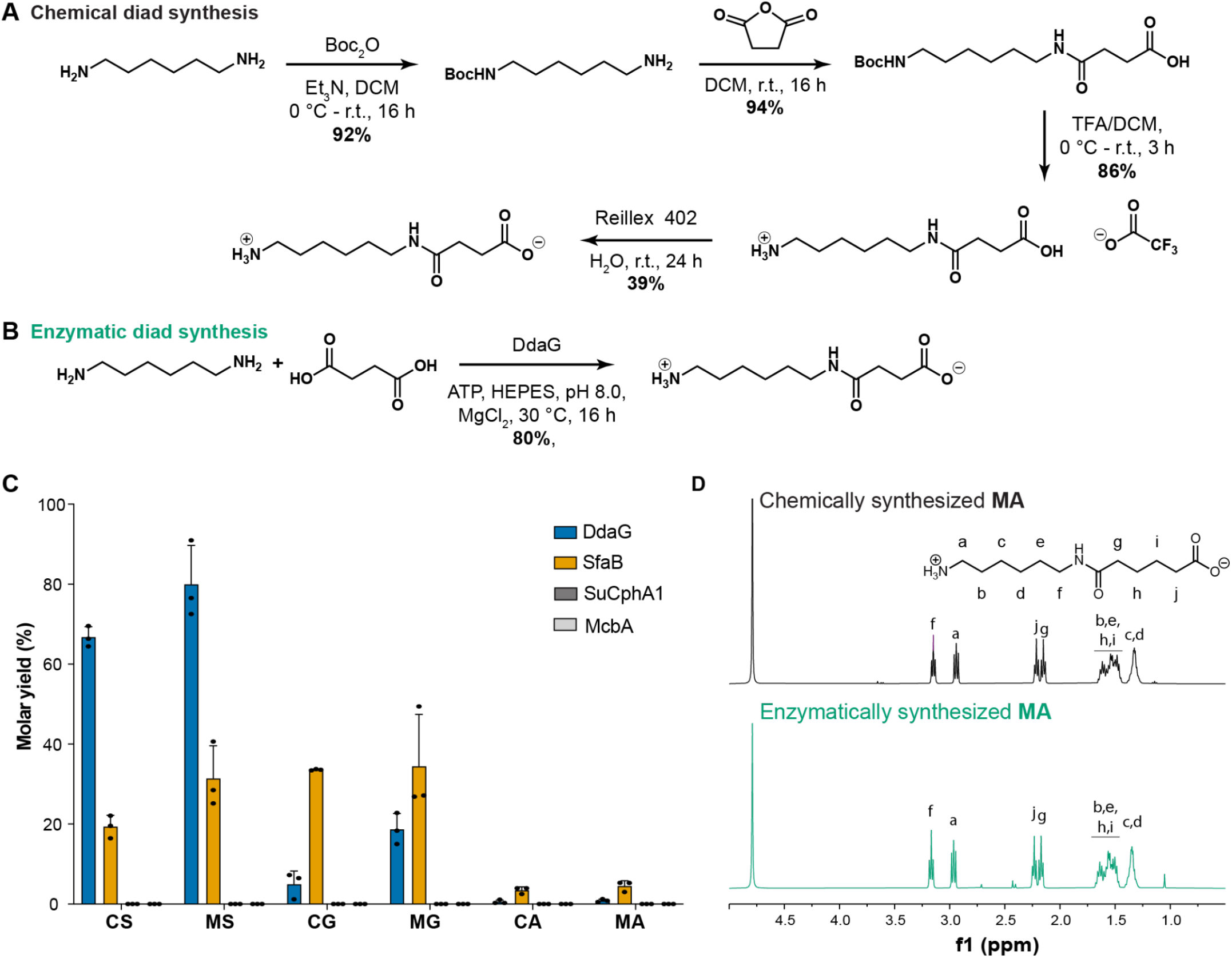
Diad quantification. A) Chemical synthesis of the hexamethylene diamine – succinic acid (**MS**) diad. B) Enzymatic synthesis of the **MS** diad. C) Small-scale quantification of enzymatically synthesized nylon diads. *In vitro* biochemical assays were conducted by incubating different diacids (i.e., succinic acid, glutaric acid, or adipic acid) with diamines (i.e., cadaverine or hexamethylenediamine) in the presence of enzymes (i.e., DdaG, SfaB, SuCphA1 and McbA). Products were quantified by comparing to the chemically-synthesized standards using I.DOT/OPSI-MS. Sample size is *N* = 3. Error bars are shown as the mean ± standard deviation. D) ^1^H NMR spectra of enzymatic-synthesized and chemically-synthesized **MA** diad.

Encouragingly, DdaG and SfaB showed significant activities and distinct specificities for nylon diad synthesis. The molar yields after overnight incubation ranged from 3% to 80%, with DdaG demonstrating the highest activity in synthesizing **MS** (∼80%) and **CS** (∼60%) but low production of **MG** and **MA**. In contrast, SfaB exhibited lower activity than DdaG for production of **MS** and **CS** but produced higher amounts of **MG** (∼20%) and **MA** (∼3%). We next scaled up **MA** production with SfaB. To confirm the structure of **MA**, we developed a purification protocol wherein the crude reaction mixture was reacted with 2-acetyldimedone (dde-OH) to provide a dde-functionalized adduct that is easily isolated chromatographically, and subsequently deprotected via hydrazinolysis (Scheme S1 and Figures S57 and S58). This allowed us to confirm the structure of enzymatically synthesized **MA** by comparing the ^1^H NMR spectra to the chemically synthesized standard (Figure 2D).

We did not observe detectable product formation in reactions with SuCphA1 or McbA, consistent with findings of a previous report that McbA does not accept aliphatic substrates.^31^ Interestingly, previous studies showed that SfaB cannot adenylate diacids for subsequent condensation with hydroxylamine, based on colorimetric detection of pyrophosphate released during adenylation.^37^ However, our observations reveal that SfaB can act on a range of diacids (i.e., succinic acid, glutaric acid and adipic acid) in combination with cadaverine and hexamethylenediamine. These differing results suggest that SfaB also exhibits specificity towards the amine acceptor.

### Scale up and optimization of diad biosynthesis

To demonstrate the utility of enzymatic synthesis for diad production, we next focused on optimizing conditions for scale-up and isolation, using synthesis of **MS** by DdaG as a model reaction. We increased the concentration of each substrate to 50 mM and increased the reaction volume to 50 mL, while keeping the concentration of DdaG at 10 μM. Based on OPSI-MS analysis, the reaction yield was 44%. The crude reaction mixture was then purified following the protocol previously described for **MA** to provide 60 mg (∼11%) of spectroscopically pure **MS** (Figures S59 and S60).

DdaG and SfaB are ATP-dependent amide synthetases, and providing super-stoichiometric ATP to ensure effective conversion is cost-prohibitive. As an alternative, we implemented an ATP recycling system using a purified type 2-III polyphosphate kinase (PPK12).^33,44,45^ In addition, we included inorganic pyrophosphatase from *E. coli* (ecPPase) to prevent potential inhibition by pyrophosphate. We then performed reaction optimization using **MS** production by DdaG. We iteratively optimized the concentrations of the substrates, polyphosphate, magnesium, AMP, and pH (Figures S61-65). Based on these results, the finalized optimal conditions for **MS** production were 50 mM succinic acid, 100 mM hexamethylenediamine, 100 mM MgCl_2_, 10 μM DdaG, 1 mg/mL PPK12, 5 mM AMP, 0.2 U ecPPase and 100 mM HEPES (pH 8). With these optimal conditions, we subsequently scaled up our reaction system with ATP regeneration from 50 μL to a 5 mL reaction volume. After 24 h incubation, a small volume (50 μL) of sample was analyzed by I.DOT/OPSI-MS, indicating a promising yield of ∼50% (∼5 g/L) and demonstrating a more than 3-fold improvement from the initial conditions (Figure 3).

**Figure 3:**
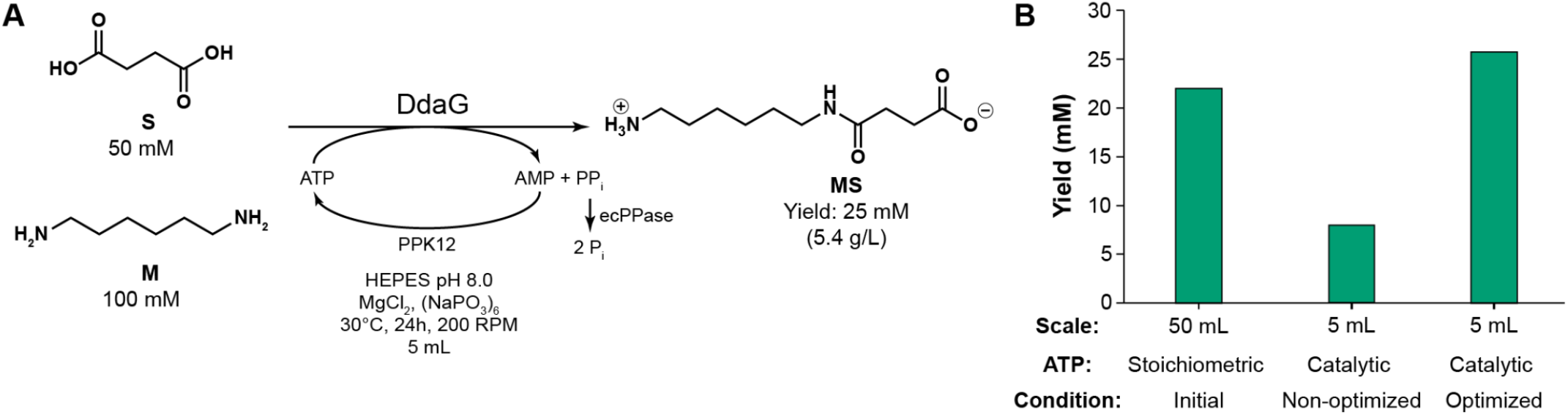
Diad scale up and optimization. A) ATP-recycled scale-up of **MS** production. A 5 mL reaction with increased substrate loading and an ATP regeneration system was performed, and the yield was quantified by mass spectrometry. B) Comparison of **MS** production under different approaches. The 50 mL reaction was performed with initial setup: 50 mM succinic acid, 50 mM hexamethylenediamine, 10 mM MgCl_2_, 10 μM DdaG, 50 mM ATP and 100 mM HEPES (pH 8). For the non-optimized 5 mL ATP-recycled reaction, 50 mM ATP was replaced with 1 mg/mL PPK12, 1 mM ATP, 1 mM AMP, 25 mM SHMP and 0.2 U ecPPase with other components unchanged. For the optimized condition, 100 mM hexamethylenediamine, 100 mM MgCl_2_ and 5 mM AMP were used with other components unchanged from the non-optimization. All the reactions were incubated for 24 hours at 30 ºC with shaking at 200 rpm. These experiments were conducted in *N*=1.

### Enzymatic synthesis of novel nylon diads

Since DdaG and SfaB exhibit activities in producing nylon-relevant diads, we next sought to explore their substrate range, which could provide opportunities to develop novel nylons. Therefore, we conducted *in vitro* biochemical assays using DdaG and SfaB with a diverse range of substrates, including traditional nylon constituents (adipic acid **A**, hexamethylenediamine **M**, and terephthalic acid **T**) and novel compounds varying in chain length, functional groups (e.g., thiol, ketone, and hydroxyl groups), aromaticity, and cyclicity, including dicarboxylic acids, diamines, and ω-amino acids (Figure S1).

We initially detected product formation using direct injection IDOT/OPSI-MS for untargeted characterization of enzymatic reactions.^46^ Using this approach, 41 out of 96 reaction combinations generated detectable products above the control using either DdaG or SfaB (Figures 4 and S66). To more accurately identify these products, we confirmed 24 structures through analysis of fragmentation patterns by MS^2^ (Figures S67-S90). Both enzymes displayed activities in ligating the carboxylic groups of the diacids with a wide spectrum of amine acceptors including linear diamines (cadaverine **C**, hexamethylenediamine **M**), an aromatic diamine (*p-*xylylenediamine **X**), a cyclic diamine (*cis*-1,4-cyclohexanediamine **N**), and the amine groups of ω-amino acids (6-aminohexanoic acid **H**).

**Figure 4:**
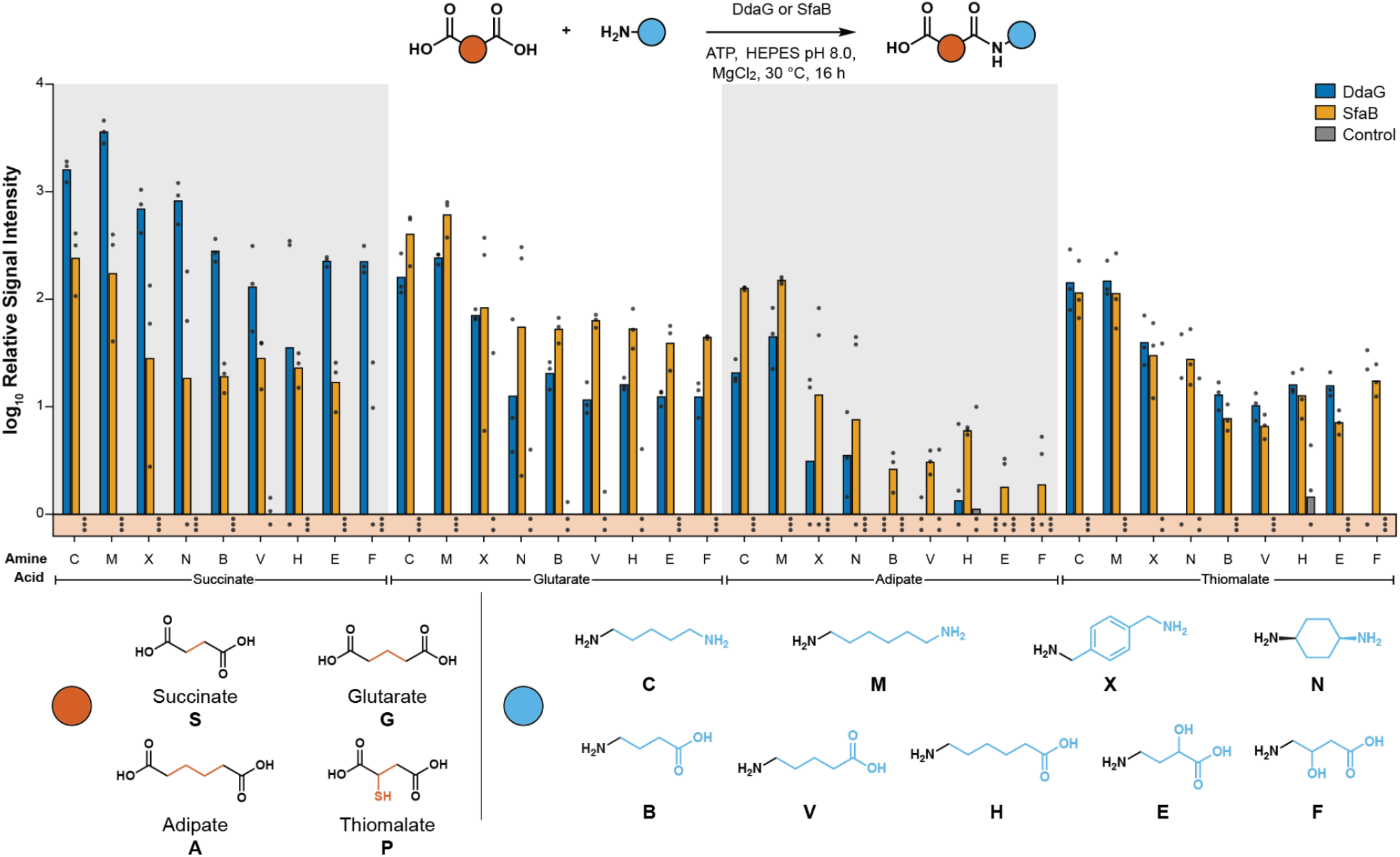
Amide synthetases have broad substrate range for potential nylon precursors. *In vitro* biochemical assays were conducted by incubating different diacids (i.e., succinic acid, glutaric acid, adipic acid and thiomalic acid) with diamines or ω-amino acids in the presence of enzymes (i.e., DdaG or SfaB) or no-enzyme control. Products were assayed using OPSI-MS. The log_10_MS signal intensity was cut off at 5.0 to filter out noise. **C**: cadaverine; **M**: hexamethylenediamine; **X**: *p*-xylylenediamine; **N**: *cis*-1,4-cyclohexanediamine; **B**: 4-aminobutyrate; **V**: 5-aminovalerate; **H**: 6-aminohexanoic acid; **E**: 4-amino-2-hydroxybutanoic acid; **F**: 4-amino-3-hydroxybutanoic acid. Sample size is *N* = 3. Data for **CS, MS, CG, MG, CA**, and **MA** shown in Fig. 2B are replotted here as unquantified MS intensities for comparison.

Notable differences between DdaG and SfaB were observed in their preference for carboxylic acid donors, consistent with our quantification of their activity in producing more conventional nylon-relevant diads. DdaG exhibited excellent activity in ligating succinic acid (**S**) with a wide range of amine acceptors and showed lower activities with glutaric acid (**G**) and adipic acid (**A**). In contrast, SfaB displayed higher efficiency using **G** and **A** as carboxylic donors with diverse amine acceptors. In addition, DdaG can also catalyze reactions involving terephthalic acid (**T**) and α-ketoglutaric acid (**K**) with multiple diamines, while SfaB was inactive in these reactions. Alternatively, SfaB was capable of activating 1,4-cyclohexanedicarboxylic acid (**O**) with specific diamines (i.e., **M** and **X**), whereas DdaG showed negligible activity (Figure S66). It is also notable that both enzymes were unable to activate the carboxylic groups of ω-amino acids. Interestingly, we noticed both enzymes could activate diacids bearing additional functional groups such as thiomalic acid (**P**). The broad substrate scope of DdaG and SfaB suggested the potential of using amide synthetases to produce novel value-added oligoamides.

We also determined that DdaG regioselectively ligated **S** with the asymmetric polyamide spermidine. Of the two possible primary amines in spermidine, DdaG selectively coupled **S** with the primary amine nearer the middle amine, producing only a single diad product. This regioselectivity highlights the potential of amide synthetases to synthesize ordered polymers from asymmetric substrates, which is challenging for traditional chemical synthesis methods (Figure S91)^47^.

Given the distinct activities of these enzymes with diacids and their considerable tolerance for various diamines, we investigated the effect of diacid and diamine chain length on enzyme activity. We tested linear diacids and diamines with carbon lengths of 7 to 10, excluding C10 diamines due to solubility issues (Figure S92). The results for DdaG were consistent with our previous findings, showing a preference for **S** as the diacid and comparable activity across the full range of diamine lengths. In contrast, SfaB, although less active in ligating **S** with longer diamines, showed activity with C7-C10 diacids and **M**. We hypothesize that SfaB has a larger substrate-binding pocket to accommodate longer diacids.

To further characterize DdaG performance, we conducted kinetic studies using succinic acid and hexamethylenediamine as substrates. The *k*_cat_ and *K*_m_ values for DdaG with these substrates were determined to be 3.2 ± 0.2 min^−1^ and 8.8 ± 1.8 mM (Figure S93), respectively. Compared to its native reaction with fumaric acid and 2,3-diaminopropionic acid,^35^ the *k*_cat_ with non-native substrates is 4-fold lower and *K*_m_ is 15-fold higher.

Finally, we tested amide synthetase activity using cell-free enzyme expression. When expressed in a cell-free system, DdaG showed high activity ligating its native substrates but low production of the PA64 diad (Figure S94). Similarly, SfaB exhibited higher activity with a short chain fatty acid (i.e., 5-chlorovalerate) than glutarate. These results suggest that amide synthetases have substantial engineering potential to enhance their activity with non-native or non-preferred substrates, and a cell-free system could serve as an efficient platform for high-throughput enzyme engineering.^31^

## Conclusion

In this work, we demonstrated that nylon diads can be polymerized to yield polymers with higher molecular weight than the corresponding traditional salts, particularly for Nylon 65 incorporating the bioderived monomer glutaric acid. Soluble amide synthetases provide a simple route to regioselectively couple unprotected, nylon-relevant diacids, diamines, and ω-amino acids into amide diads. Reaction molar yields reached up to 80%, while reaction kinetics demonstrated substantial room for further improvement through enzyme engineering. Reaction scale-up and optimization highlighted the potential for simple and efficient enzymatic synthesis of sufficient quantities of oligoamides for further characterization. Moreover, the broad substrate scope of these amide synthetases allows for the facile synthesis of diverse novel nylon oligomers, thereby facilitating the development of next-generation polyamides with enhanced or specialized properties. Notably, this approach could be extended from *in vitro* to *in vivo* systems by integrating enzyme-catalyzed amide bond formation with microbial monomer synthesis via engineered metabolic pathways. By reducing the need for pH control and product separation, simultaneous biosynthesis and ligation would offer substantial process advantages for nylon synthesis from renewable feedstocks.

## Methods

### Materials

LB agar, LB medium and Terrific Broth were purchased from BD Difco. All chemicals were purchased from ThermoFisher, Alfa Aesar, or Tokyo Chemical Industry (TCI). Pyrophosphatase was purchased from New England Biolabs.

### Synthesis and expression of enzymes

Genes encoding DdaG^35^ (GenBank: ADN39487.1), SfaB^48^ (GenBank: ATY72527.1), McbA^34^ (GenBank: AGL76720.1), and PylC^43^ (GenBank: AAZ69813.1) were synthesized by GenScript (Piscataway, NJ) with a N-terminal His_6_ tag for purification and cloned into pET28a(+). The gene encoding SuCphA1^42^ (GenBank: BAA17890.1) was synthesized by GenScript (Piscataway, NJ) with a N-terminal His6 tag for purification and cloned into pETDuet-1. For the His-SUMO-PylC construct, PCR amplified PylC was inserted into Plasmid #112793 purchased from Addgene and replaced the location of EfaCas1 using Gibson Assembly. Plasmids containing His-DdaG, His-SfaB, His-McbA, His-PylC, His-SUMO-PylC, His-SuCphA1, His-PPK12 were transformed into chemically competent BL21 (DE3) *Escherichia coli* cells (Sigma-Aldrich, St Louis, MO) according to the manufacturer’s specifications.

His-PPK12 was produced as described previously.^44^ Expression and purification of His-DdaG, His-SfaB, His-McbA, His-PylC, His-SUMO-PylC and His-SuCphA1 were conducted following previously published methods. A colony or glycerol stock of BL21(DE3) *E. coli* containing plasmid DNA was used to inoculate 10 mL of LB medium supplemented with appropriate antibiotics. The culture was incubated at 37 °C and 250 rpm overnight. The overnight culture was then diluted 100-fold into 200 mL of Terrific Broth. The subculture was incubated at 37 °C at 250 rpm until an OD of 0.4-0.6 was reached. Protein expression was induced by adding IPTG to a final concentration of 0.1 mM. The temperature was decreased to 16 °C and the culture was grown with shaking overnight. The cells were harvested by centrifugation (4000 rpm, 40 min, 4 °C), resuspended in 5 mL of lysis buffer (50 mM HEPES, 300 mM NaCl, 10 mM imidazole, 10mM MgCl_2_, 10% glycerol, pH 8.0), and sonicated on ice. Cell debris was removed by centrifugation (10,000 rpm, 30 min, 4 °C). The supernatant was then filtered with an MCE filter (0.22 μm) and loaded at 3 mL/min onto an ÄKTA Start FPLC (Cytiva Marlborough, MA) equipped with a 5-mL His-Trap^TM^ column (Cytiva). The column was washed with wash buffer (50 mM HEPES, 300 mM NaCl, 30 mM imidazole, 10 mM MgCl_2_, 10% glycerol, pH 8.0) before eluting with elution buffer (50 mM HEPES, 300 mM NaCl, 250 mM imidazole, 10 mM MgCl_2_, 10% glycerol, pH 8.0) at 3 mL/min. Purified proteins were concentrated with an Amicon® Ultra centrifugal filter at 10 kDa cutoff, and buffer exchanged into exchange buffer (50 mM HEPES, pH 8.0, 100 mM NaCl, 10mM MgCl_2_, 10% glycerol). The final purified product was analyzed by SDS-PAGE and enzyme concentration was measured using a NanoDrop™ 1000 Spectrophotometer (Thermo Scientific) with exchange buffer as a blank, and the extinction coefficient of each protein was calculated using ProtParam from the ExPASy Proteomics Server. The molecular weight of each protein was calculated using ProtParam from the ExPASy Proteomics Server, and further confirmed by the SDS-PAGE method. Pure enzymes were stored at −80°C for subsequent activity assays.

### In vitro enzyme activity assays

A typical enzymatic reaction was carried in triplicate at 50 μL containing 100 mM HEPES (pH 8.0), 10 mM ATP, 10 mM MgCl_2_, 10 μM enzyme, 5 mM carboxyl group-containing compounds and 5 mM amine group-containing compounds, and was incubated at 30 °C for 16 h with shaking at 200 rpm. The reaction was quenched by adding 50 μL methanol. The samples were centrifuged at 4,000 rpm for 10 min, and the supernatant was subjected to MS analysis to determine the formation of corresponding products. For the 50 ml reaction, 100 mM HEPES (pH 8.0), 50 mM ATP, 50 mM MgCl_2_, 10 μM enzyme, 50 mM disodium succinic acid and 50 mM hexamethylenediamine were incubated at 30 °C for 16 h with shaking at 200 rpm. The reaction was quenched by adding 50 mL methanol. The samples were centrifuged at 4,000 rpm for 20 min, and the supernatant was subjected for product isolation.

### Enzymatic reaction optimization coupling with ATP regeneration system

Reactions with ATP regeneration were performed in 100 mM HEPES pH 8, 1 mM ATP, 1 or 5 mM AMP, 10-100 mM MgCl_2_, 5-25 mM SHMP, 50 mM disodium succinic acid, 50-250 mM hexamethylene diamine,10 μM DdaG, 0.2 U ecPPase, and 1 mg/mL PPK12. These reactions were then incubated at 30 ºC with shaking at 200 rpm. After 24h, reaction mix was quenched with 1 volume of methanol and samples were collected after centrifugation to clear insoluble precipitates.

### Kinetic analysis of DdaG on reaction of succinate and hexamethylenediamine

The ligation of succinate and hexamethylenediamine by purified DdaG was monitored by observation of the product MS formation according to OPSI-MS. Kinetic assays were performed in triplicate with a reaction mixture containing 0.5 μM DdaG, 10 mM ATP, 100 mM HEPES (pH 8), 10 mM MgCl_2_, 0.5 mM to 50 mM disodium succinate, 50 mM hexamethylenediamine in total volume of 50 μl reaction. Samples were taken after incubation at 30 °C for 5 min, 15 min, 25 min, and 35 min. The reaction was quenched by adding 50 μL methanol. The samples were centrifuged at 4,000 rpm for 10 min, and the supernatant was subjected to OPSI-MS analysis to determine the formation of corresponding products. Initial velocities were determined by linear regression analysis of the data. The kinetic parameters *k*_cat_, and *K*_m_ and standard error were calculated by Graphpad Prism 9.

### I.DOT/OPSI-MS Analysis of Enzyme Activity

The immediate drop-on-demand technology (Dispendix GmbH, Stuttgart, Germany) coupled with open port sampling interface-mass spectrometry (I.DOT/OPSI-MS) was used to analyze enzymatic reactions as previously described in detail.^46^ Briefly, single droplets were mixed with solvent in an on-line fashion, enabling sampling throughputs of up to 2 s/sample. Here, enzymatic reactions were diluted 1:100 (*v/v*) in high performance liquid chromatography (HPLC) grade water with 500 nM propranolol acting as an internal standard for droplet capture. 40 μL of diluted reactions were transferred to a I.DOT S.100 96-well plate and analyzed without further separations. The I.DOT system was used to eject 20 nL of sample into the OPSI, into a flow of 75/25/0.1 (*v/v/v*) acetonitrile/water/formic acid and transported the sample to the electrospray ion source of the mass spectrometer. Dispensing throughput was 4 s/sample and each sample was measured in triplicate. Peak widths were ∼1.2 s wide.

For identification of products by high mass resolution, a Thermo Q-Exactive HF mass spectrometer (ThermoFisher Scientific) in positive ion mode was used for characterization. Scan settings were sheath gas = 80, auxiliary gas = 40, electrospray voltage = 4 kV, ion injection time = 50 ms, automatic gain control = 3e^6^, capillary temperature = 200 °C, mass/charge (*m/z*) range = 100-750 *m/z*, and OPSI solvent flow of 250 μL/min. A 60,000 mass resolution was used for quantitative analyses while 240,000 resolution was used for compound identification. Tandem MS data were acquired using the Q-Exactive HF, but water adduct formation was commonly observed in the ion trap. To avoid water adducts in tandem mass spectra, tandem mass spectrometry data was collected using a Sciex 5600 TripleTOF time of flight mass spectrometer (Sciex) with the following settings: GS1 = 75, GS2 = 35, electrospray voltage = 5.5 kV, capillary temperature = 200 °C, and OPSI solvent flow of 150 μL/min. Collision energies across 10-50 eV were evaluated for each ion.

Control of I.DOT settings (positioning, well selection, timing and droplet dispensing cycles), OPSI peak finding, data extraction from vendor file formats, and signal integration were enabled through custom software packages written in Delphi, Python, and C#, available upon request. For each droplet sampling event, the average mass spectrum from the resulting mass spectral peak was normalized to propranolol internal standard signal and background subtracted. Log-log matrix matched calibration curves were used for quantitation. Mass spectrometric signals typically spanned several orders of magnitude, thus, log-log calibrations resulted in reduced error across the full concentration range.

### Tandem mass spectra

I.DOT/OPSI-MS^2^ data were acquired for the possible observed compounds shown in Figures 2 and 3 in the main text. MS^2^ data were acquired on a Sciex 5600 TripleTOF+ instrument using the same OPSI-MS probe as described previously. A time-of-flight system was preferred as in trap water adducts were observed for some compounds when fragmented with the orbitrap system. Putative fragment identifications are given for each compound using hypothesized fragmented pathways. All tandem mass spectra shown used a collision energy of 20eV and a declustering potential of 100. Additional collision energies were used to help with fragment identifications as necessary (data not shown). For brevity, a single tandem mass spectrum is shown for each possible compound. For compounds without a tandem mass spectrum, signals for those compounds were either too weak to generate clear tandem MS or were obfuscated by overlapping background ions in the 1 *m/z* isolation window.

### Nuclear magnetic resonance (NMR) spectroscopy

^1^H and ^13^C nuclear magnetic resonance NMR spectroscopy was performed on a Bruker Avance III 400 NMR spectrometer operating at 400 MHz (^1^H) or 100 MHz (^13^C). Chemical shifts are reported in parts per million (ppm) relative to residual protonated solvent.

### Size exclusion chromatography (SEC)

SEC analysis was performed on an Agilent 1260 Infinity II LC system equipped with an Agilent PL HFIPgel guard column (9 μm, 50 × 4.6 mm) and two Agilent PL HFIPgel columns (9 μm, 250 × 4.6 mm). A 0.02 M CF_3_COONa HFIP solution was used as the mobile phase at 40 °C and a flow rate of 0.300 mL min^-1^. Elution time was monitored using a differential refractive index (dRI) detector, a light scattering detector operating with two angles at 90° and 15°, and a differential viscometer. Number-average molecular weights (*M*_n_), weight-average molecular weights (*M*_w_) and dispersities (Ð = *M*_w_/*M*_n_) were calculated based on calibrations with PMMA standards using the Agilent GPC/SEC software.

### Thermogravimetric analysis (TGA)

TGA was performed on a TA Instruments Discovery TGA 55 under a nitrogen atmosphere. Sample mass was recorded as a function of temperature while the sample was heated from room temperature to 600 °C at a heating rate of 20 °C min^-1^. Degradation temperature (*T*_d_) was taken as the temperature at which the sample had lost 5% of its initial mass.

Isothermal TGA studies were performed on a TA Instruments Discovery TGA 55 using a platinum pan under a nitrogen atmosphere. Samples were initially conditioned at 100 °C for 1 h to remove any residual moisture or solvent, and subsequently heated to 220 °C for 8 h to simulate SSP conditions. Theoretical mass loss (*ΔΔm*_th_) was calculated from the following equation:

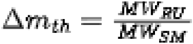

where *MW*_RU_ is the molecular weight of the polymer repeat unit, and *MW*_SM_ is the molecular weight of the salt or diad starting material. Excess mass loss was then calculated as the difference between the experimentally measured mass loss (*ΔΔm*) and *ΔΔm*_th_.

### Differential scanning calorimetry (DSC)

DSC was performed on a TA Instruments Discovery DSC 250. Measurements were carried out in standard aluminum pans using a heat-cool-heat cycle with 10 and 5 °C heating and cooling rates, respectively. Melting (*T*_m_) temperature and enthalpy of melting were measured from the first heat cycle while glass transition (*T*_g_) temperatures were measured from the second heating cycle.

### Fourier-transform Infrared (FTIR) Spectroscopy

Attenuated total reflectance (ATR) Fourier-transform infrared spectroscopy was performed using a PerkinElmer Spectrum Two FTIR spectrometer. Transmission spectra were collected from 4000 to 500 cm^-1^.

## Acknowledgements

Research was sponsored by the Laboratory Directed Research and Development Program of Oak Ridge National Laboratory, managed by UT-Battelle, LLC under Contract No. DE-AC05-00OR22725, for the U. S. Department of Energy. Work by N.T.Z was supported by the U.S. Department of Energy, Office of Science, Office of Workforce Development for Teachers and Scientists (WDTS) under the Science Undergraduate Laboratory Internships (SULI) program. We thank Madan Gopal and Wilfred Chen (University of Delaware) for the PPK12 expression plasmid. LQ, ITD, JCF, and JKM are inventors on a patent application related to this work.

## Author contributions

L.Q. performed the majority of the enzymatic characterization and optimization. I.T.D.: oligomer synthesis, polymerization, and characterization. D.L.C: Analytical characterization. V.K.: Analytical method development. N.T.Z.: Characterization of long-chain substrates. N.D.O.: Polymer thermal characterization. N.A.T.: Oligomer synthesis. J.F.C.: Analytical methods development and analysis, supervision of analytical research. J.C.F.: Conceptualization, supervision of chemistry research. J.K.M: Conceptualization; funding acquisition; supervision of biochemistry research.

