## Supplementary Information for "Improved nylon polymerization using amide diads"

**Supplemental Materials and Methods**

All solvents and reagents were used as received unless otherwise stated. *N*-Boc-1,6-hexanediamine, *N*-boc-cadaverine, monomethyl adipate, glutaric anhydride, and 1-Ethyl-3-(3′-dimethylaminopropyl)carbodiimide hydrochloride (EDC·HCl) were purchased from Ambeed. Absolute ethanol (EtOH), trifluoroacetic acid, and Reillex 402 were purchased from Sigma Aldrich. Acetic anhydride ($Ac_2O$), dichloromethane (DCM), methanol (MeOH), and diethyl ether ($Et_2O$) were purchased from VWR. Lithium hydroxide was purchased from Thermo Scientific. Succinic anhydride was purchased from Beantown Chemical. 1,1,1,3,3,3-hexafluoroisopropanol (HFIP) was purchased from Apollo Scientific.

**Procedure A:** *General procedure for ring-opening reaction of cyclic anhydride*: In a 500 mL RBF, mono-boc-protected diamine (1.1 equiv) was added to a solution of cyclic anhydride (1 equiv) dissolved in DCM (0.1 M). The reaction was stirred at r.t. overnight. The resulting precipitate was filtered, rinsed with additional DCM, and dried under vacuum.

**Procedure B:** *General procedure for removal of Boc protecting groups*: In a 500 mL RBF, Boc-protected amine was dissolved in DCM (15 mL $g^{-1}$). The RBF was chilled to 0 °C in an ice bath and trifluoroacetic acid (5 mL $g^{-1}$) was added slowly to the reaction. The flask was stirred overnight, warming gradually to room temperature. The majority of trifluoroacetic acid was removed by concentrating the reaction *in vacuo* and diluting with DCM for 3 cycles. After the third cycle, the concentrated crude was precipitated into diethyl ether. The ether layer was decanted and the product was dried under vacuum.

**Procedure C:** *General procedure for freebasing TFA salt*: Reillex 402 (5 equiv.) was added to a stirred solution of TFA salt (1 equiv.) dissolved in deionized water (10 mL $g^{-1}$) and stirred overnight. The reaction was filtered and the filtrate was concentrated *in vacuo* to provide a slightly yellow residue that was further purified by triturating with ethanol.

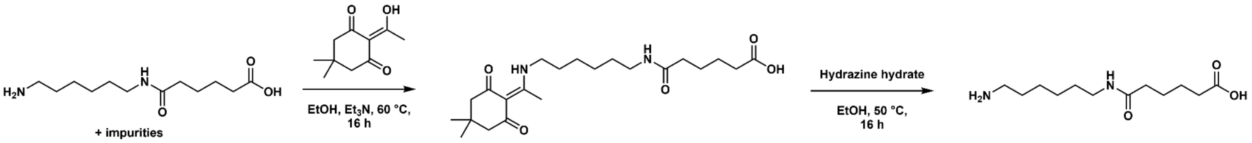


**Scheme S1.** Purification protocol for isolation of enzymatically synthesized MA.

*Isolation of dde-functionalized MS*. The crude enzymatic reaction mixture was transferred to a 100 mL rbf and dried *in vacuo* to provide a white solid that was resuspended in EtOH (25 mL). To this suspension, 2-acetyldimedone (660 mg, 1.5 equiv relative to initial concentration of hexamethylene diamine) and $Et_3N$ (0.5 mL) were added. The flask was equipped with a reflux condenser and the mixture was heated to 50 °C overnight. The reaction mixture was concentrated *in vacuo*, redissolved in $H_2O$ (25 mL), and acidified to *ca*. pH 4 with a 1 M HCl solution. The aqueous solution was extracted 5X with DCM (25 mL). The combined organic extracts were washed with brine (25 mL), dried over sodium sulfate, and concentrated *in vacuo*. The resulting yellow residue was further purified by flash chromatography using a stepwise gradient (0-10% MeOH in DCM) to provide the product as a yellow oil (118.6 mg; XX%). $^1H$ NMR (400 MHz, $CDCl_3$) δ 13.24 (s, 1H), 6.32 (s, 1H), 3.34 (q, $J$ = 6.3 Hz, 2H), 3.18 (q, $J$ = 6.4 Hz, 2H), 2.62 (t, $J$

= 6.5 Hz, 2H), 2.48 (s, 3H), 2.41 (t, $J$ = 6.4 Hz, 2H), 2.30 (s, 4H), 1.63 (t, $J$ = 7.1 Hz, 2H), 1.46 (t, $J$ = 7.0 Hz, 2H), 1.41 – 1.23 (m, 4H), 0.96 (s, 6H). $^{13}$C NMR (101 MHz, $CDCl_3$) δ 198.03, 175.29, 173.91, 172.56, 107.89, 52.65, 43.34, 39.35, 31.16, 30.16, 29.17, 28.64, 28.25, 26.37, 26.20, 18.13.

*Deprotection of dde-functionalized MS*. Hydrazine hydrate (15.1 µL, 0.31 mmol, 1 equiv) was added to a solution of dde-MS (118.6 mg, 0.31 mmol, 1 equiv) in EtOH (2 mL) in a scintillation vial. The reaction was heated to 50 °C overnight to form a white precipitate. The contents of the reaction were transferred to a centrifuge tube along with an additional 2 mL EtOH, and the reaction was centrifuged. The supernatant was discarded. The resulting white powder was rinsed with an additional 2 mL EtOH and dried under vacuum to provide MS as a white powder (58.0 mg, 86.4%). Analytical data was found to be identical to chemically synthesized MS.

*Isolation of dde-functionalized MA*. $^{1}$H NMR (400 MHz, $CDCl_3$) δ 13.31 (s, 1H), 6.24 (t, $J$ = 5.6 Hz, 1H), 3.41 (q, $J$ = 6.3 Hz, 2H), 3.27 (q, $J$ = 6.3 Hz, 2H), 2.56 (s, 3H), 2.37 (s, 6H), 2.29 – 2.21 (m, 2H), 1.83 – 1.62 (m, 6H), 1.56 (p, $J$ = 6.9 Hz, 2H), 1.44 (dddt, $J$ = 18.8, 15.3, 9.3, 4.8 Hz, 4H), 1.04 (s, 6H). $^{13}$C NMR (101 MHz, $CDCl_3$) δ 198.08, 176.55, 173.89, 173.00, 107.89, 52.64, 43.56, 39.27, 36.10, 33.36, 30.17, 29.06, 28.72, 28.25, 26.52, 26.43, 24.84, 24.08, 18.13.

*Deprotection of dde-functionalized MA.* Dde-functionalized MA was deprotected using analogous conditions as dde-MS. MA was collected as a white powder (10.3 mg, 78.9%). Analytical data was found to be identical to chemically synthesized MA.

*Preparation of nylon salts*: A 50 wt% solution of diamine dissolved in water was added to a 10 wt% solution of dicarboxylic acid dissolved in ethanol (adipic or glutaric acid) or methanol (succinic acid) while heated to 50 °C. The reaction was stirred for an additional two hours and subsequently cooled to room temperature. The resulting white precipitate was filtered, rinsed with additional alcohol, and dried at room temperature under vacuum. In some cases, it was necessary to chill the reaction to -20 °C overnight for the salt to precipitate.

*Solid state polymerization:* Diad or nylon salt (1 g) was weighed into a 50 mL RBF. The flask was sealed with a rubber septum and purged with Ar gas for 15 min. The RBF was heated to 220 °C for 1 h under a continuous flow of Ar. The flask was then cooled to RT and an aliquot of the prepolymer was collected for SEC analysis. The flask was then equipped with a flow control adapter and the prepolymer was further polymerized at 220 °C under vacuum.

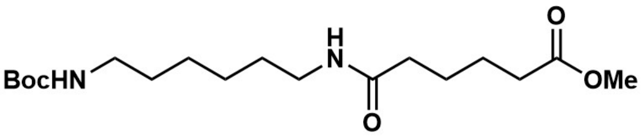


*Boc-MA-OMe.* A 1000 mL rbf was charged with monomethyl adipate (7.5 g, 46.8 mmol, 1 equiv.) and 450 mL DCM. EDC·HCl (9.87 g, 51.5 mmol, 1.1 equiv.) was added and the RBF was stirred at r.t. for 5 min until dissolved. *N*-Boc-1,6-hexanediamine (11.15 g, 51.5 mmol, 1.1 equiv) was then added to the reaction mixture and the reaction was stirred at r.t. overnight. The reaction was washed with 2×250 mL 1 M HCl, 3×250 mL mL saturated $NaHCO_3$ solution, and 250 mL brine. The organic layer was dried over sodium sulfate, filtered, and concentrated using a rotary evaporator. The product (11.02 g, 66%) was dried overnight in a vacuum oven and used in subsequent reactions without further purification. $^{1}$H NMR (400 MHz, $CDCl_3$) δ 5.71 (s, 1H), 4.55 (s, 1H), 3.66 (s, 3H), 3.22 (td, *J* = 7.0, 5.8 Hz, 2H), 3.10 (q, *J* = 6.7 Hz, 2H), 2.39 – 2.25 (m, 2H), 2.18 (dq, *J* = 6.7, 3.5 Hz, 2H), 1.84 – 1.57 (m, 5H), 1.43 (s, 13H), 1.32 (p, *J* = 3.9 Hz, 4H). $^{13}$C NMR (101 MHz, $CDCl_3$) δ 174.14, 172.61, 156.23, 79.18, 51.70, 40.27, 39.22, 36.44, 33.81, 30.11, 29.56, 28.54, 26.30, 26.15, 25.26, 24.54.

*Boc-MA.* To a solution of Boc-MA-OMe (11 g, 30.6 mmol) dissolved in THF (110 mL) was added 1 M LiOH (110 mL). The reaction was stirred for 3.5 h, and subsequently acidified to pH 4 with 1 M HCl. THF was removed via rotary evaporation, and the white precipitate was filtered to provide the carboxylic acid in quantitative yield. $^{1}$H NMR (400 MHz, DMSO) δ 12.01 (s, 1H), 7.75 (t, *J* = 5.7 Hz, 1H), 6.77 (t, *J* = 5.7 Hz, 1H), 3.00 (q, *J* = 6.6 Hz, 2H), 2.88 (q, *J* = 6.6 Hz, 2H), 2.19 (t, *J* = 6.9 Hz, 2H), 2.04 (t, *J* = 6.8 Hz, 2H), 1.57 – 1.41 (m, 4H), 1.37 (s, 13H), 1.23 (dt, *J* = 7.7, 3.8 Hz, 4H). $^{13}$C NMR (101 MHz, DMSO) δ 174.92, 172.12, 156.04, 77.75, 40.16, 38.78, 35.58, 33.96, 29.91, 29.61, 28.74, 26.59, 26.47, 25.35, 24.63.

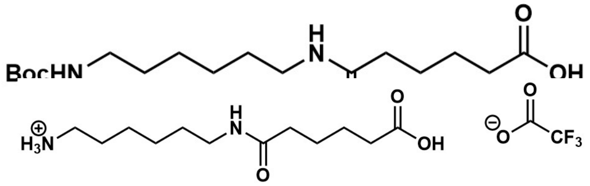

*MA TFA salt.* Product was prepared according to Procedure B and isolated as a viscous yellow oil (8.46 g, 81%). $^{1}$H NMR (400 MHz, $D_2O$) δ 3.14 (t, *J* = 6.8 Hz, 2H), 2.94 (t, *J* = 7.6 Hz, 2H), 2.44 – 2.30 (m, 2H), 2.21 (td, *J* = 6.7, 4.1 Hz, 2H), 1.70 – 1.21 (m, 12H). $^{13}$C NMR (101 MHz, MeOD) δ 177.28, 175.81, 163.18, 119.66, 40.60, 40.01, 36.71, 34.56, 30.14, 28.43, 27.30, 26.95, 26.50, 25.54.

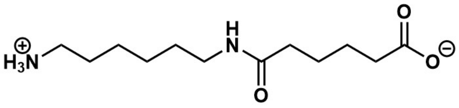

*MA*. Product was prepared according to Procedure C and isolated as a white powder (3.28 g, 60%). $^{1}$H NMR (400 MHz, $D_2O$) δ 3.05 (t, *J* = 6.7 Hz, 2H), 2.84 (t, *J* = 7.6 Hz, 2H), 2.11 (t, *J* = 6.8 Hz, 2H), 2.05 (t, *J* = 7.0 Hz, 2H), 1.67 – 1.31 (m, 8H), 1.31 – 1.09 (m, *J* = 5.5, 4.1 Hz, 4H). $^{13}$C NMR (101 MHz, $D_2O$) δ 183.28, 176.64, 39.30, 38.91, 37.11, 35.54, 27.91, 26.54, 25.34, 25.33, 25.16, 25.09.

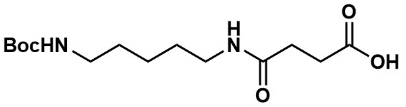

*Boc-CS*. Product was prepared according to Procedure A using N-boc-cadaverine (5 g, 24.7 mmol, 1.1 equiv.) and succinic anhydride (2.25 g, 22.5 mmol, 1 equiv.). The product was isolated as a white powder (4.24 g, 62%). $^{1}$H NMR (400 MHz, DMSO) δ 12.05 (s, 1H), 7.79 (t, *J* = 5.6 Hz, 1H), 6.75 (t, *J* = 5.7 Hz, 1H), 2.99 (q, *J* = 6.6 Hz, 2H), 2.87 (q, *J* = 6.6 Hz, 2H), 2.40 (t, *J* = 7.2 Hz, 2H), 2.28 (t, *J* = 6.9 Hz, 2H), 1.36 (s, 14H), 1.21 (qd, *J* = 7.0, 2.4 Hz, 2H). $^{13}$C NMR (101 MHz, DMSO) δ 173.95, 170.77, 155.64, 77.37, 39.82, 38.52, 30.05, 29.24, 29.23, 28.87, 28.32, 23.73.

*CS TFA salt*. Product was prepared according to Procedure B and collected as a viscous yellow oil (3.46 g, 82%). $^{1}$H NMR (400 MHz, MeOD) δ 3.19 (t, *J* = 6.9 Hz, 2H), 3.03 – 2.82 (m, 2H), 2.59 (t, *J* = 6.6 Hz, 2H), 2.46 (t, *J* = 6.6 Hz, 2H), 1.82 – 1.61 (m, 2H), 1.61 – 1.48 (m, 2H), 1.48 – 1.24

(m, 2H). $^{13}$C NMR (101 MHz, MeOD) δ 176.30, 174.58, 163.18, 49.85, 40.58, 39.83, 31.49, 30.25, 29.81, 28.07, 24.53.

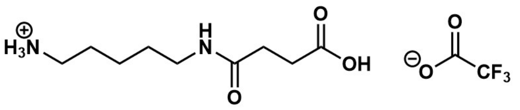

*CS*. Product was prepared according to Procedure C and isolated as a white powder (1.92 g, 80%). $^{1}$H NMR (400 MHz, $D_2O$) δ 3.16 (t, $J$ = 6.7 Hz, 2H), 2.95 (t, $J$ = 7.5 Hz, 2H), 2.41 (d, $J$ = 2.1 Hz, 4H), 1.63 (p, $J$ = 7.6 Hz, 2H), 1.50 (p, $J$ = 6.9 Hz, 2H), 1.35 (qd, $J$ = 9.0, 5.9 Hz, 2H). $^{13}$C NMR (101 MHz, $D_2O$) δ 180.87, 175.71, 39.25, 38.76, 33.06, 32.38, 27.67, 26.23, 22.74.

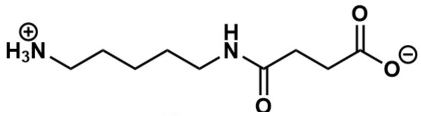

*Boc-MG*. Product was prepared according to Procedure A using N-boc-hexanediamine (10 g, 46.2 mmol, 1.1 equiv) and glutaric anhydride (4.80 g, 42.0 mmol, 1 equiv.). The product was isolated as a white powder (7.71 g, 55%). $^{1}$H NMR (400 MHz, DMSO) δ 7.76 (t, $J$ = 5.6 Hz, 1H), 6.77 (t, $J$ = 5.7 Hz, 1H), 3.00 (q, $J$ = 6.5 Hz, 2H), 2.88 (q, $J$ = 6.6 Hz, 2H), 2.18 (t, $J$ = 7.4 Hz, 2H), 2.06 (t, $J$ = 7.4 Hz, 2H), 1.69 (p, $J$ = 7.4 Hz, 2H), 1.36 (s, 14H), 1.21 (p, $J$ = 3.6 Hz, 4H). $^{13}$C NMR (101 MHz, DMSO) δ 174.26, 171.40, 155.63, 77.34, 39.80, 38.38, 34.50, 33.08, 29.50, 29.17, 28.32, 26.17, 26.05, 20.79.

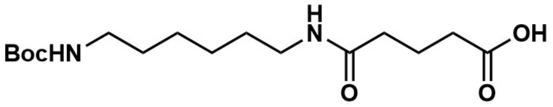

*MG TFA salt*. Product was prepared according to Procedure B and collected as a white solid (6.98 g, 96%). $^{1}$H NMR (400 MHz, DMSO) δ 7.80 (dd, $J$ = 13.8, 8.0 Hz, 4H), 3.01 (q, $J$ = 6.5 Hz, 2H), 2.76 (h, $J$ = 6.0 Hz, 2H), 2.18 (t, $J$ = 7.4 Hz, 2H), 2.07 (t, $J$ = 7.4 Hz, 2H), 1.69 (p, $J$ = 7.4 Hz, 2H), 1.51 (p, $J$ = 7.3 Hz, 2H), 1.43 – 1.12 (m, 6H). $^{13}$C NMR (101 MHz, DMSO) δ 174.66, 171.93, 158.97, 158.64, 118.57, 115.63, 40.51, 40.31, 40.10, 39.89, 39.68, 39.47, 39.26, 39.21, 38.69, 34.92, 33.48, 29.40, 27.39, 26.32, 25.92, 21.21.

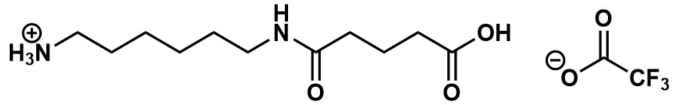

*MG*. Product was prepared according to Procedure C and collected as a white powder (3.62 g, 68%). $^{1}$H NMR (400 MHz, $D_2O$) δ 3.15 (t, $J$ = 6.8 Hz, 2H), 2.95 (t, $J$ = 7.6 Hz, 2H), 2.20 (t, $J$ = 7.5 Hz, 2H), 2.15 (t, $J$ = 7.5 Hz, 2H), 1.79 (p, $J$ = 7.5 Hz, 2H), 1.62 (p, $J$ = 7.3 Hz, 2H), 1.49 (p, $J$ = 6.9 Hz, 2H), 1.42 – 1.20 (m, $J$ = 4.8, 3.8 Hz, 4H). $^{13}$C NMR (101 MHz, $D_2O$) δ 182.22, 176.16, 39.30, 38.97, 36.42, 35.42, 27.92, 26.52, 25.34, 25.08, 22.43.

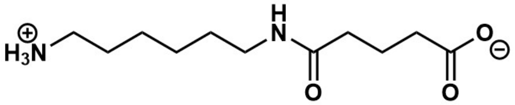

*MS diad*. A modified version of the procedure described by Madhavachary et al. was used [1]. In a 500 RBF, hexamethylenediamine (4 g, 34.4 mmol, 1 equiv.) was dissolved in THF (240 mL). The rbf was equipped with a dropwise addition funnel containing succinic anhydride (3.44 g, 34.4 mmol, 1 equiv.) dissolved in THF (160 mL). The contents of the funnel were added to the RBF dropwise over the course of *ca.* 2 h, and the flask was left to stir at r.t. overnight. The contents of the flask were filtered to provide a white precipitate that was purified twice by flash chromatography (9:1 MeOH:DCM) to provide

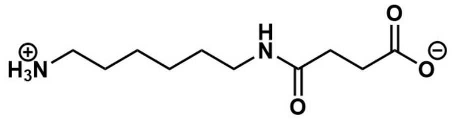

the product as a white powder (3.80 g, 51%). $^{1}$H NMR (400 MHz, $D_2O$) δ 3.14 (t, *J* = 6.7 Hz, 1H), 2.98 – 2.90 (m, 1H), 2.40 (s, 2H), 1.68 – 1.56 (m, 1H), 1.47 (p, *J* = 6.9 Hz, 1H), 1.41 – 1.25 (m, 2H). $^{13}$C NMR (101 MHz, $D_2O$) δ 180.40, 175.50, 39.28, 38.98, 32.67, 32.15, 27.92, 26.50, 25.25, 25.03.

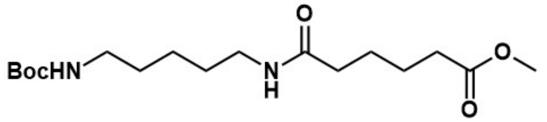


*Boc-CA-OMe.* A 1000 mL rbf was charged with monomethyl adipate (11.11 g, 69.4 mmol, 1 equiv.) and 700 mL DCM. EDC·HCl (14.63 g, 76.3 mmol, 1.1 equiv.) was added and the RBF was stirred at r.t. for 5 min until dissolved. *N*-Boc-1,5-pentanediamine (15.43 g, 76.3 mmol, 1.1 equiv) was then added to the reaction mixture and the reaction was stirred at r.t. overnight. The reaction was washed with 2×250 mL 1 M HCl, 3×250 mL mL saturated $NaHCO_3$ solution, and 250 mL brine. The organic layer was dried over sodium sulfate, filtered, and concentrated using a rotary evaporator. The product (12.54 g,73%) was dried overnight in a vacuum oven and used in subsequent reactions without further purification. $^{1}$H NMR (400 MHz, $CDCl_3$) δ 5.72 (s, 1H), 4.57 (s, 1H), 3.60 (s, 3H), 3.17 (q, J = 6.6 Hz, 2H), 3.04 (q, J = 6.5 Hz, 2H), 2.54 – 2.19 (m, 2H), 2.19 – 2.05 (m, 2H), 1.59 (h, J = 4.6 Hz, 4H), 1.51 – 1.39 (m, 5H), 1.37 (s, 10H), 1.32 – 1.17 (m, 2H). $^{13}$C NMR (101 MHz, $CDCl_3$) δ 174.0, 172.6, 156.1, 79.1, 51.56, 40.2, 39.3, 36.2, 33.7, 29.7, 29.2, 28.4, 25.1, 24.4, 24.4, 23.9.

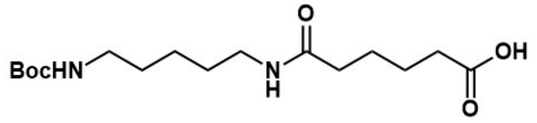


*Boc-CA.* To a solution of Boc-CA-OMe (10 g, 29.0 mmol) dissolved in THF (100 mL) was added 1 M LiOH (100 mL). The reaction was stirred for 3.5 h, and subsequently acidified to pH 4 with 1 M HCl. THF was removed via rotary evaporation, and the white precipitate was filtered to provide the carboxylic acid (7.91 g, 83%). $^{1}$H NMR (400 MHz, DMSO-$d_6$) δ 11.98 (s, 1H), 7.73 (t, J = 5.6 Hz, 1H), 6.75 (t, J = 5.8 Hz, 1H), 3.00 (q, J = 6.5 Hz, 2H), 2.88 (q, J = 6.7 Hz, 2H), 2.20 (t, J = 6.6 Hz, 2H), 2.03 (d, J = 7.0 Hz, 2H), 1.47 (hept, J = 4.8 Hz, 4H), 1.37 (s, 13H), 1.22 (q, J = 8.0 Hz, 2H). $^{13}$C NMR (101 MHz, DMSO-$d_6$) δ 174.85, 172.11, 156.04, 77.77, 38.81, 35.58, 33.87, 29.64, 29.32, 28.74, 25.32, 24.61, 24.17.

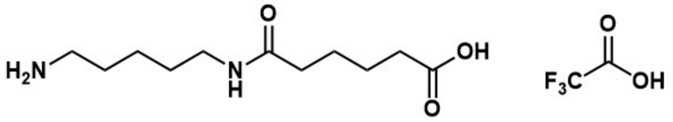


*CA TFA salt.* Product was prepared according to Procedure B and collected as a viscous oil (6.78 g, 93 %).

*CA*. Product was prepared according to Procedure C and collected as a white powder (1.32 g, 40%). $^{1}$H NMR (400 MHz, $D_2O$) δ 3.11 (t, J = 6.7 Hz, 2H), 2.90 (t, J = 7.6 Hz, 2H), 2.15 (dt, J = 14.0, 6.6 Hz, 4H), 1.58 (p, J = 7.6 Hz, 2H), 1.48 (dp, J = 15.1, 7.7 Hz, 5H), 1.38 – 1.11 (m, 2H).

$^{13}C$ NMR (101 MHz, $D_2O$) δ 182.67, 176.73, 39.32, 38.82, 36.60, 35.56, 27.77, 26.33, 25.25, 24.98, 22.93.

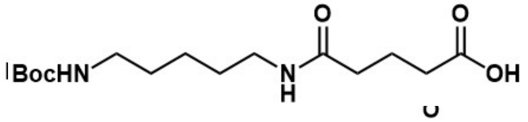

*Boc-CG.* Product was prepared according to Procedure A using N-boc-1,5-pentanediamine (9.75 g, 48.2 mmol, 1.1 equiv) and glutaric anhydride (5 g, 43.8 mmol, 1 equiv.). The product was isolated as a viscous yellow oil (11.72 g, 85%). $^1H$ NMR (400 MHz, DMSO-$d_6$) δ 12.02 (s, 1H), 7.75 (t, J = 5.6 Hz, 1H), 6.75 (t, J = 5.7 Hz, 1H), 3.00 (q, J = 6.6 Hz, 2H), 2.88 (q, J = 6.6 Hz, 2H), 2.19 (t, J = 7.3 Hz, 2H), 2.07 (t, J = 7.4 Hz, 2H), 1.70 (p, J = 7.4 Hz, 2H), 1.37 (s, 13H), 1.22 (q, J = 8.0 Hz, 3H). $^{13}C$ NMR (101 MHz, DMSO-$d_6$) δ 174.64, 171.80, 156.04, 77.77, 38.82, 34.93, 33.51, 29.63, 29.30, 28.74, 24.16, 21.20.

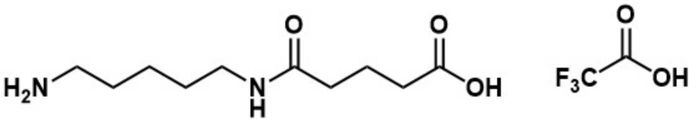

*CG TFA salt.* Product was prepared according to Procedure B using Boc-CG (10 g, 31.6 mmol) and collected as a viscous oil (6.78 g, 93 %). $^1H$ NMR (400 MHz, $D_2O$) δ 3.09 (t, J = 6.8 Hz, 2H), 2.89 (t, J = 7.6 Hz, 2H), 2.51 – 2.25 (m, 2H), 2.18 (q, J = 7.0 Hz, 2H), 1.79 (pd, J = 7.1, 2.0 Hz, 2H), 1.57 (p, J = 7.7 Hz, 2H), 1.44 (p, J = 7.2 Hz, 2H), 1.39 – 1.12 (m, 2H). $^{13}C$ NMR (101 MHz, $D_2O$) δ 177.8, 176.4, 175.8, 117.7, 114.8, 52.13, 39.3, 38.9, 34.8, 34.8, 32.8, 27.7, 26.3, 22.9, 20.7.

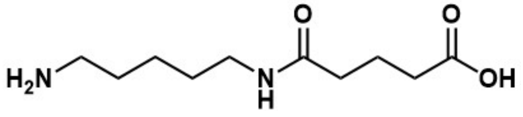

*CG.* Product was prepared according to Procedure C using CG TFA salt (2.5 g, 7.6 mmol) and collected as a white powder (0.8 g, 49%) after crystallizing from ethanol at - 20 °C for ~1 month. $^1H$ NMR (400 MHz, $D_2O$) δ 3.11 (t, J = 6.7 Hz, 2H), 2.91 (t, J = 7.6 Hz, 2H), 2.16 (t, J = 7.5 Hz, 2H), 2.11 (t, J = 7.5 Hz, 2H), 1.74 (p, J = 7.6 Hz, 2H), 1.59 (p, J = 7.6 Hz, 2H), 1.47 (p, J = 7.0 Hz, 2H), 1.31 (td, J = 8.5, 4.1 Hz, 2H). $^{13}C$ NMR (101 MHz, $D_2O$) δ 182.3, 176.3, 39.3, 38.8, 36.5, 35.4, 27.7, 26.3, 22.9, 22.4.

*N-Boc-1,6-hexanediamine*. N-Boc-hexamethylenediamine was prepared according to literature precedent [2]. A 500 mL RBF was charged with 5.81 g HMDA (50 mmol, 5 equiv.), 150 mL DCM, and 2.2 mL $Et_3N$(15 mmol, 1.5 equiv.). The flask was equipped with a dropwise addition funnel containing 2.3 mL $Boc_2O$ (10 mmol, 1 equiv.) dissolved in 50 mL ethyl acetate. The RBF was cooled to 0 °C in an ice bath, and the contents of the addition funnel were added to the reactor dropwise over the course of *ca*. 15 min. The reaction was allowed to stir overnight. The RBF was concentrated in vacuo and redissolved in 150 mL DCM. This crude reaction mixture was washed with 2x150 mL brine and dried over sodium sulfate. The reaction mixture was concentrated *in vacuo* and residue was purified by flash chromatography (20% MeOH in DCM) to provide the product (1.99 g, 92%) as a viscous, colorless liquid. Characterization of this product matched commercial samples.

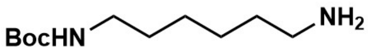

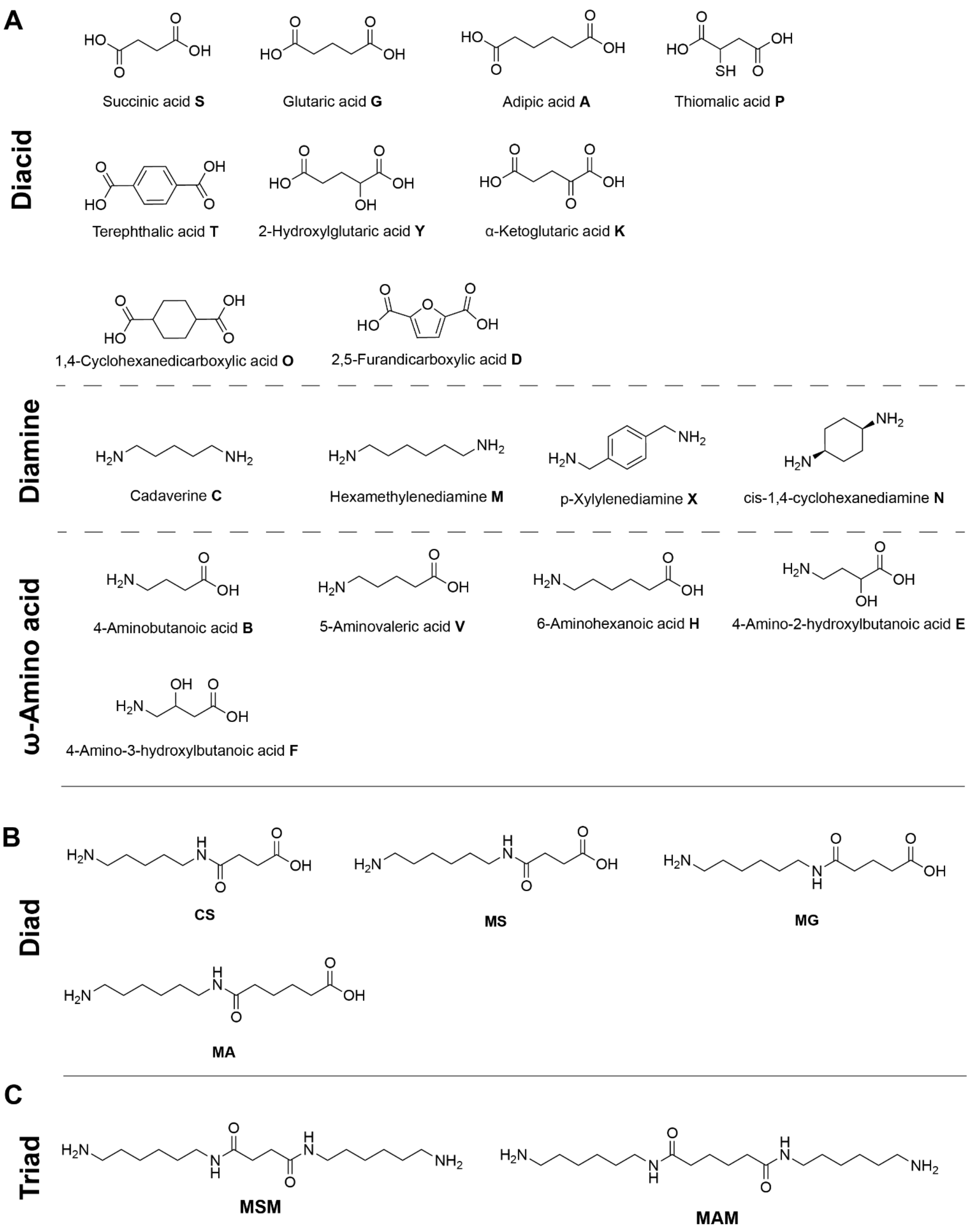


**Figure S1: Structures of substrates in this work**. A) Structures and one-letter codes of tested polymer-relevant monads. B) Structures of diads. C) Structures of triads.

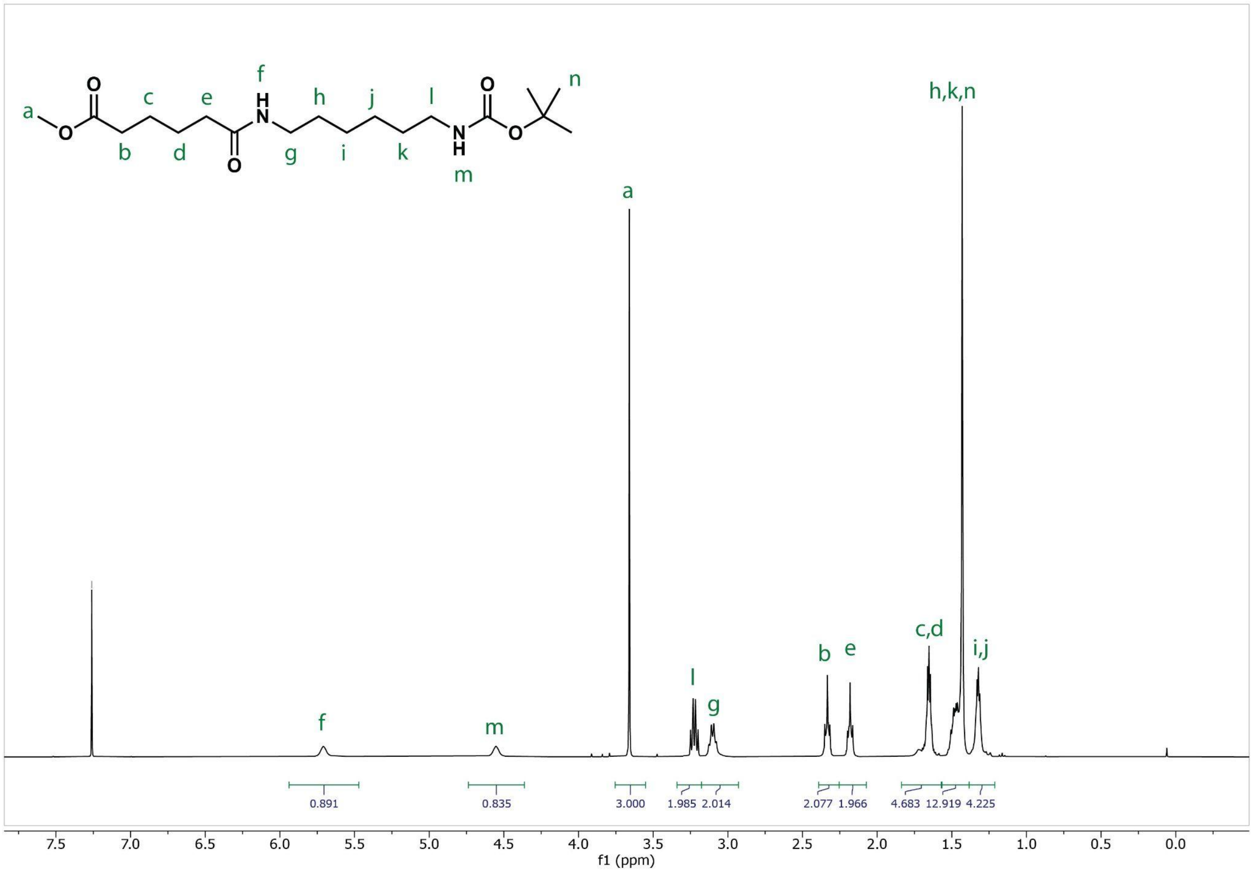


**Figure S2: [1]H NMR spectrum of BocHN-MA-OMe in $CDCl_3$.**

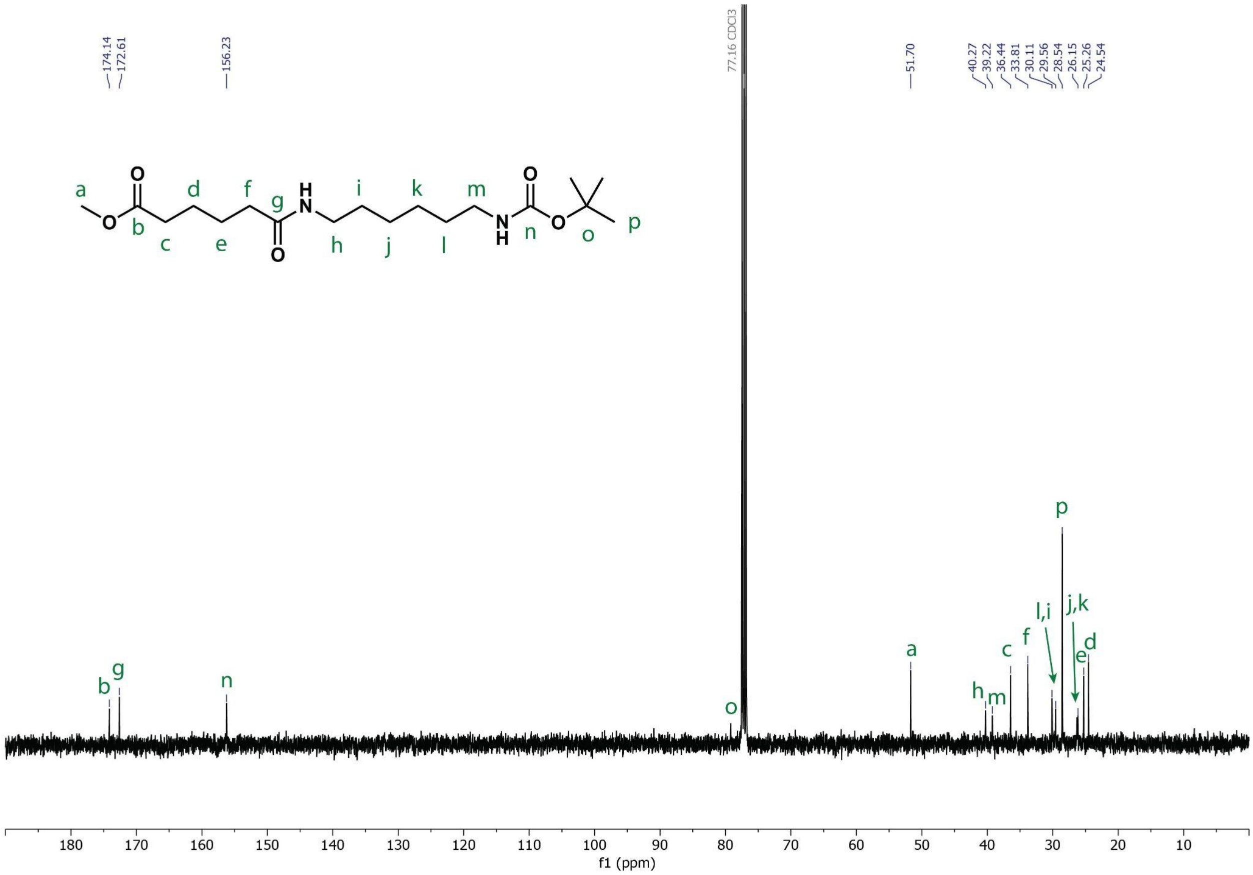


**Figure S3: $^{13}$C NMR spectrum of BocHN-MA-OMe in $CDCl_3$.**

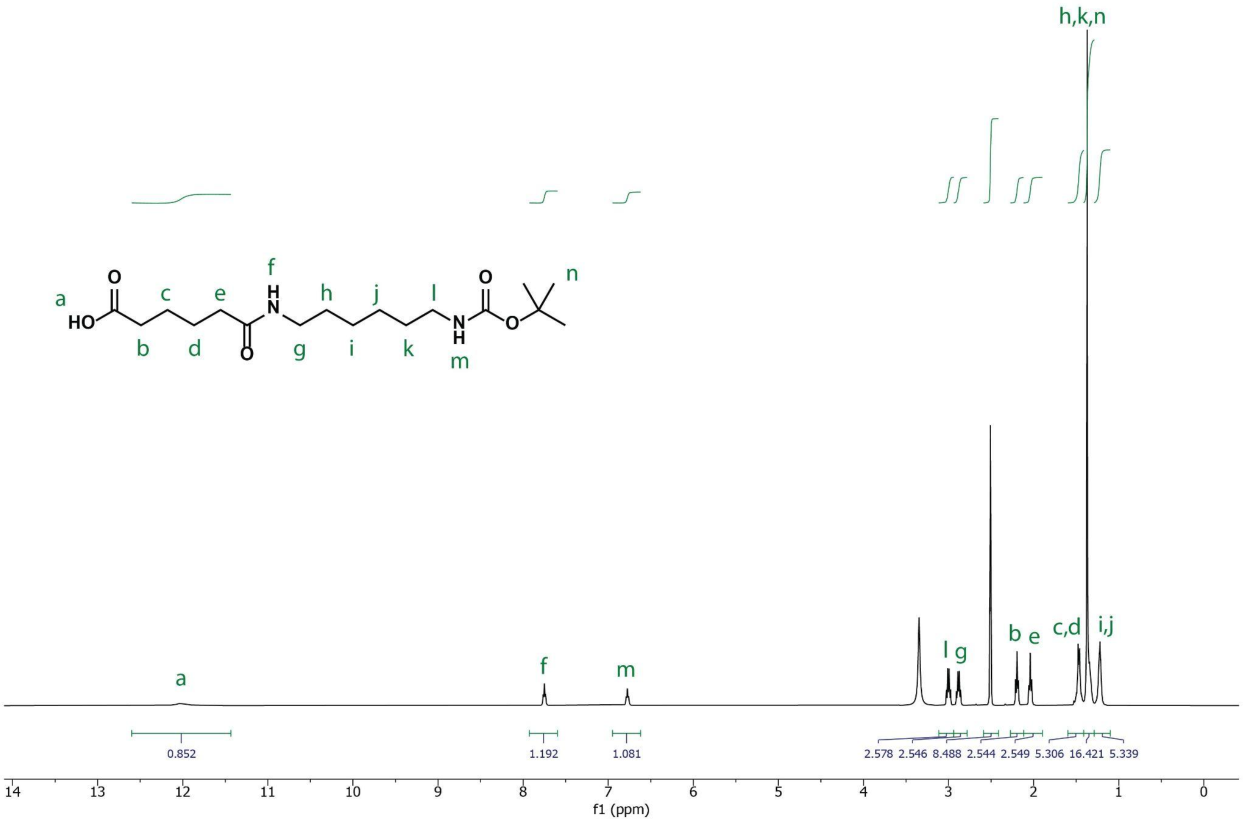


**Figure S4: $^{1}$H NMR spectrum of BocHN-MA in DMSO-$d_6$.**

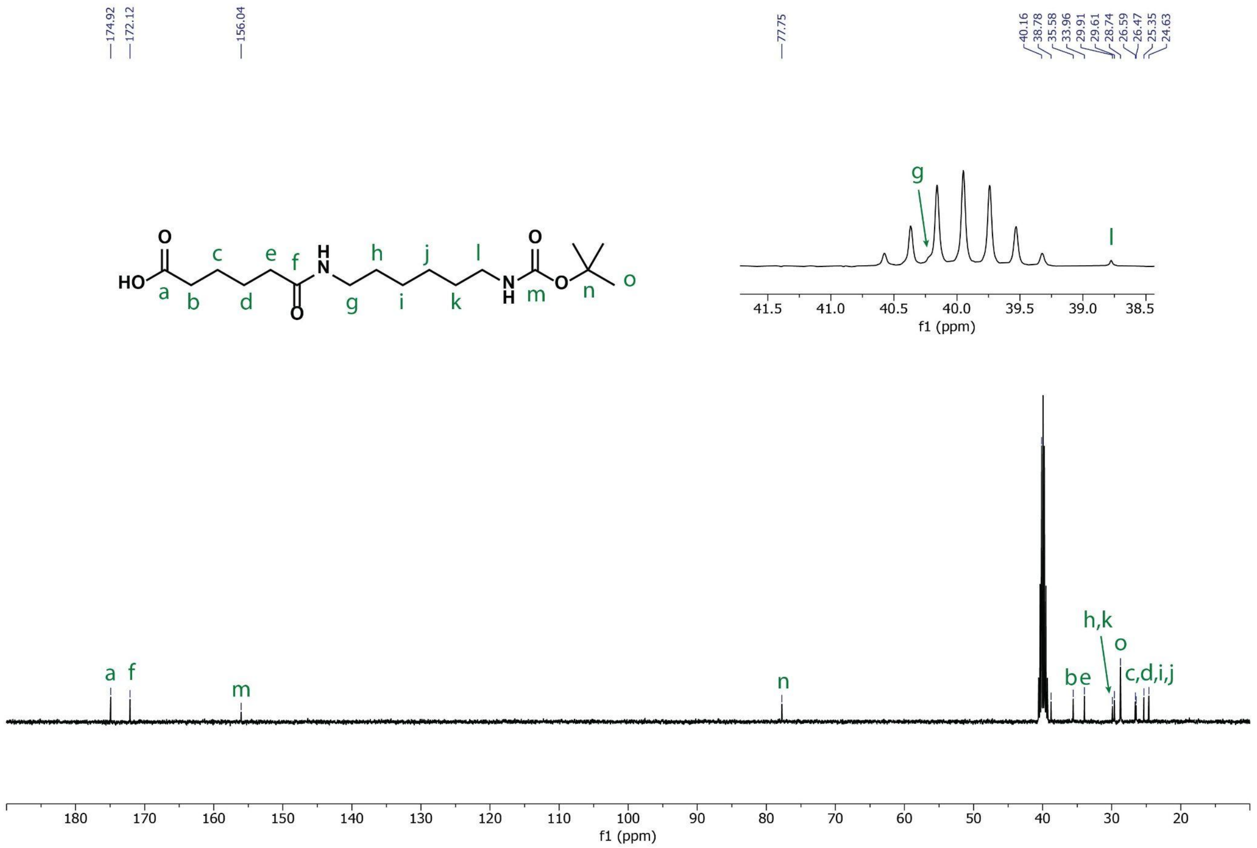


**Figure S5: $^{13}$C NMR spectrum of BocHN-MA in DMSO-$d_6$.**

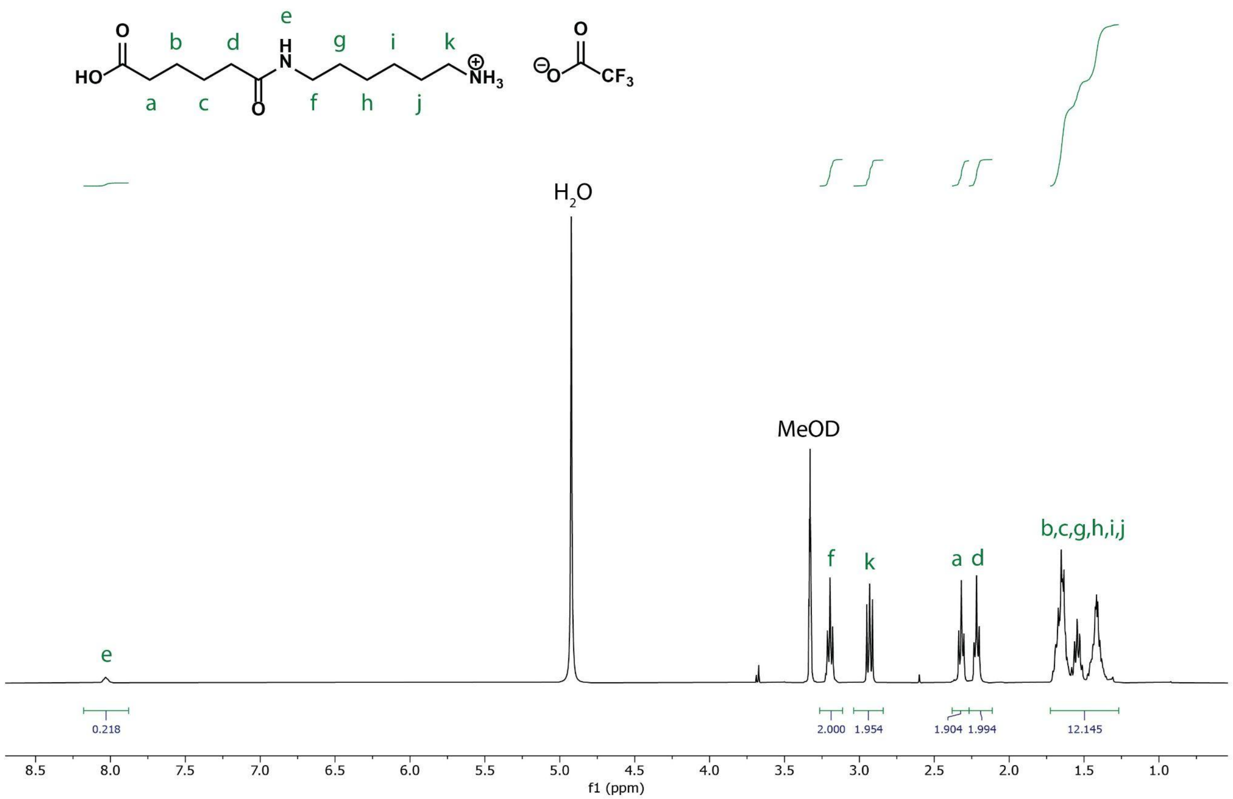


**Figure S6: $^1$H NMR spectrum of MA TFA salt in MeOD.**

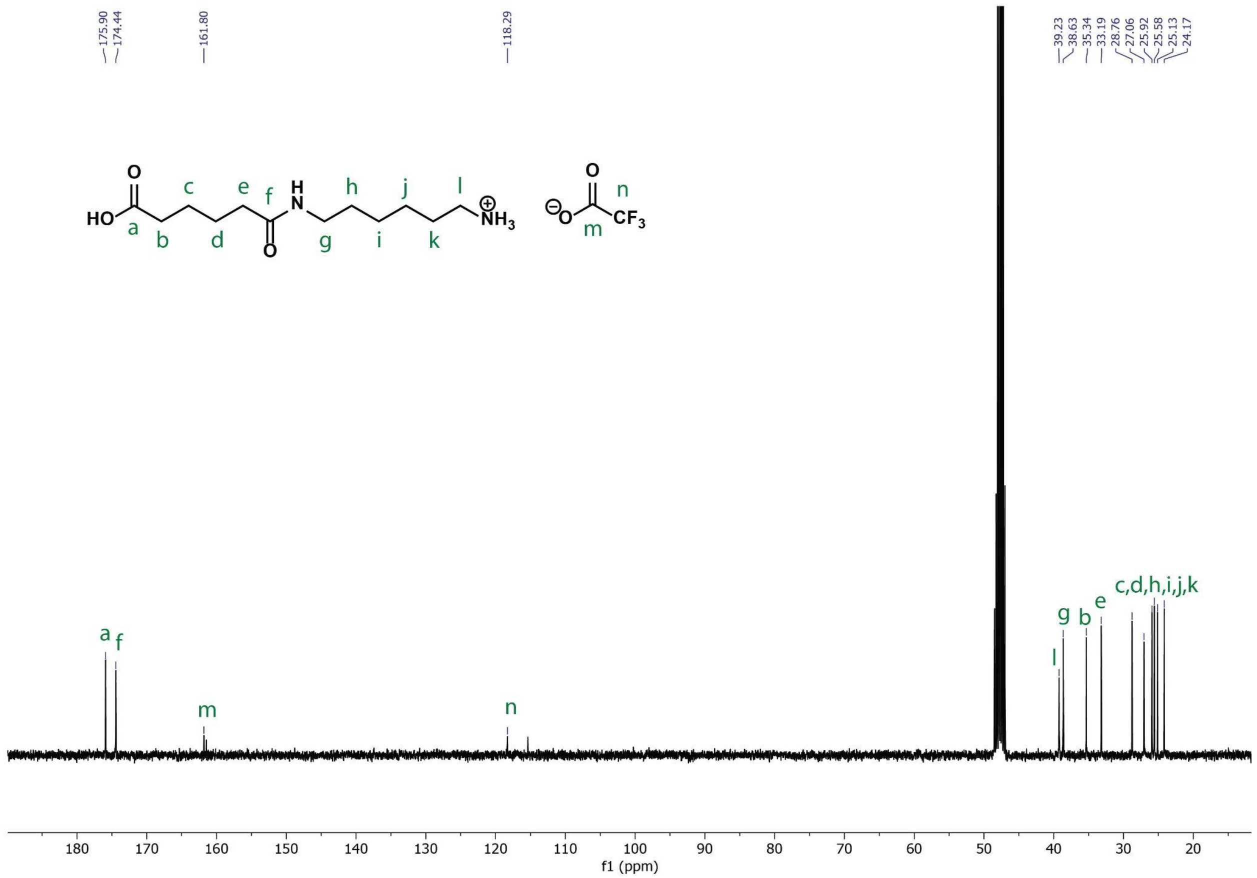


**Figure S7: $^{13}$C NMR spectrum of MA TFA salt in MeOD.**

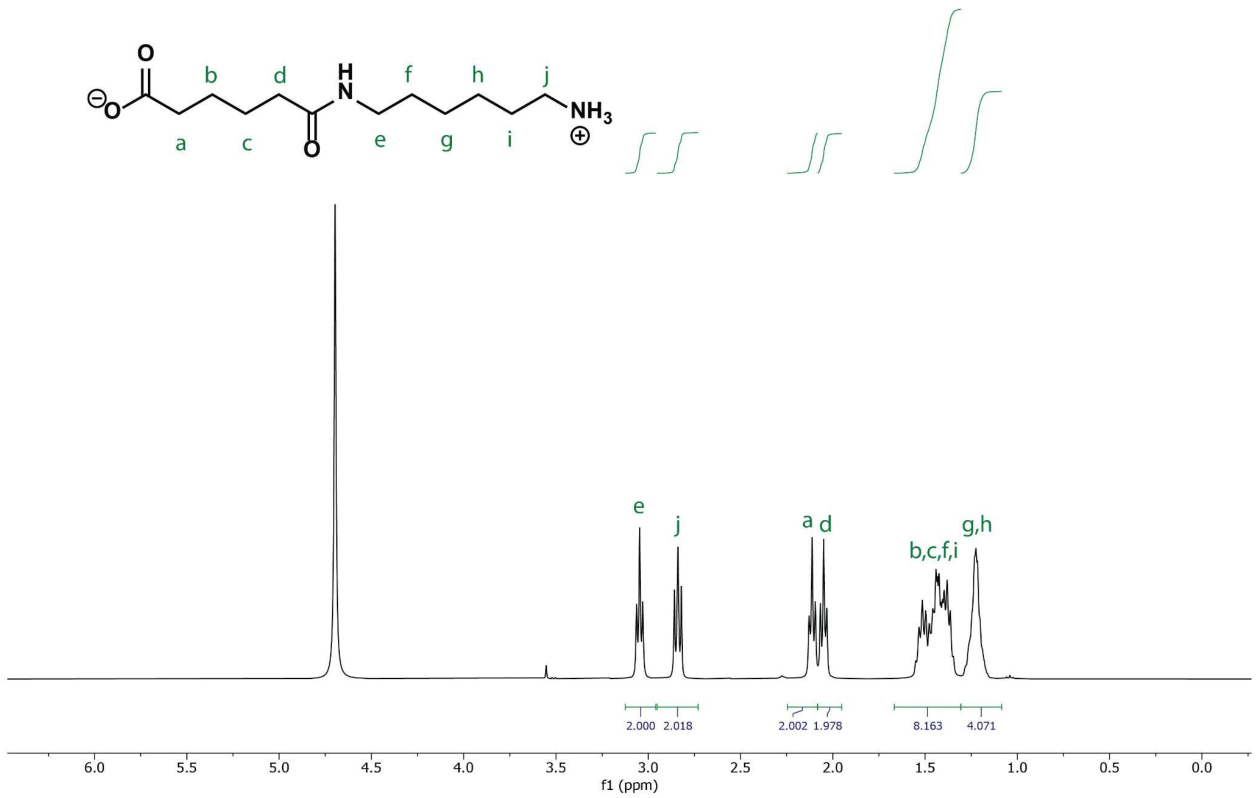


**Figure S8: [1]H NMR spectrum of MA in $D_2O$.**

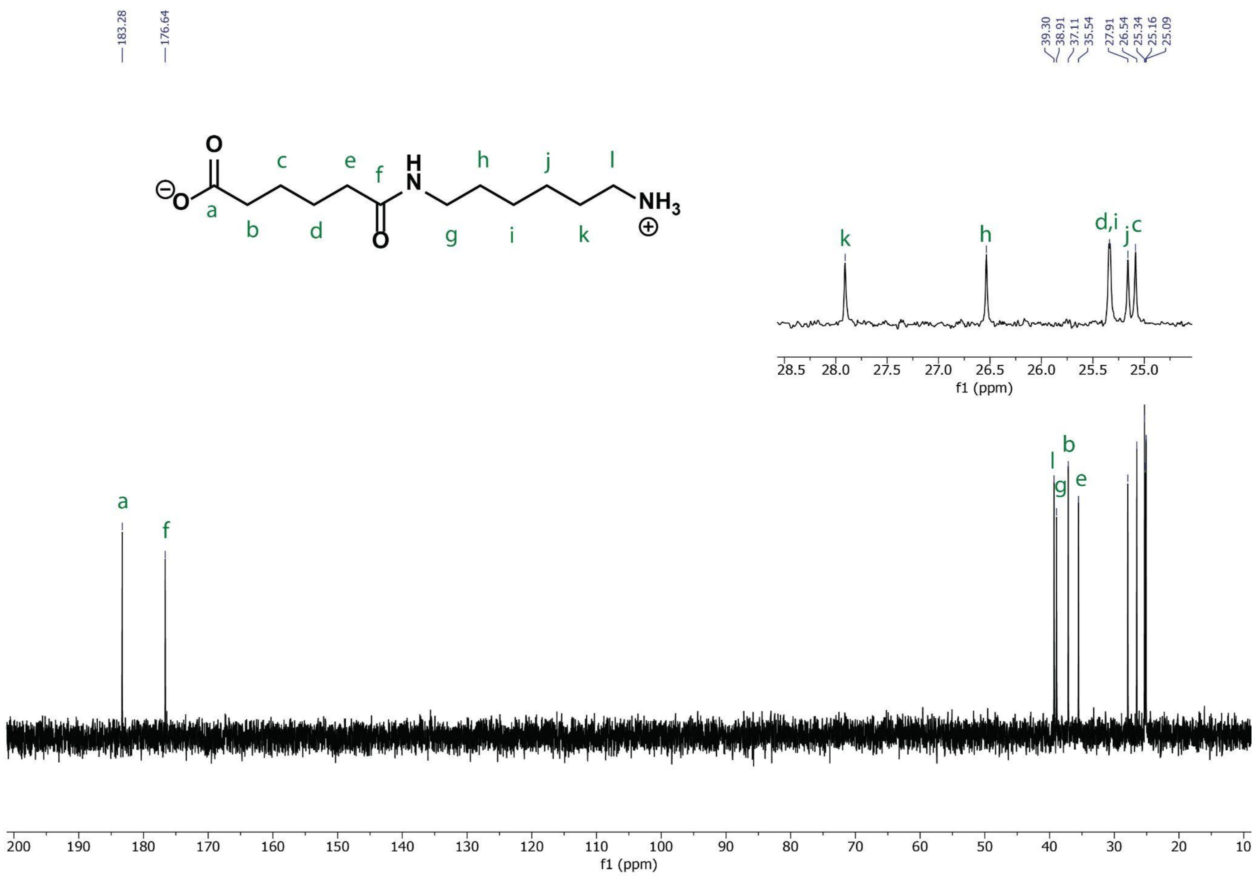


**Figure S9: $^{13}C$ NMR spectrum of MA in $D_2O$.**

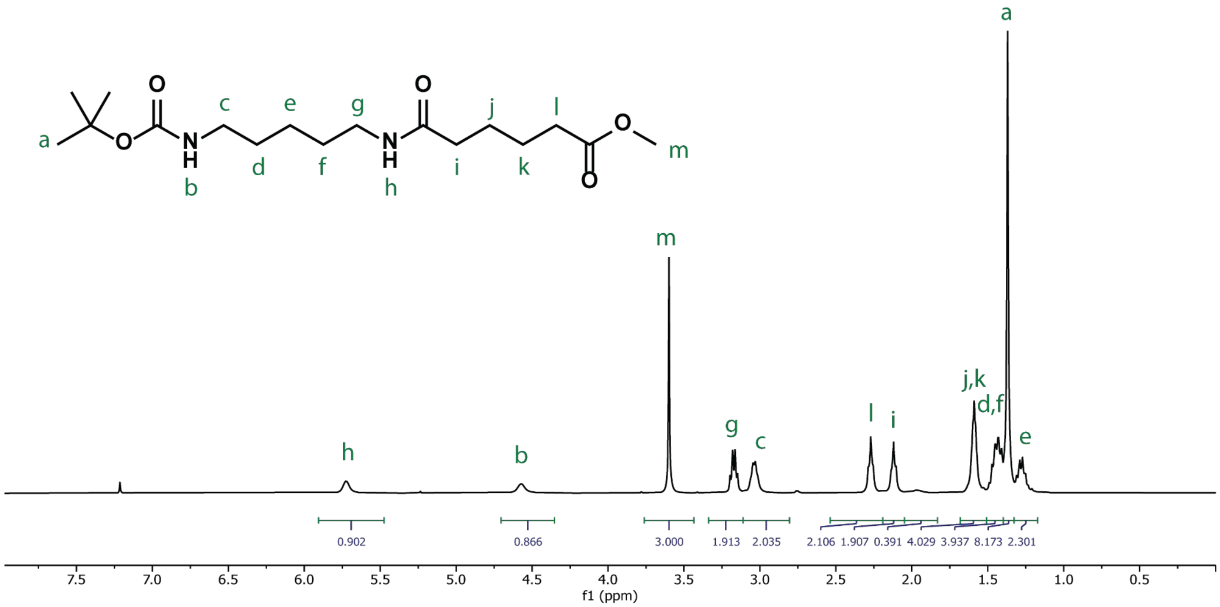


**Figure S10: [1]H NMR spectrum of Boc-CA-OMe in $CDCl_3$.**

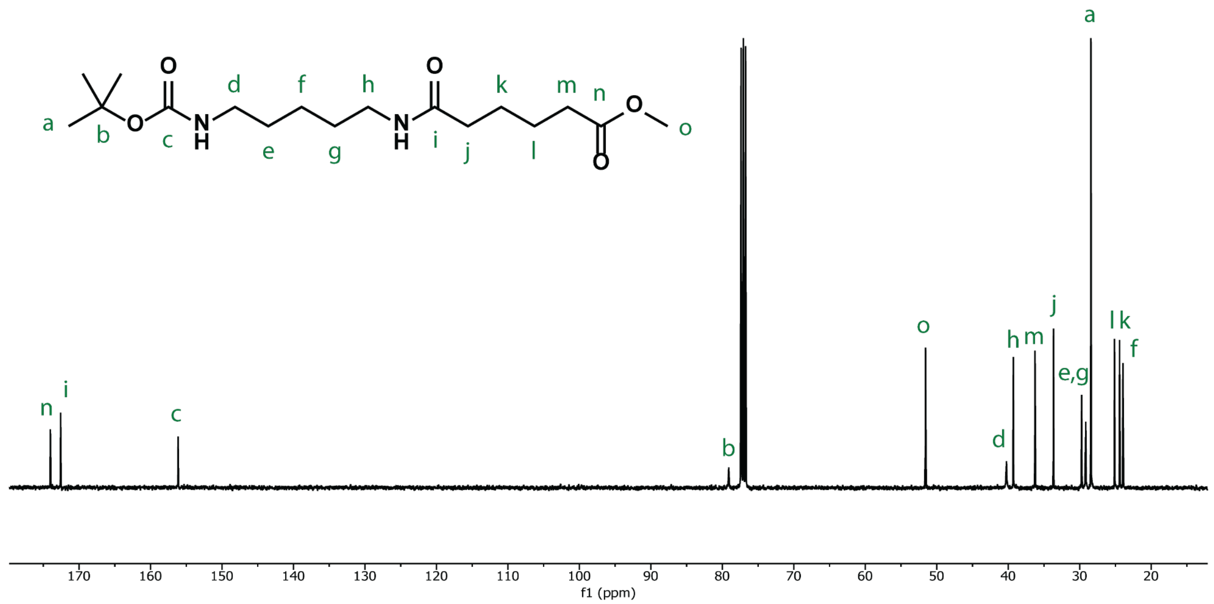


**Figure S11: 13C NMR spectrum of Boc-CA-OMe in $CDCl_3$.**

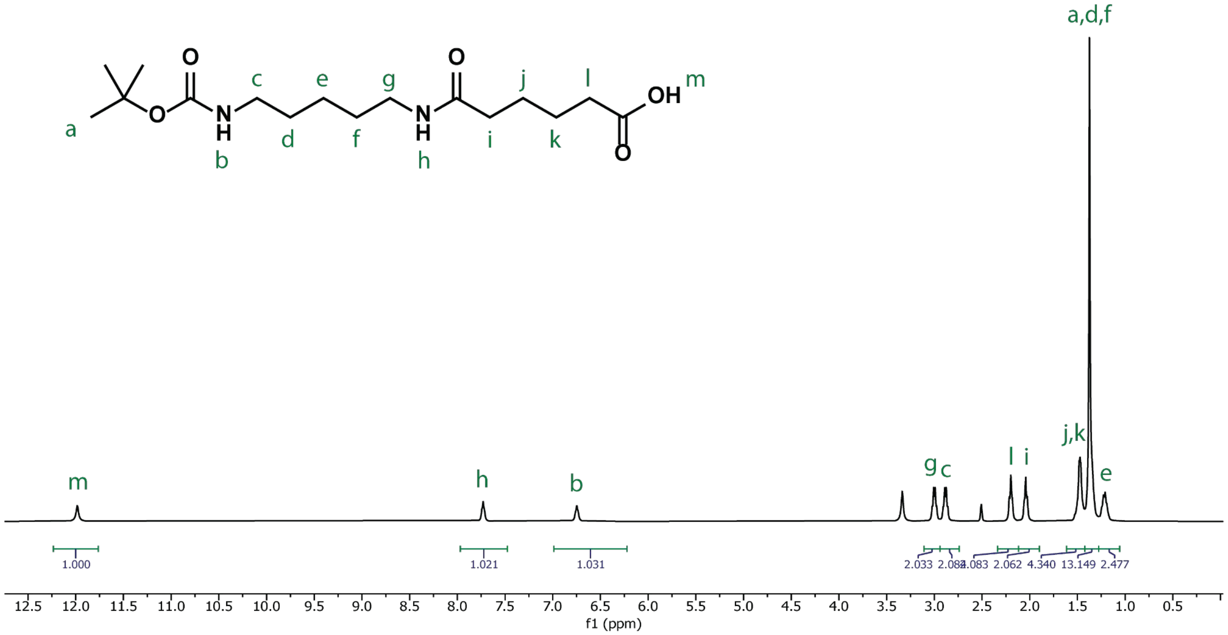


**Figure S12: [1]H NMR spectrum of Boc-CA in DMSO-d6.**

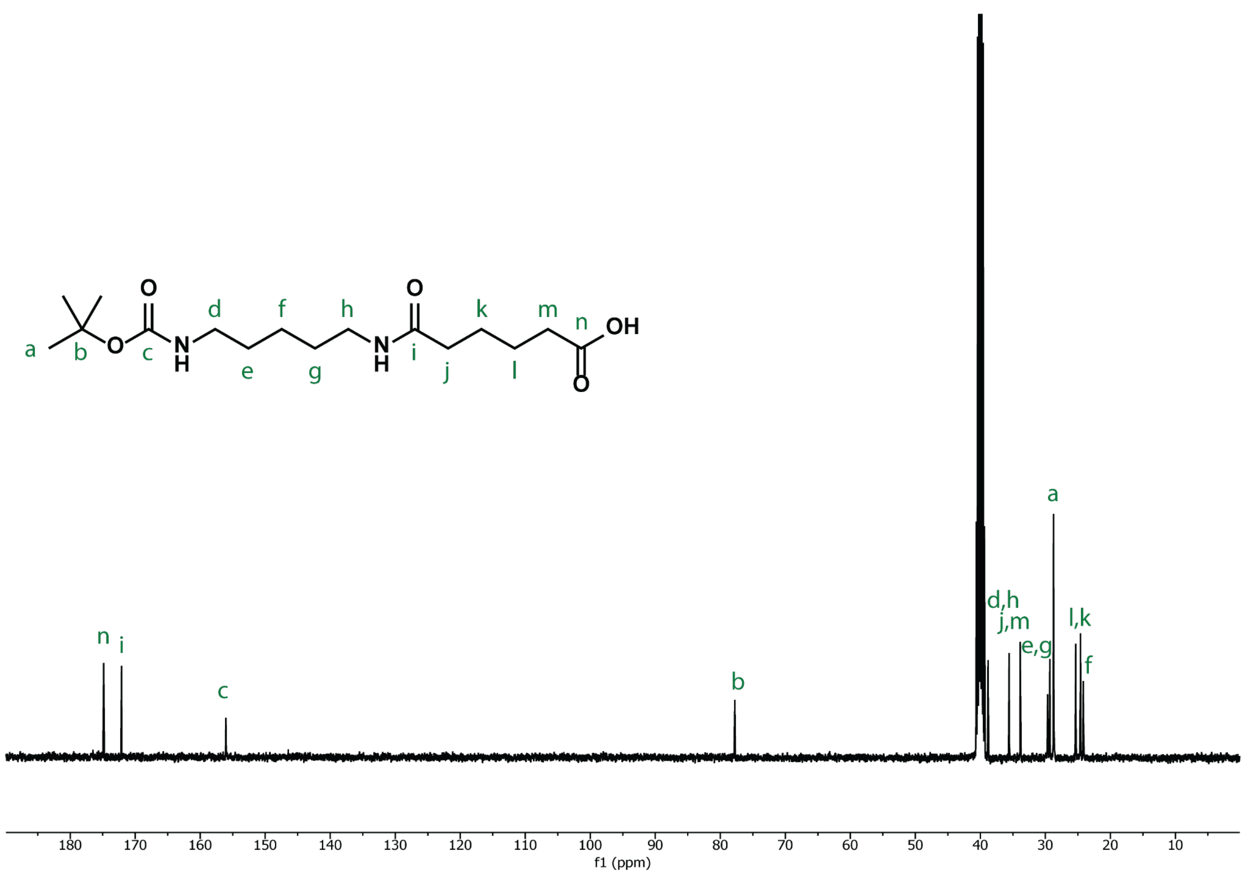


**Figure S13: $^{13}$C NMR spectrum of Boc-CA in DMSO-$d_6$.**

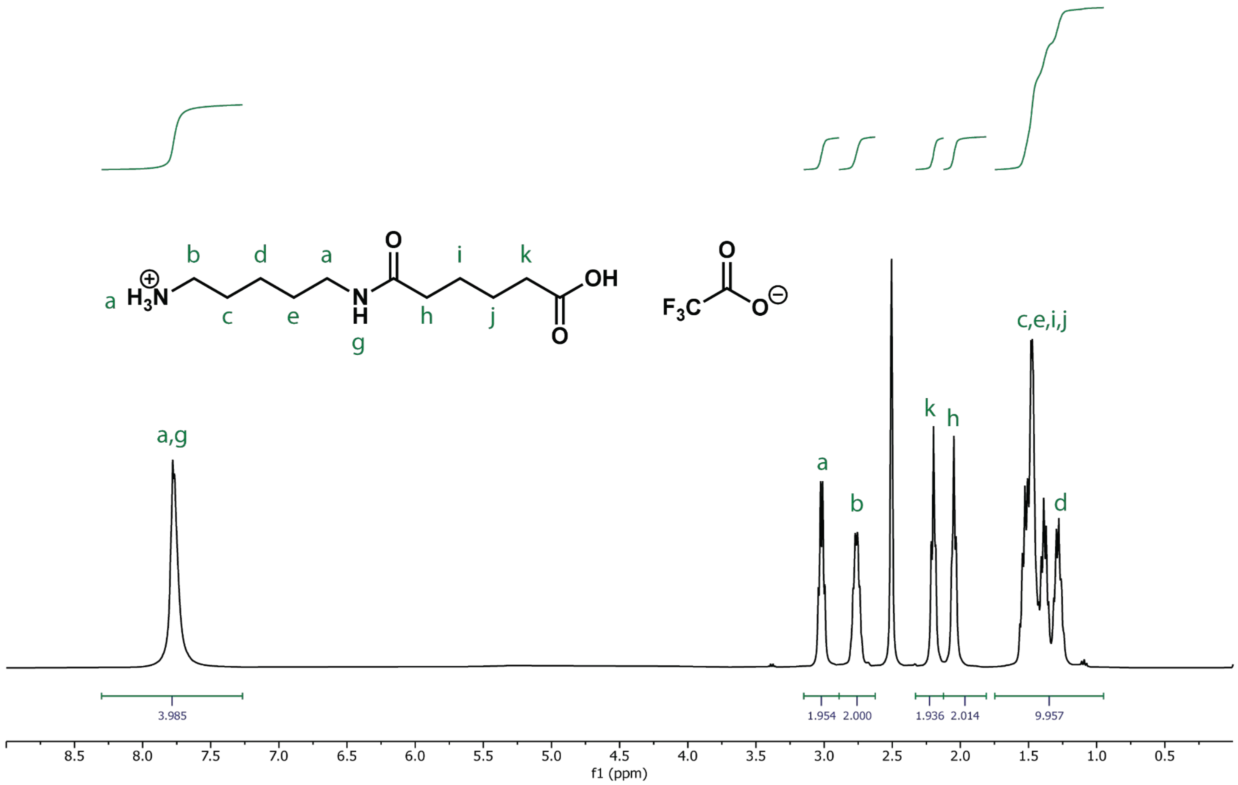


**Figure S14: [1]H NMR spectrum of CA TFA salt in DMSO-d₆.**

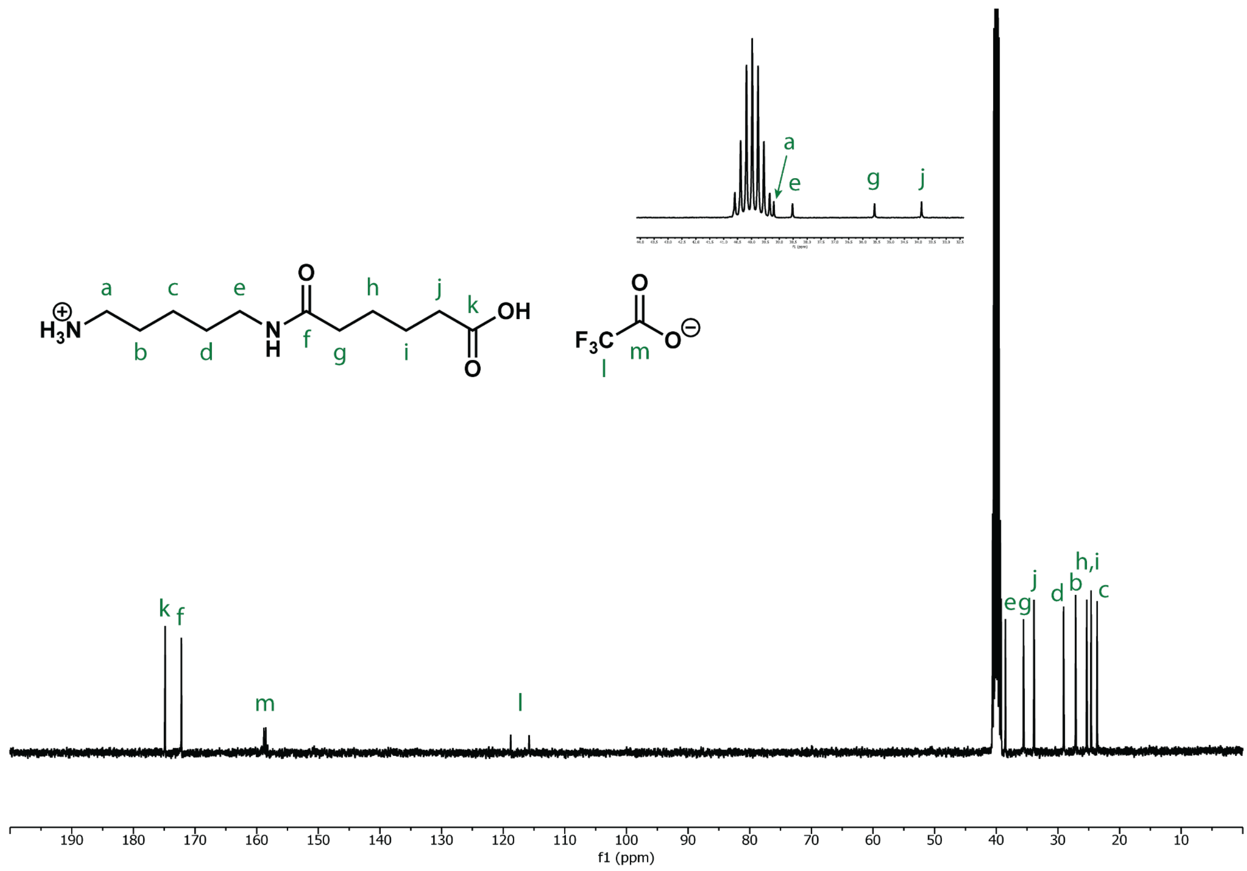


**Figure S15: $^{13}$C NMR spectrum of CA TFA salt in DMSO-d$_6$.**

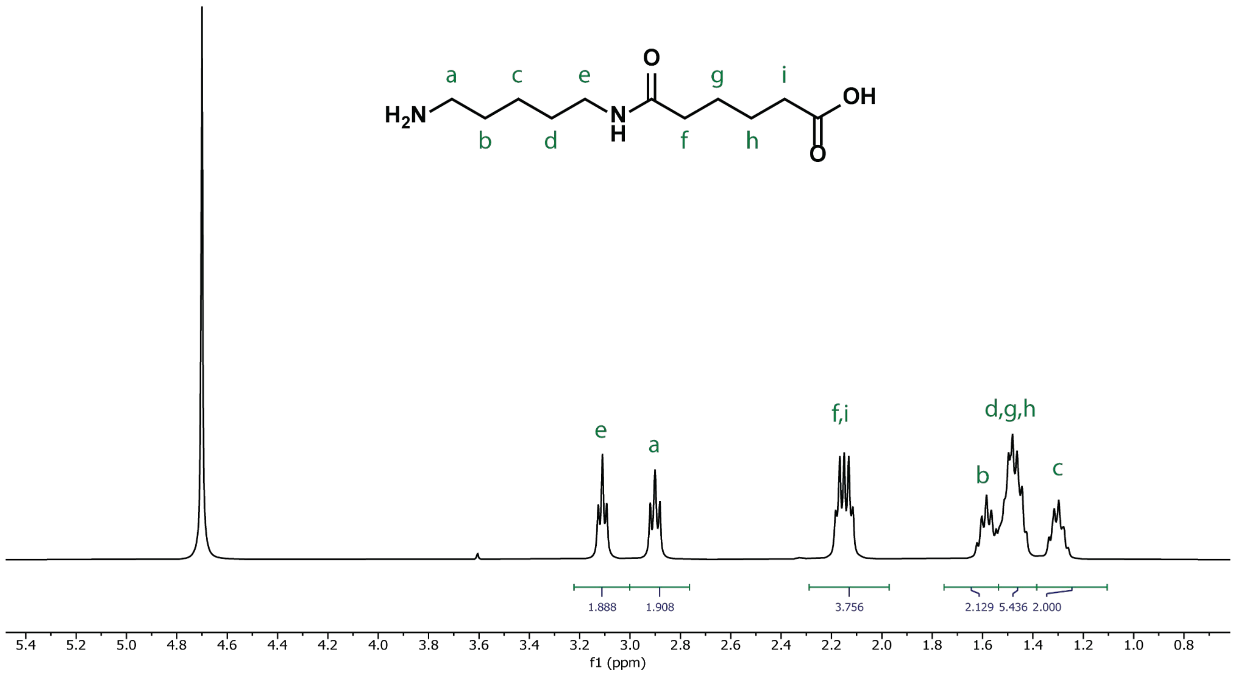


**Figure S16: [1]H NMR spectrum of CA in $D_2O$.**

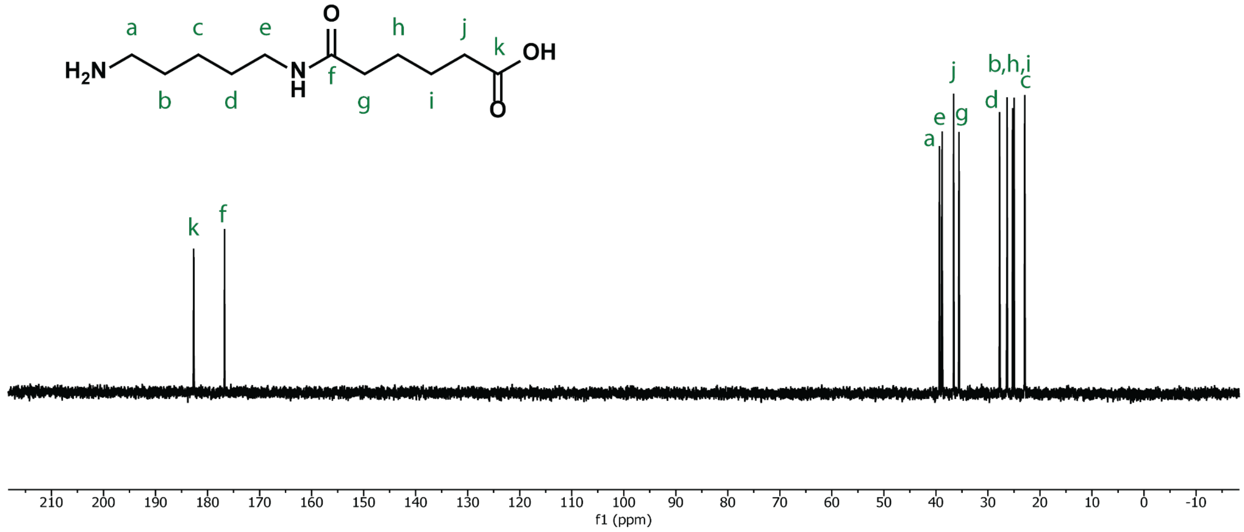


**Figure S17: $^{13}C$ NMR spectrum of CA in $D_2O$.**

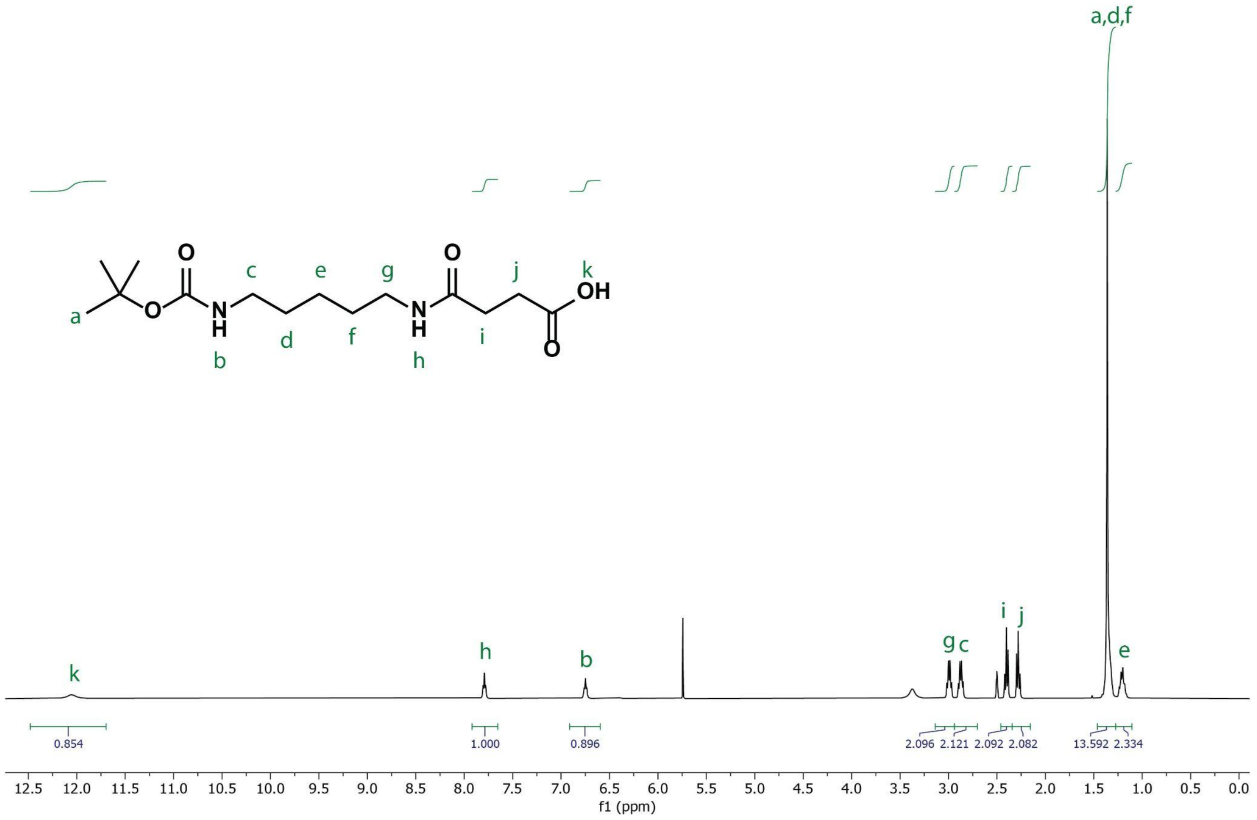


**Figure S18: [1]H NMR spectrum of BocHN-CS in DMSO-d₆.**

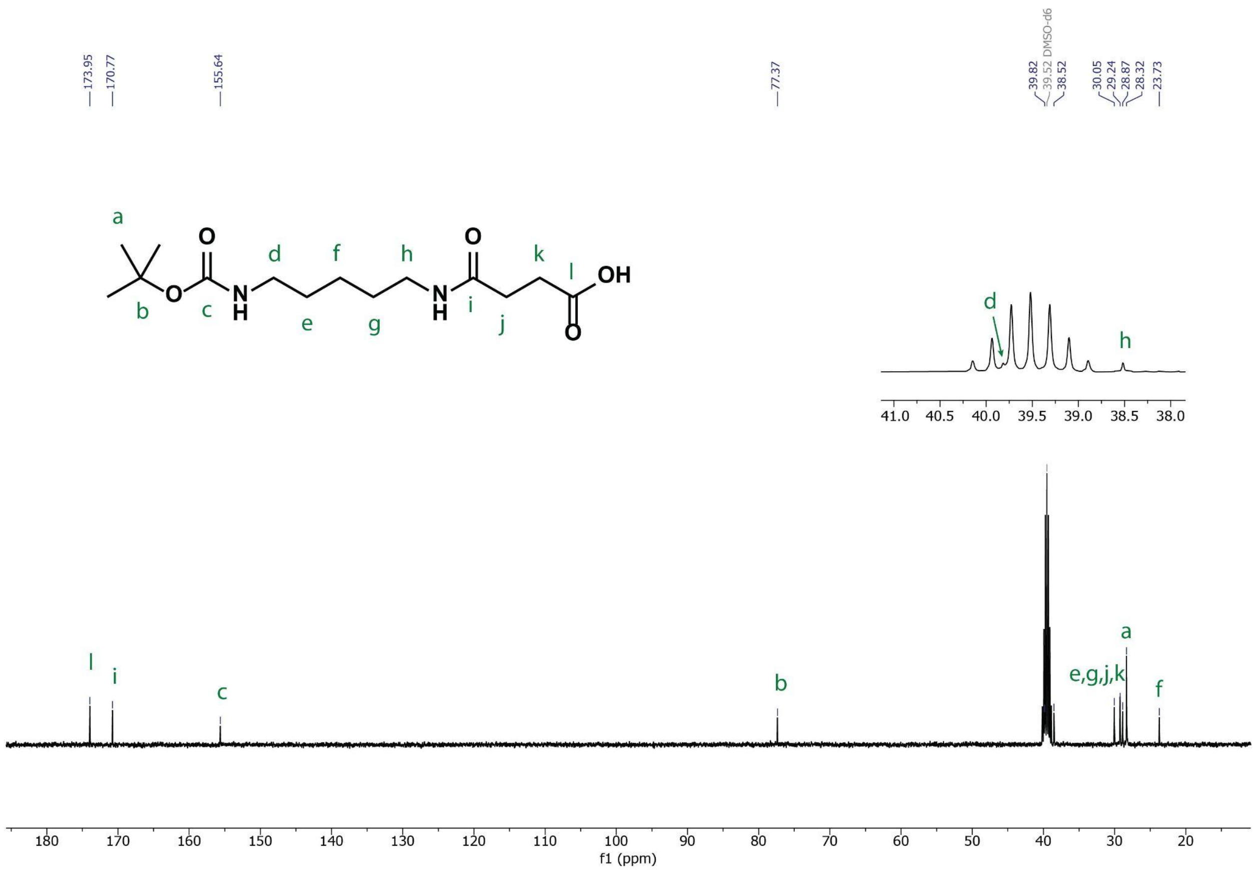


**Figure S19: $^{13}$C NMR spectrum of BocHN-CS in DMSO-$d_6$.**

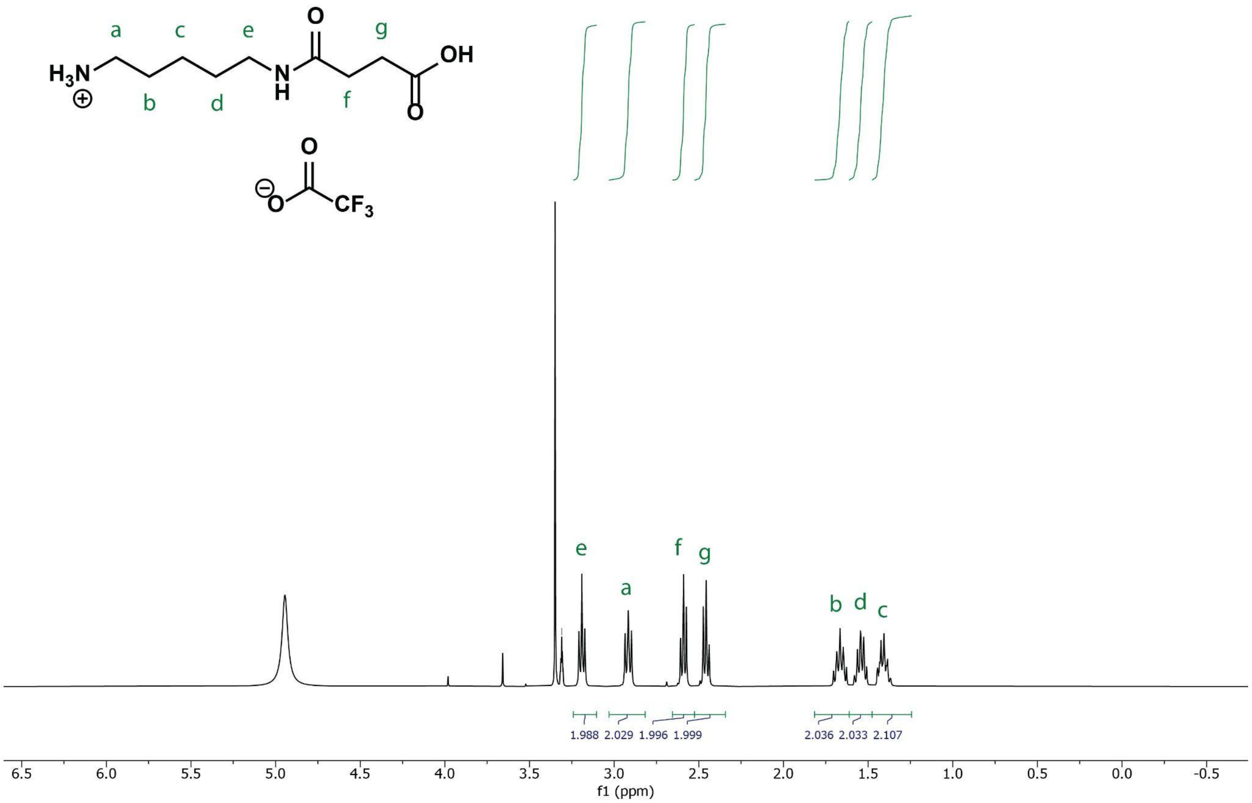


**Figure S20: [1]H NMR spectrum of CS TFA salt in MeOD.**

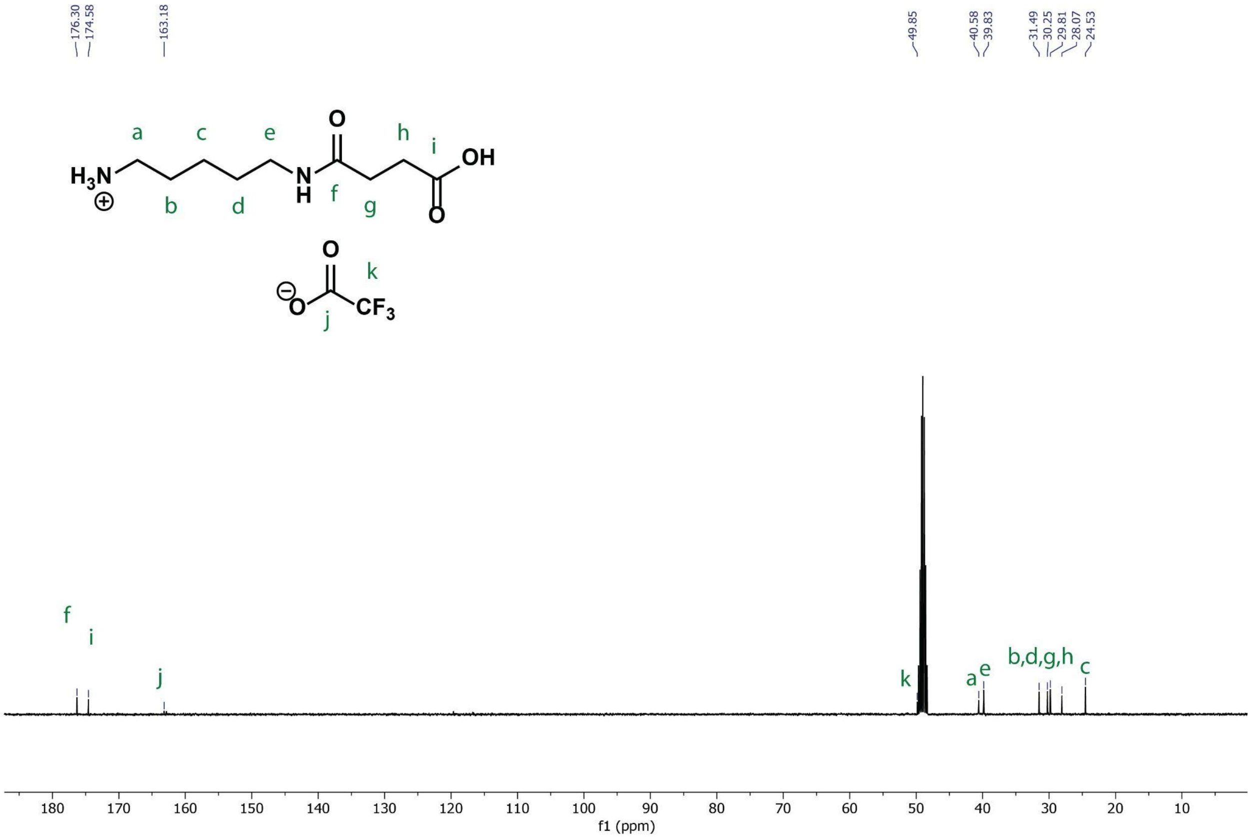


**Figure S21: $^{13}$C NMR spectrum of CS TFA salt in MeOD.**

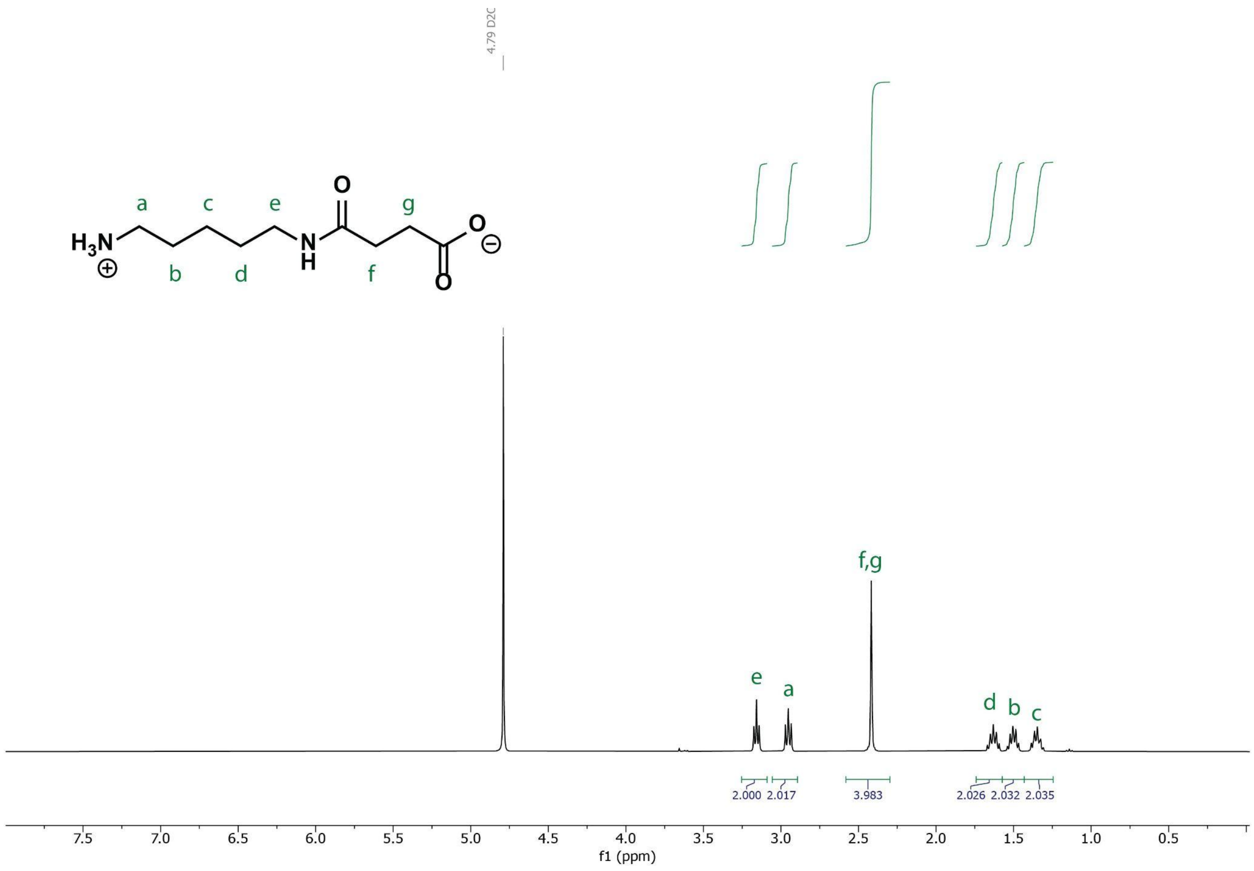


**Figure S22: $^1$H NMR spectrum of CS in $D_2O$.**

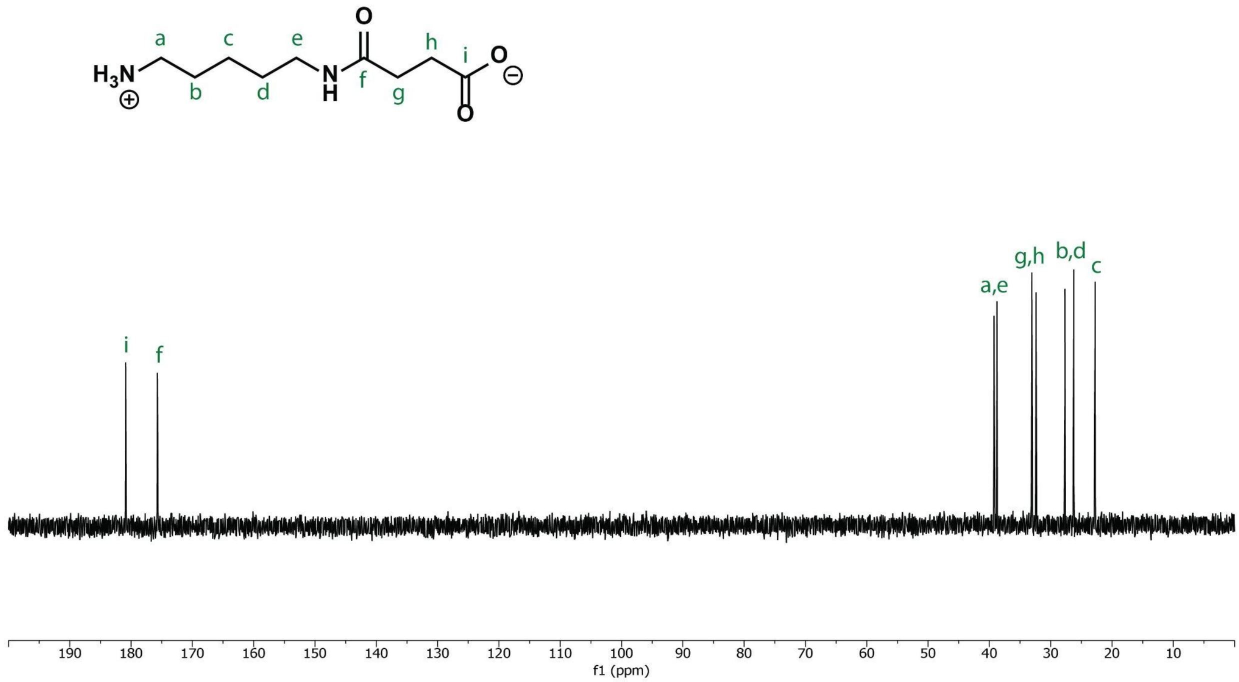


**Figure S23: $^{13}$C NMR spectrum of CS in $D_2O$.**

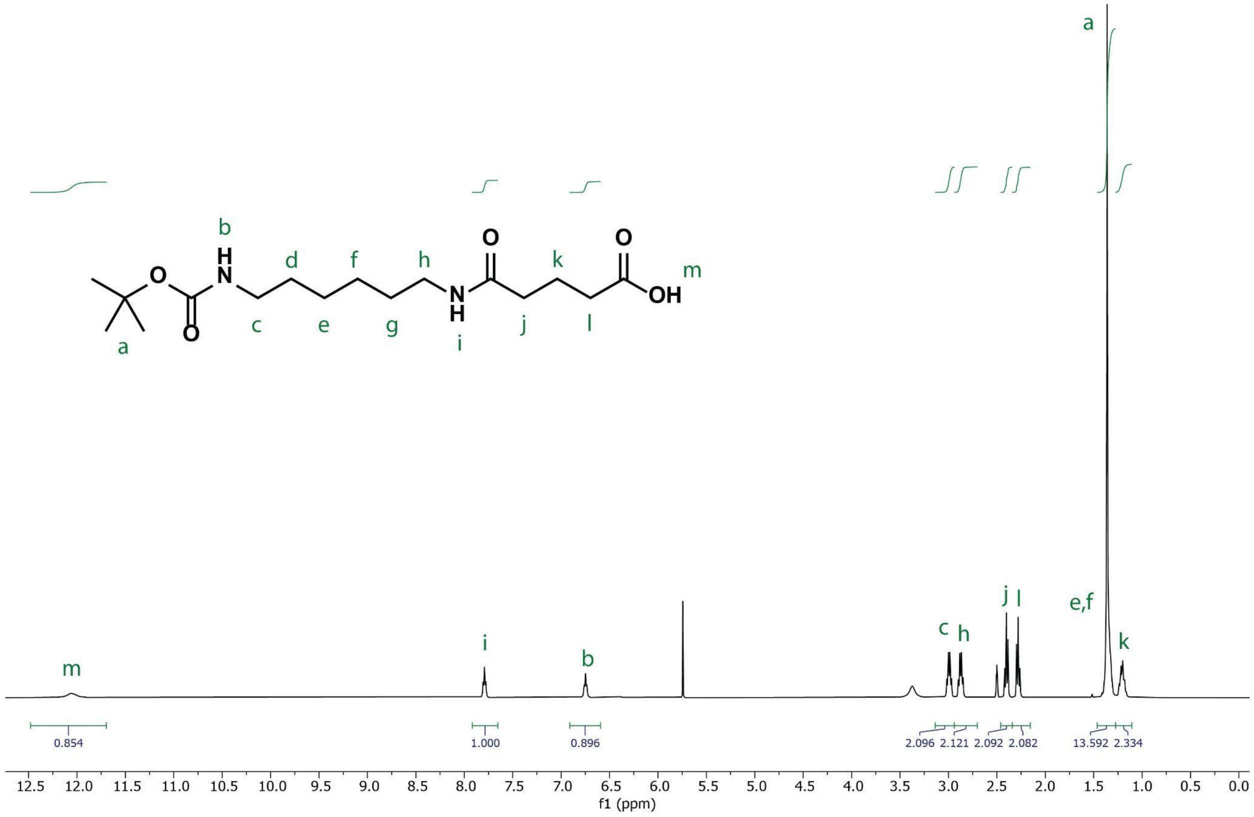


**Figure S24: $^{1}$H NMR spectrum of BocHN-MG in DMSO-d$_6$**

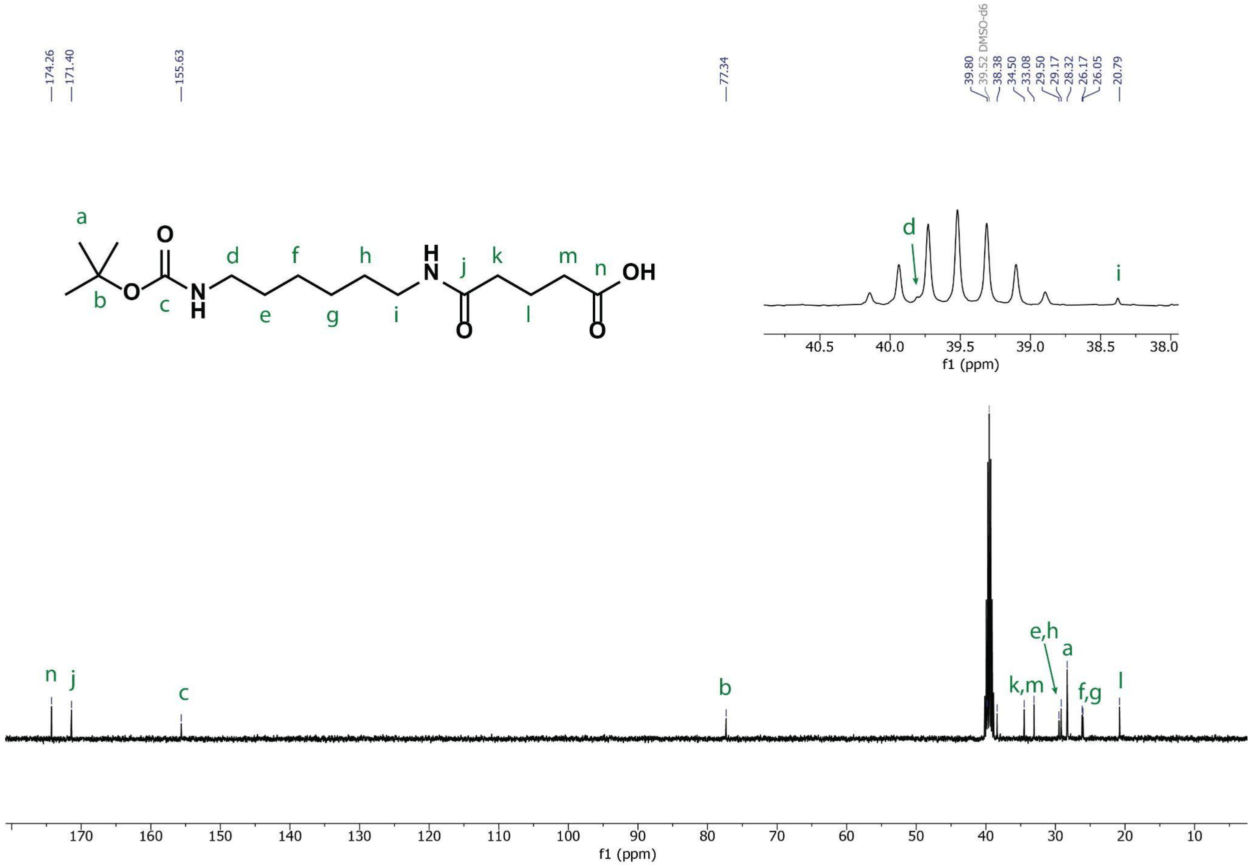


**Figure S25: $^{13}$C NMR spectrum of BocHN-MG in DMSO-d$_6$.**

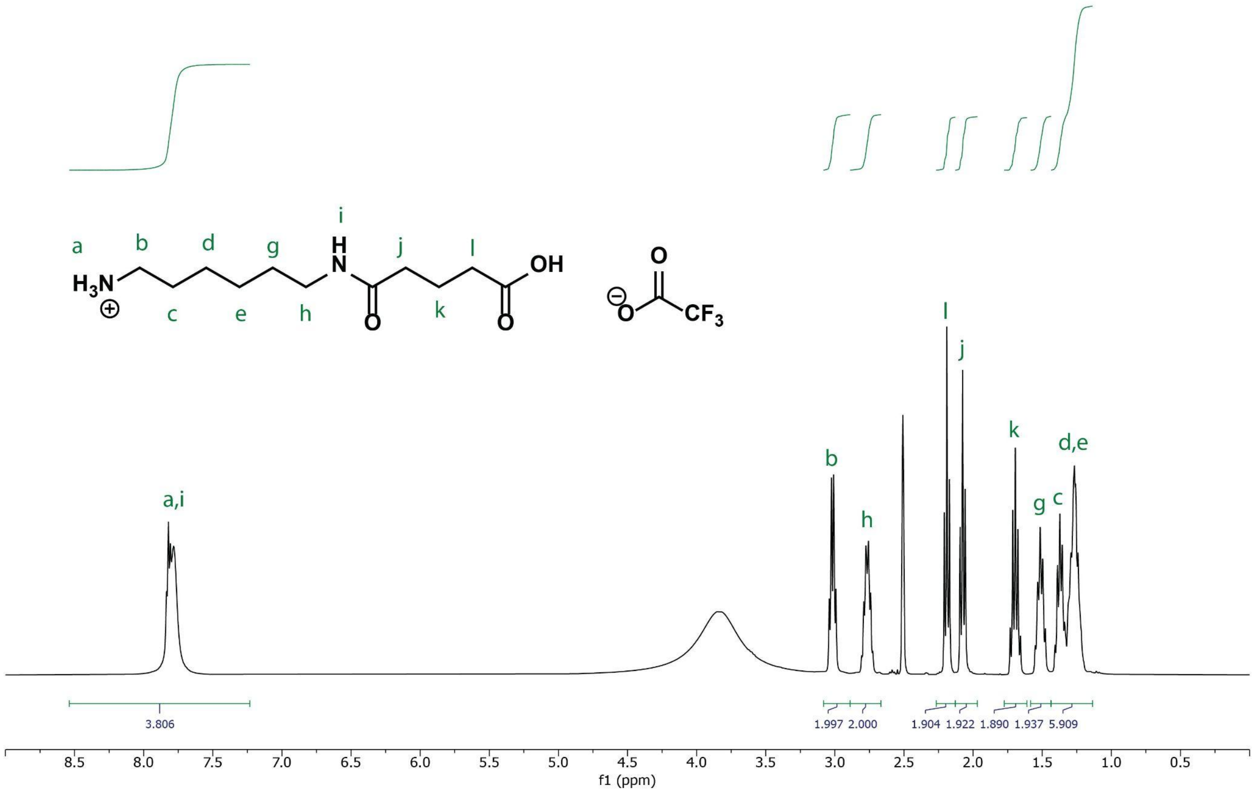


**Figure S26: ${}^{1}$H NMR spectrum of MG TFA salt in DMSO-$d_6$.**

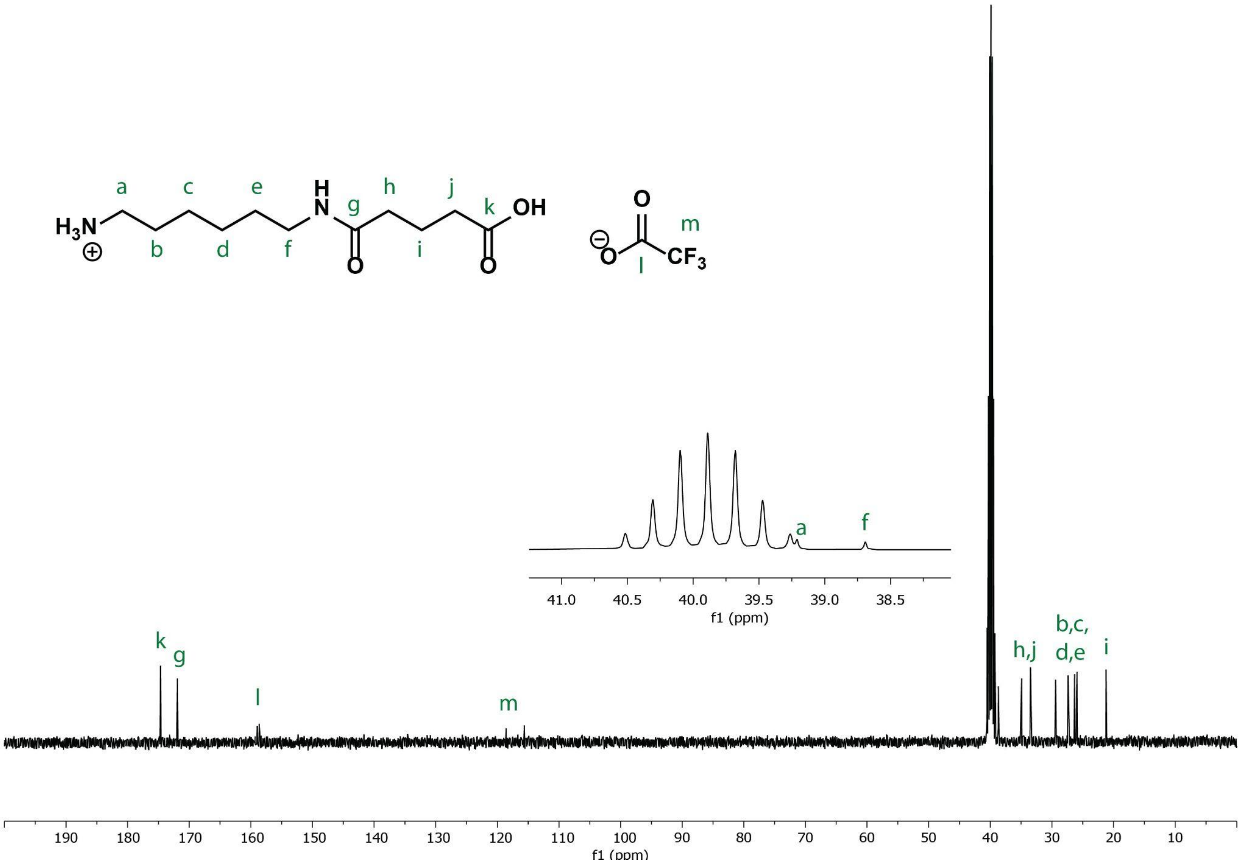


**Figure S27: $^{13}$C NMR spectrum of MG TFA salt in DMSO-d$_6$.**

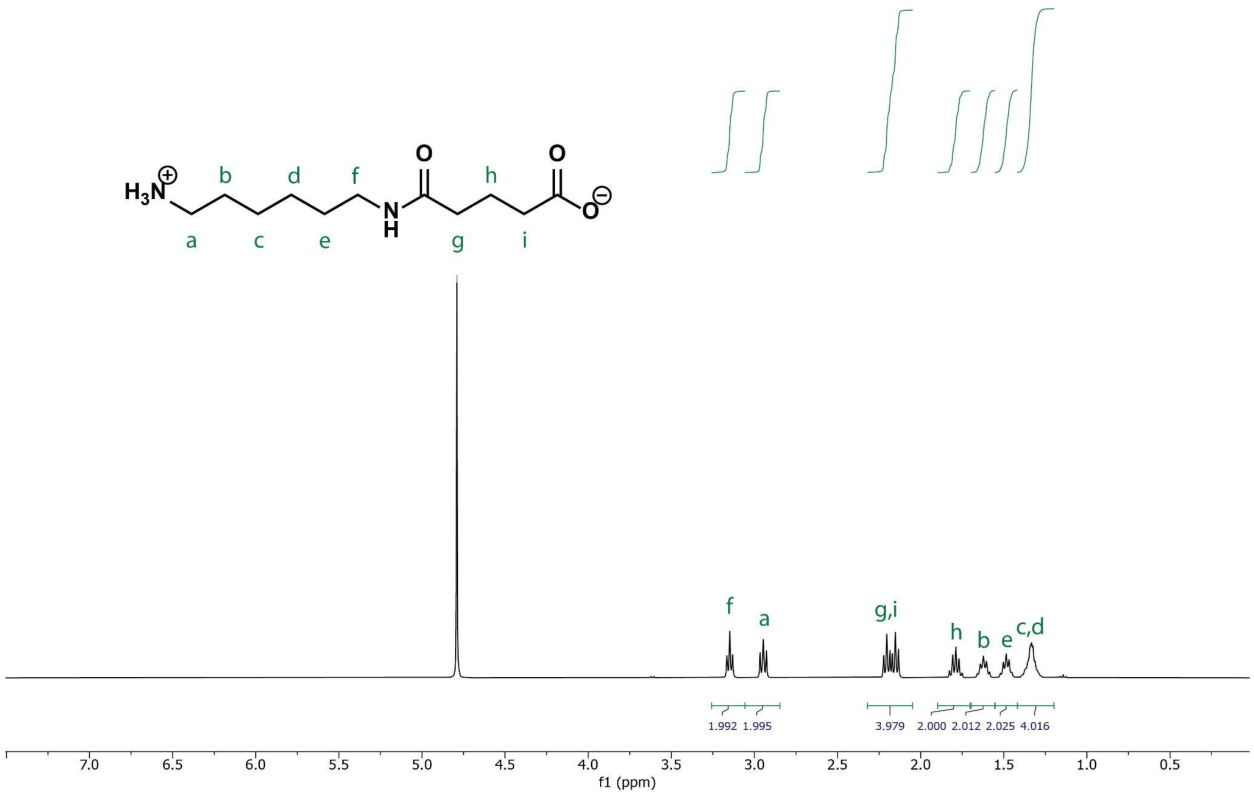


**Figure S28: $^{1}$H NMR spectrum of MG in $D_2O$.**

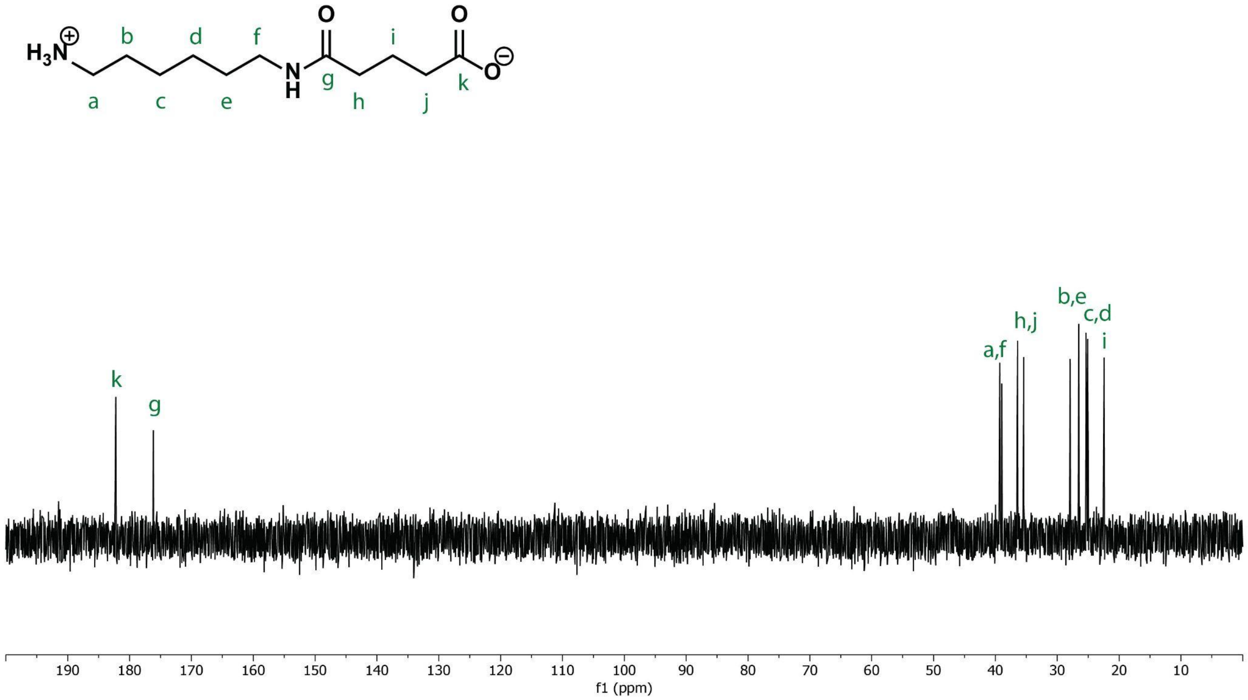


**Figure S29: $^{13}$C NMR spectrum of MG in $D_2O$.**

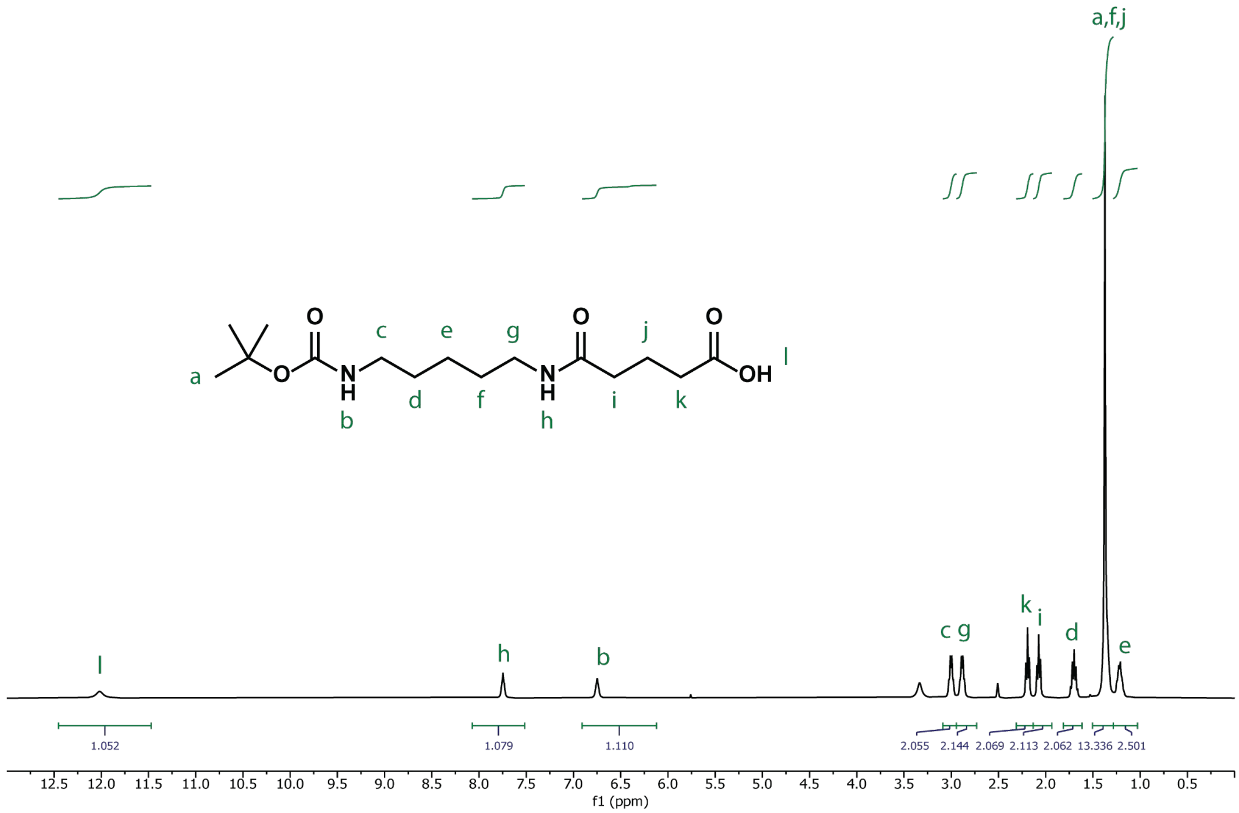


**Figure S30: $^{1}$H NMR spectrum of Boc-CG in DMSO-d$_6$.**

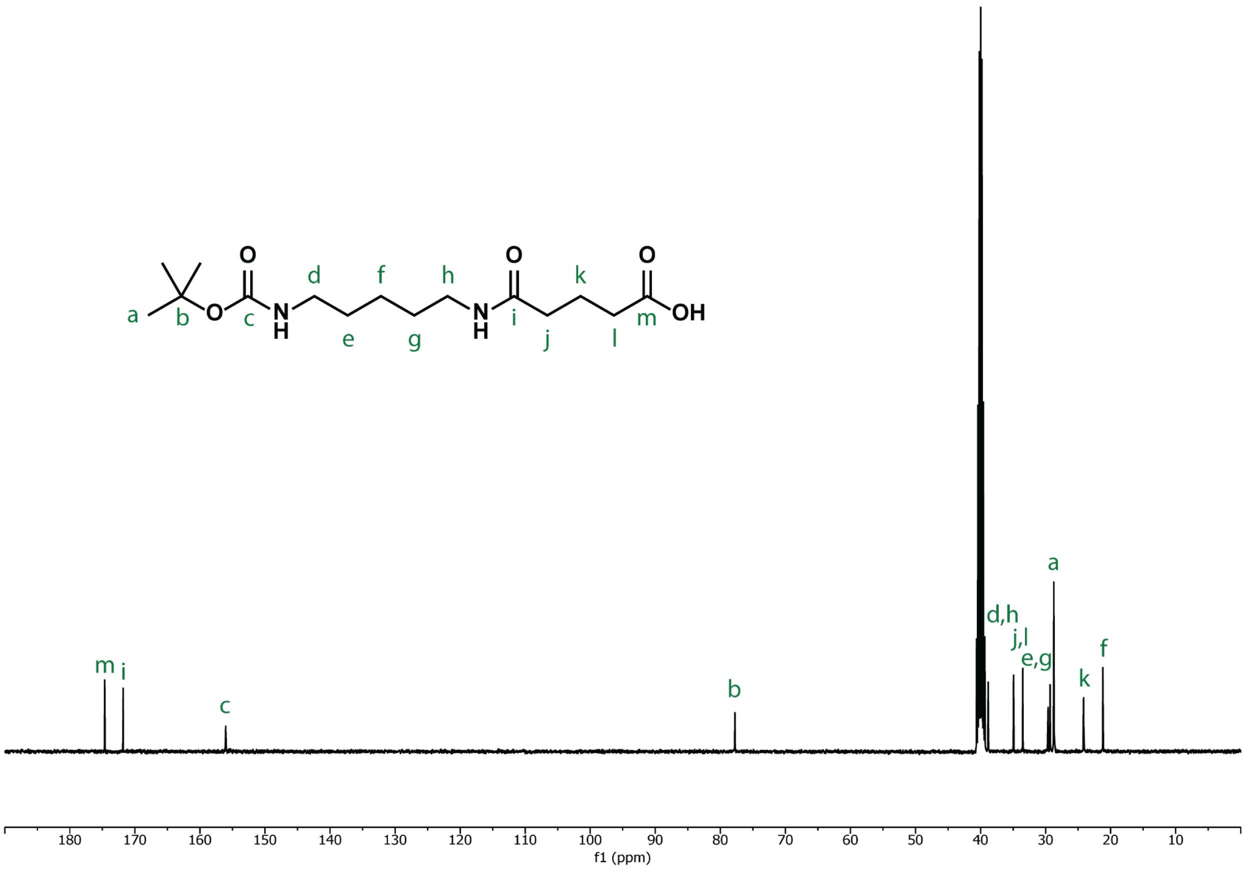


**Figure S31: [13]C NMR spectrum of Boc-CG in DMSO-d6.**

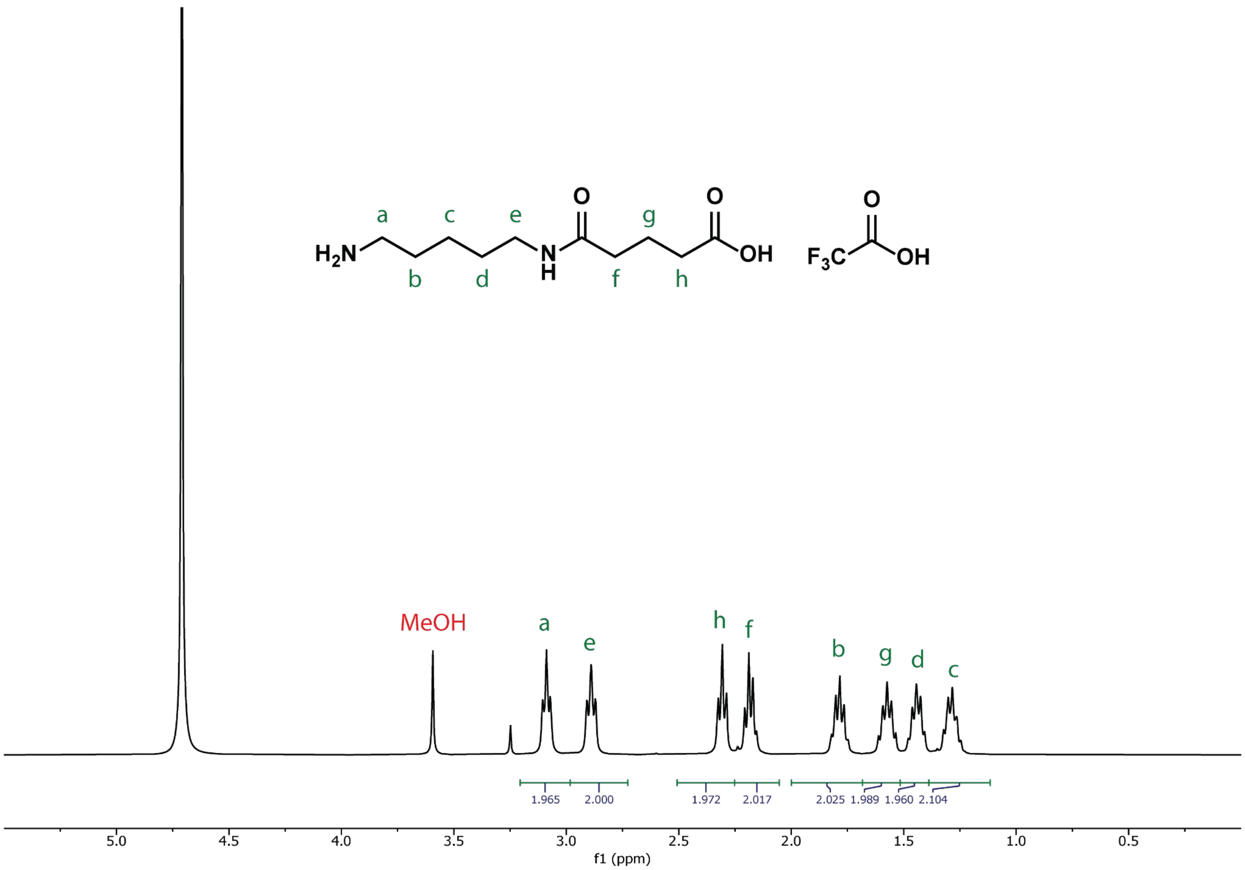


**Figure S32: $^1$H NMR spectrum of CG TFA salt in $D_2O$.**

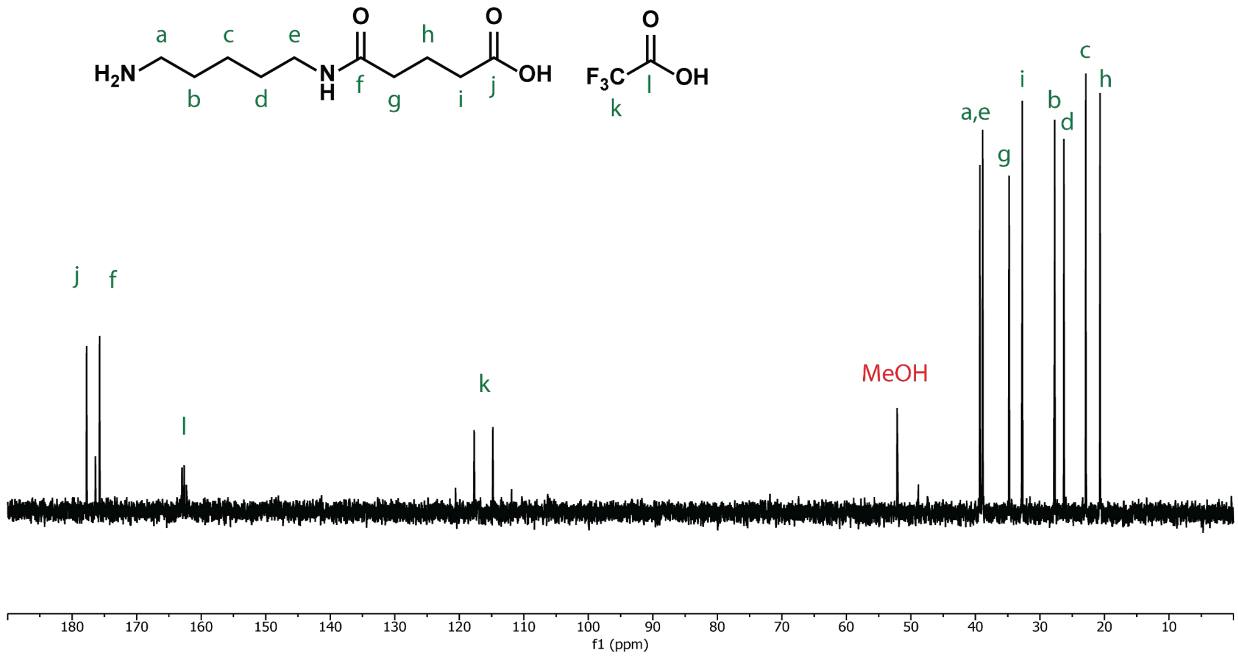


**Figure S33: $^{13}$C NMR spectrum of CG TFA salt in $D_2O$.**

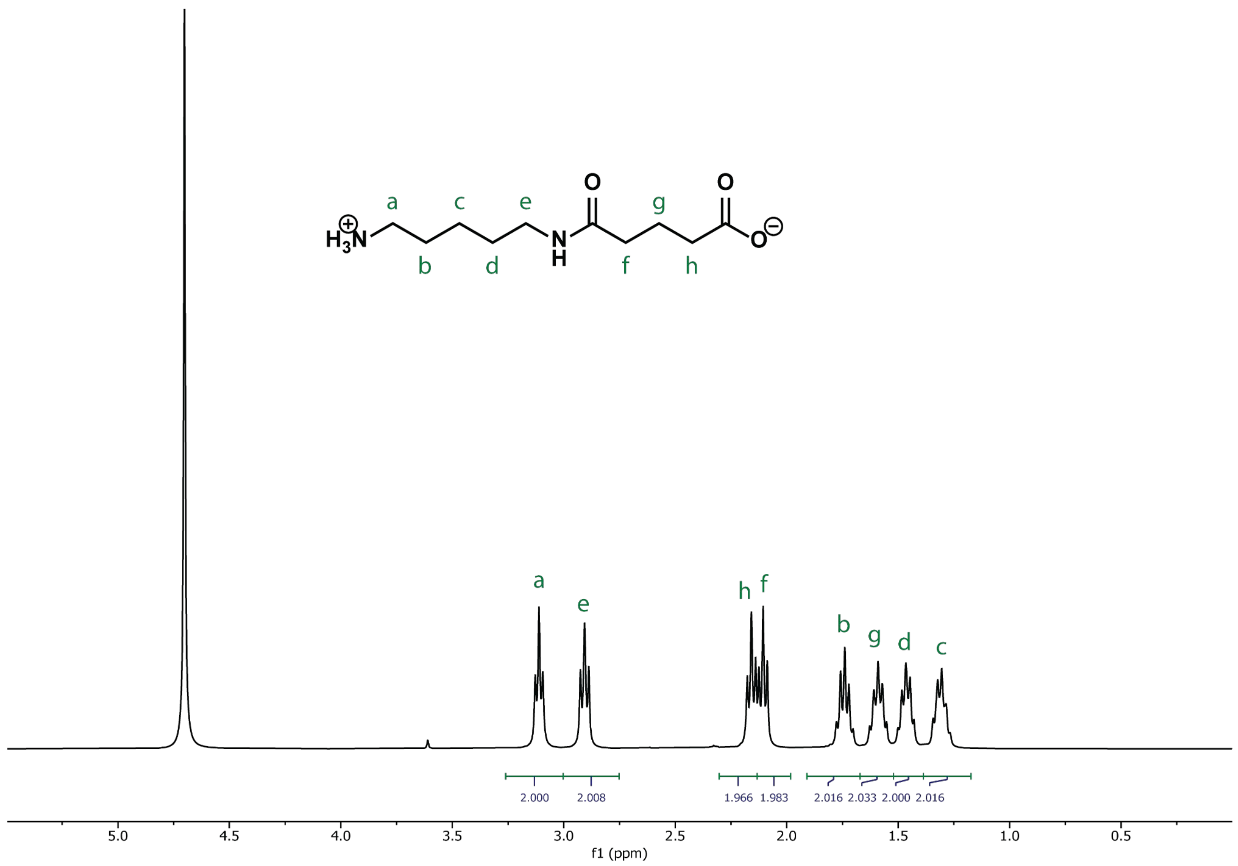


**Figure S34: $^1$H NMR spectrum of CG in $D_2O$.**

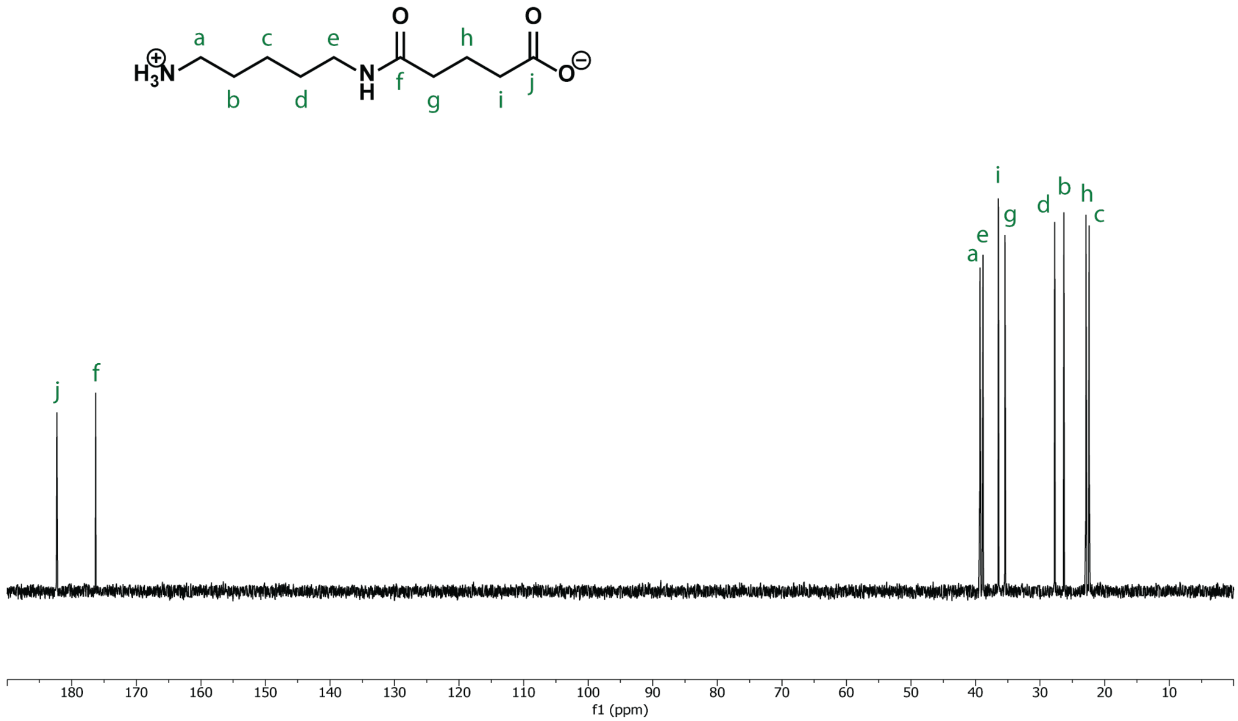


**Figure S35: $^{13}$C NMR spectrum of CG in $D_2O$.**

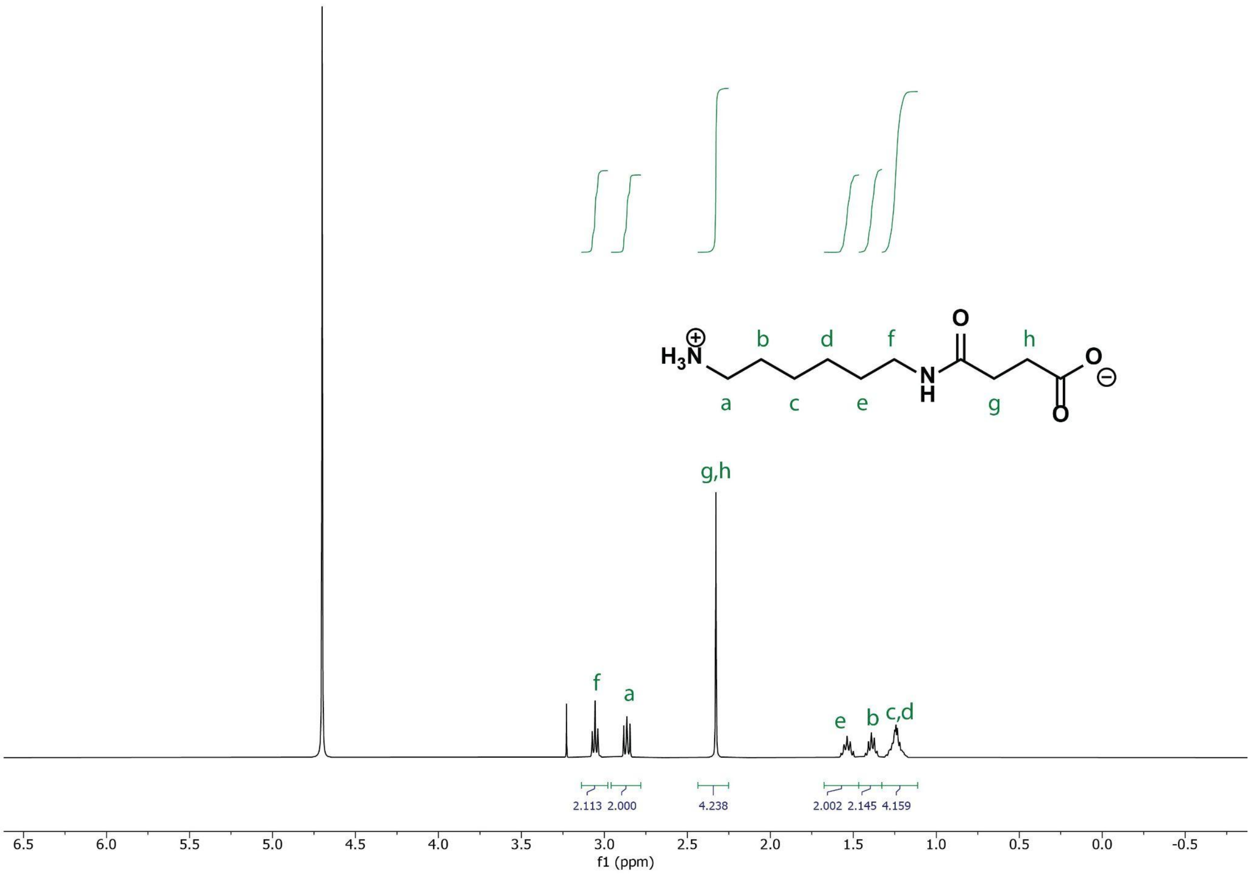


**Figure S36: $^1$H NMR spectrum of MS in $D_2O$.**

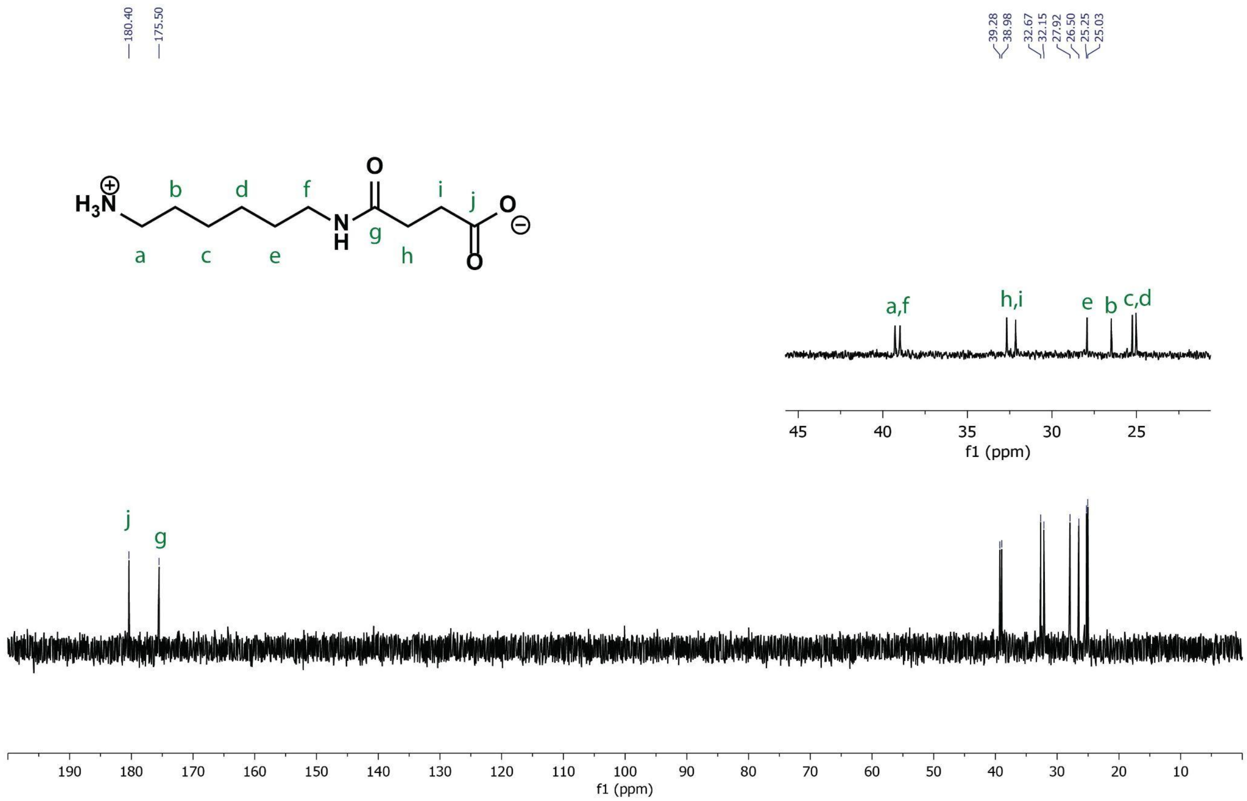


**Figure S37: $^{13}$C NMR spectrum of MS in $D_2O$.**

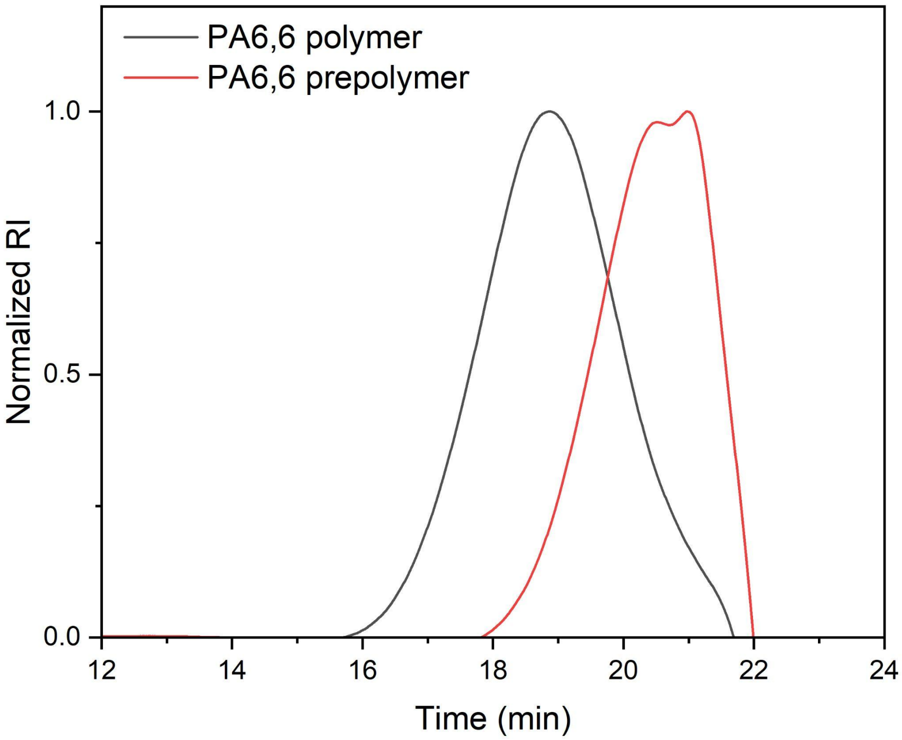

**Figure S38: SEC chromatogram of PA6,6 and prepolymer prepared from MA TFA salt.**

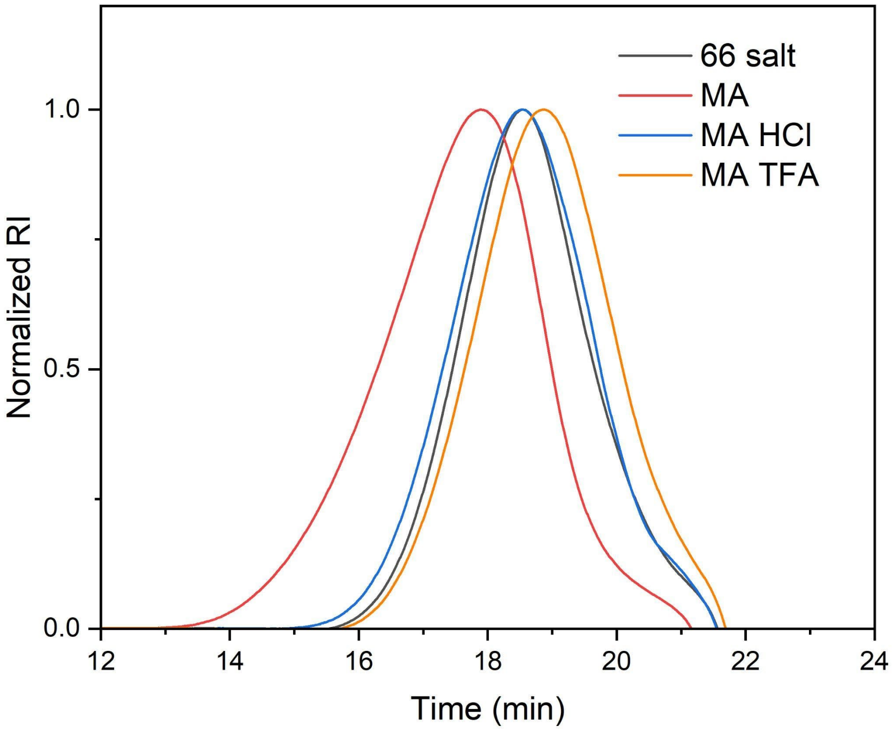


**Figure S39: SEC chromatograms of PA6,6 synthesized from different starting materials.** MA HCl and MA TFA reach lower molecular weight than freebased MA when polymerized under the same conditions.

**Table S1: Molecular weight data of PA6,6 synthesized from MA HCl and MA TFA.**

| Starting material | $M_n$ (kDa) | $M_w$ (kDa) | *Đ* |
|---|---:|---:|---:|
| MA TFA prepolymer | 0.47 | 1.5 | 3.11 |
| MA TFA | 2.3 | 9.0 | 3.89 |
| MA HCl | 3.4 | 12.8 | 3.78 |

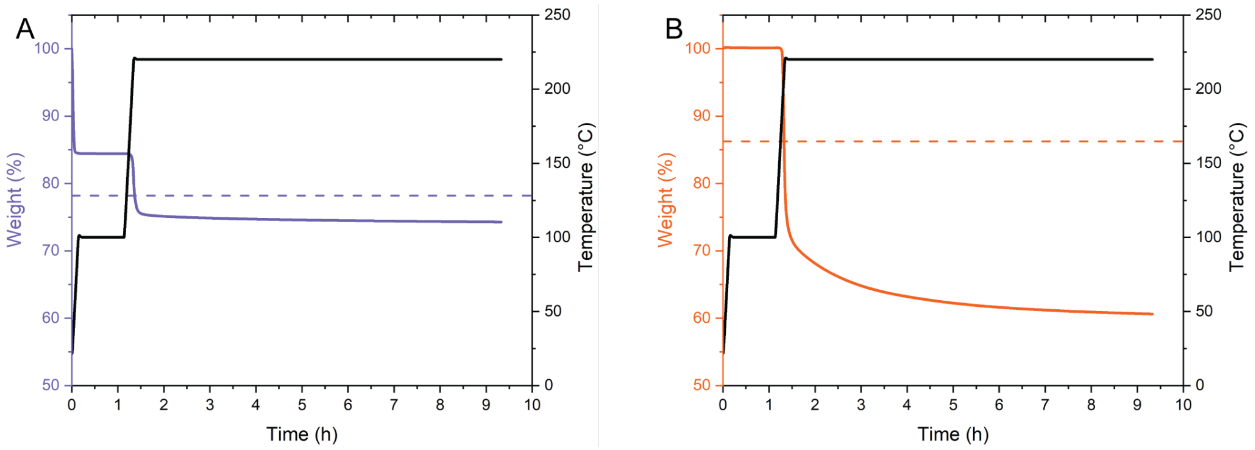


**Figure S40: Mass loss of PA6,6 salt (A) and diad (B) as measured via isothermal TGA. Samples were conditioned at 100 °C to remove any residual moisture or solvent, and subsequently polymerized at 220 °C for 8 h. Dashed line represents the maximum theoretical mass loss assuming 100% conversion of end groups with only the loss of water occurring.**

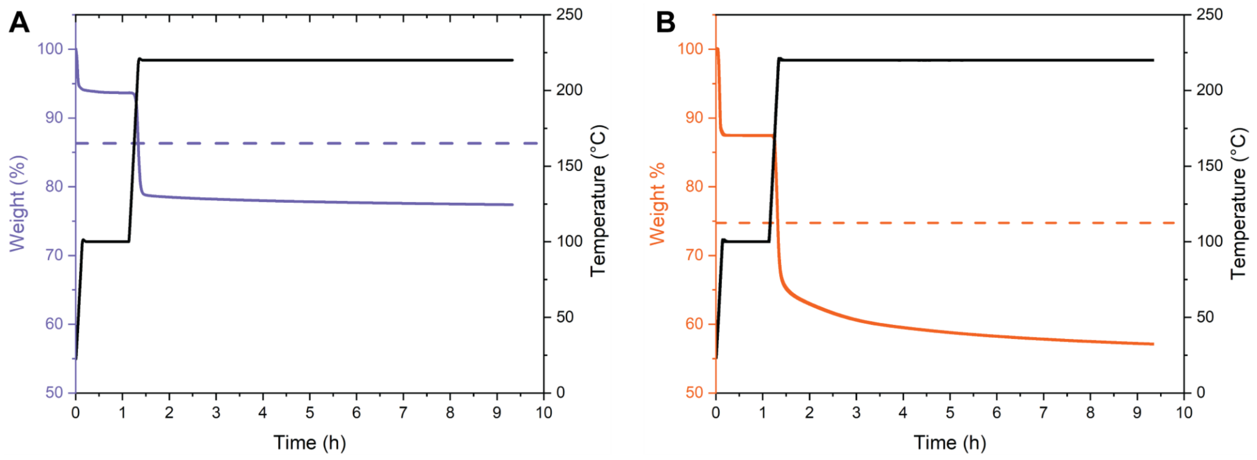


**Figure S41: Mass loss of PA5,6 salt (A) and diad (B) as measured via isothermal TGA. Dashed line represents the maximum theoretical mass loss assuming 100% conversion of end groups with only the loss of water occurring.**

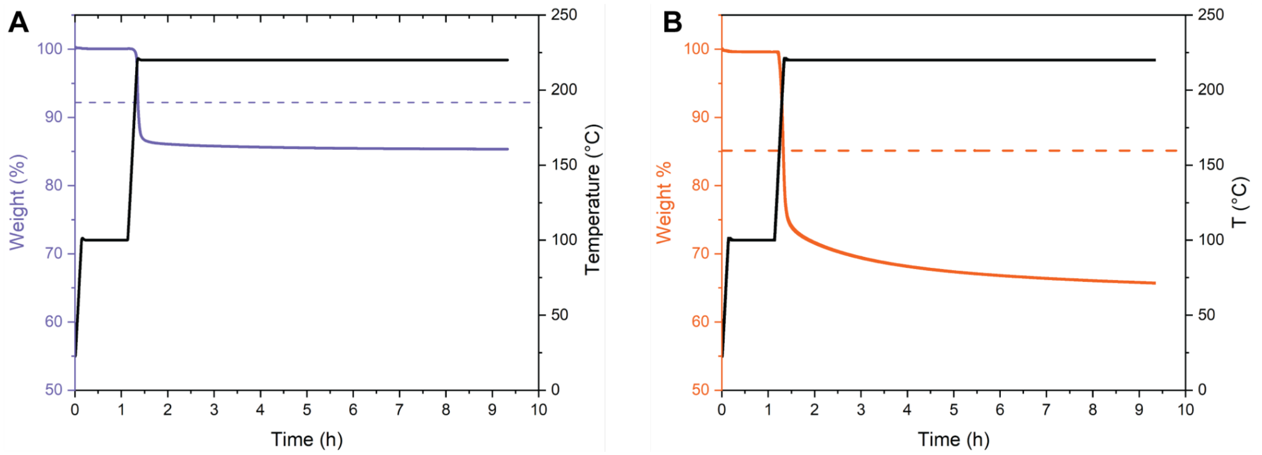


**Figure S42: Mass loss of PA6,5 salt (A) and diad (B) as measured via isothermal TGA. Dashed line represents the maximum theoretical mass loss assuming 100% conversion of end groups with only the loss of water occurring.**

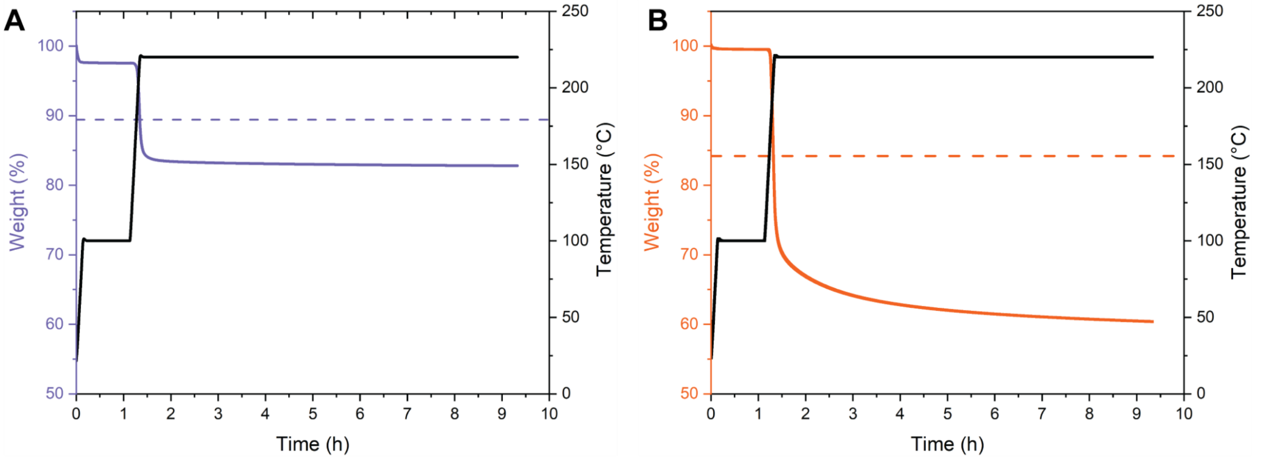


**Figure S43: Mass loss of PA5,5 salt (A) and diad (B) as measured via isothermal TGA. Dashed line represents the maximum theoretical mass loss assuming 100% conversion of end groups with only the loss of water occurring.**

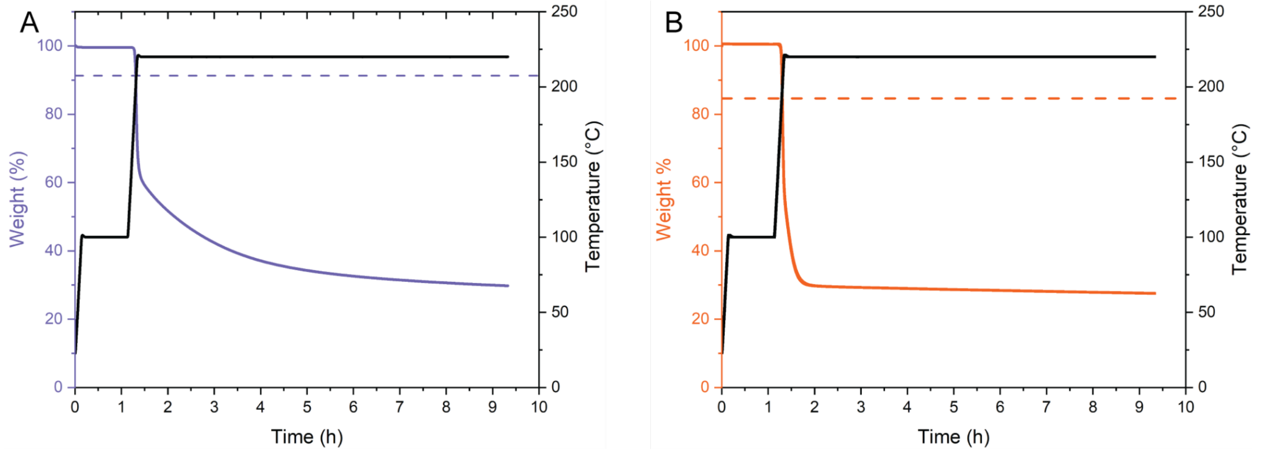

**Figure S44: Mass loss of PA6,4 salt (A) and diad (B) as measured via isothermal TGA. Dashed line represents the maximum theoretical mass loss assuming 100% conversion of end groups with only the loss of water occurring.**

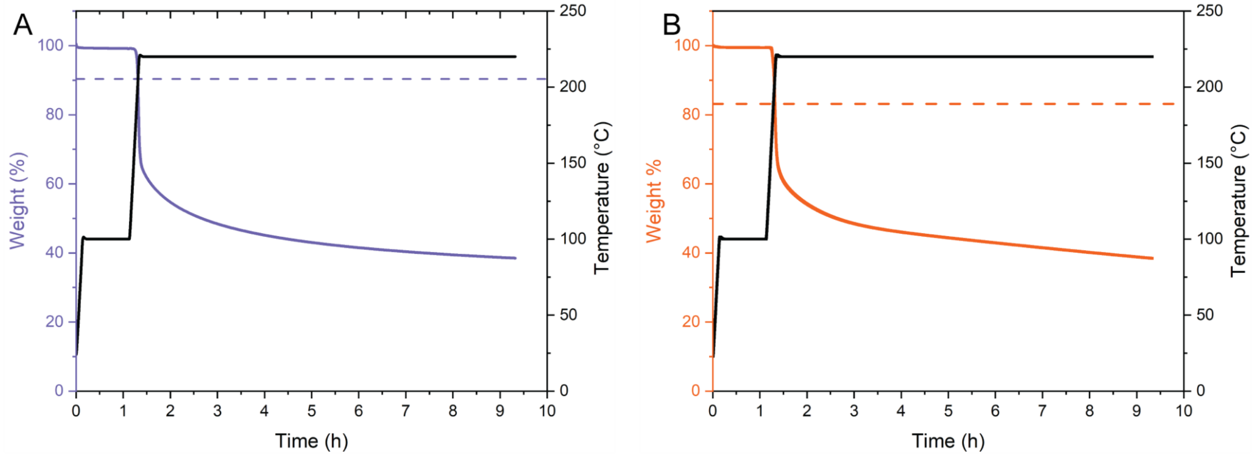


**Figure S45: Mass loss of PA5,4 salt (A) and diad (B) as measured via isothermal TGA. Dashed line represents the maximum theoretical mass loss assuming 100% conversion of end groups with only the loss of water occurring.**

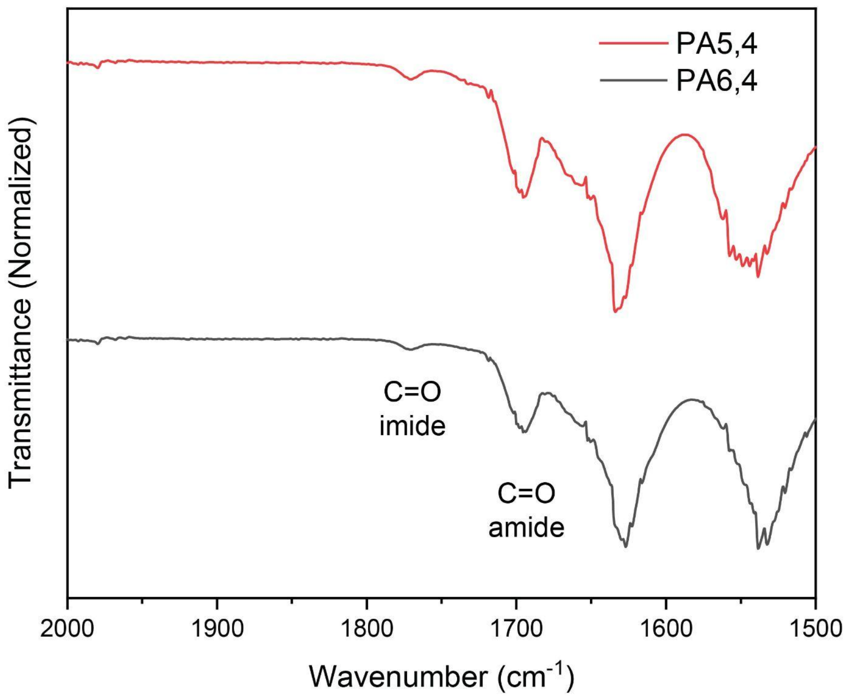


**Figure S46: Normalized FTIR spectra of PA5,4 and PA6,4.**

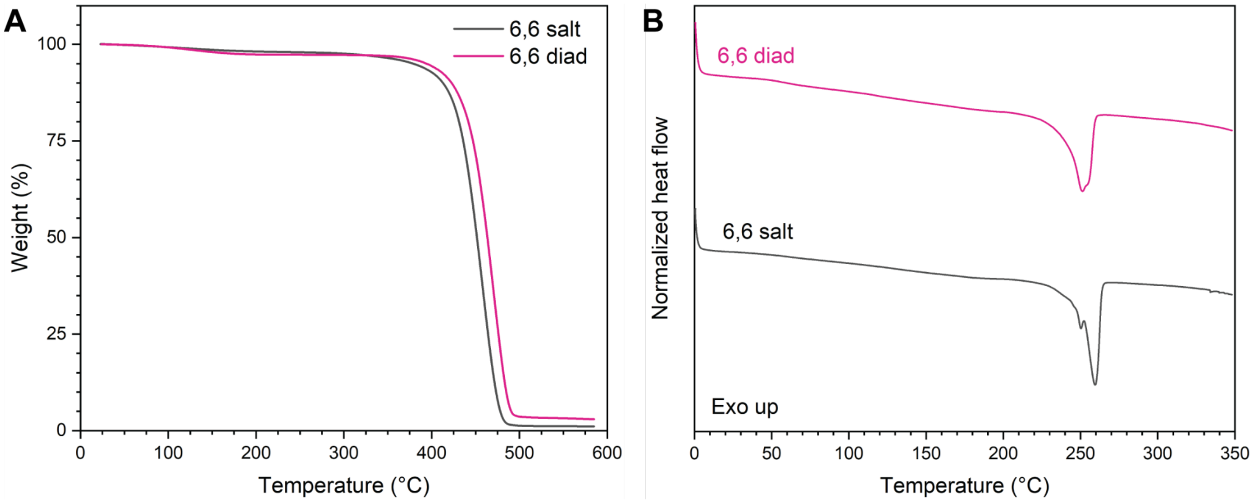


**Figure S47: TGA (A) and DSC (B; 1st heat cycle) thermograms of PA6,6 synthesized from MA and from PA6,6 salt.**

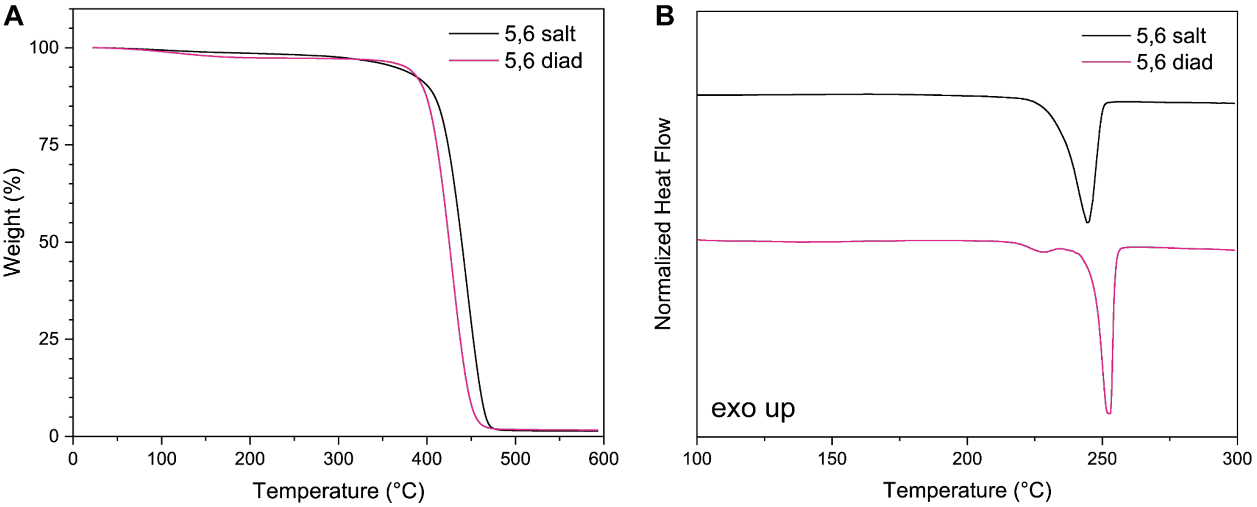


**Figure S48: TGA (A) and DSC (B; 1st heat cycle) thermograms of PA5,6 synthesized from CA and from PA5,6 salt.**

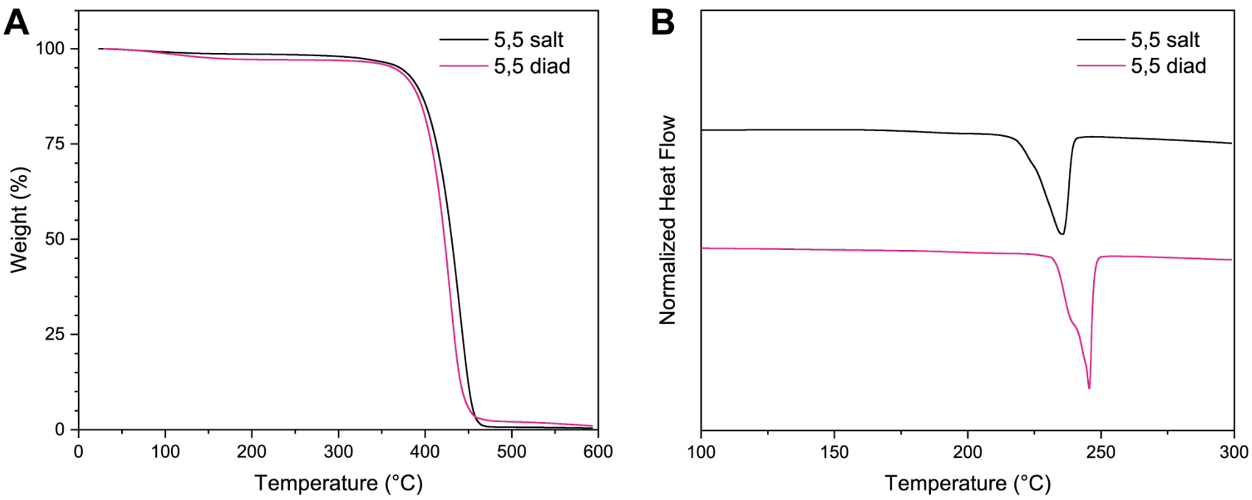


**Figure S49: TGA (A) and DSC (B; 1st heat cycle) thermograms of PA5,5 synthesized from MA and from PA5,5 salt.**

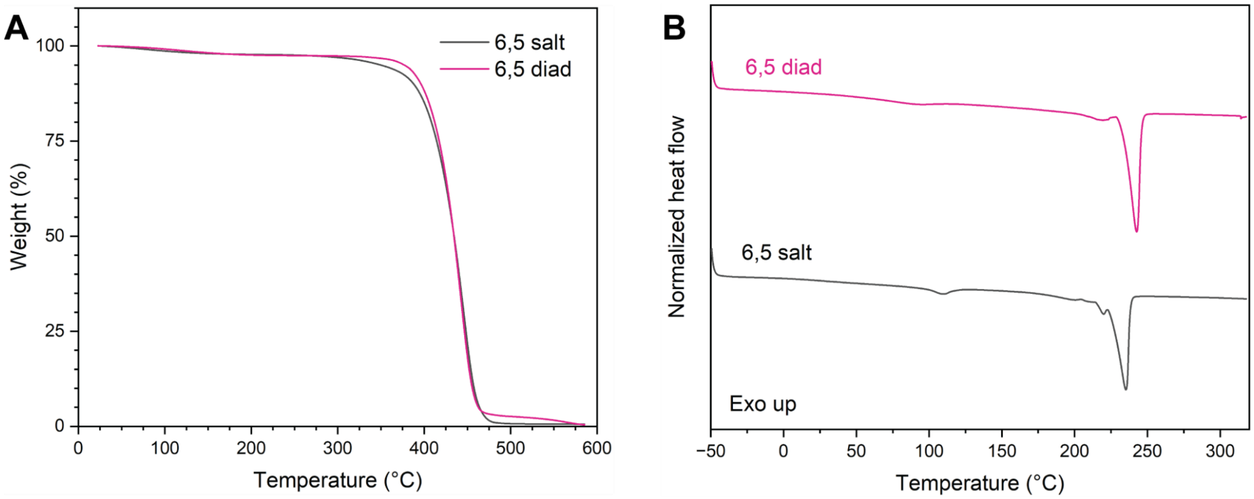


**Figure S50: TGA (A) and DSC (B; 1st heat cycle) thermograms of PA6,5 synthesized from MG and from PA6,5 salt.**

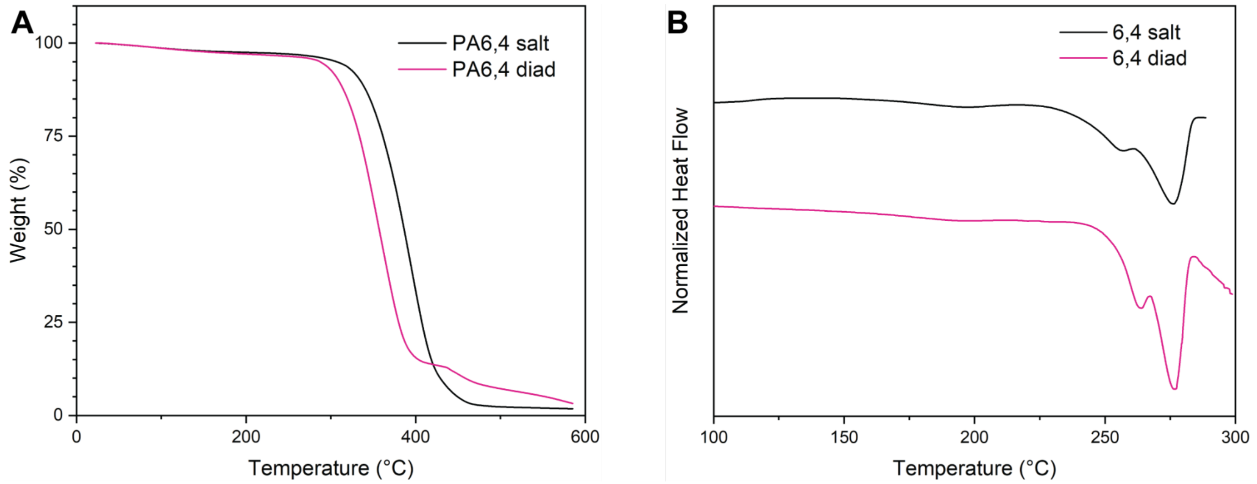


**Figure S51: TGA (A) and DSC (B; 1st heat cycle) thermograms of PA6,4 synthesized from MS and from PA6,4 salt.**

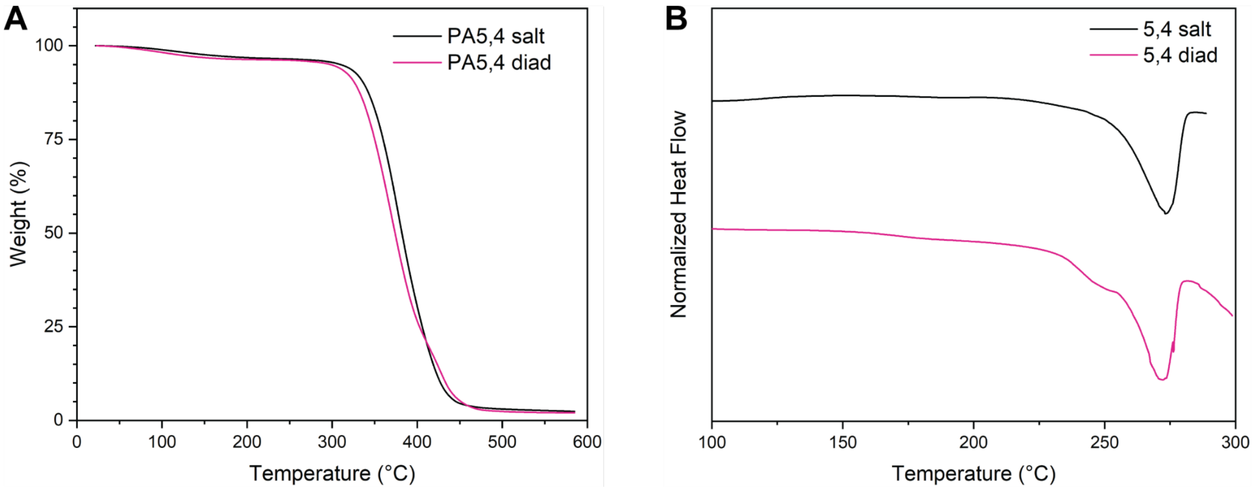


**Figure S52: TGA (A) and DSC (B; 1st heat cycle) thermograms of PA5,4 synthesized from CS and from PA5,4 salt.**

**Table S2: Thermal characterization data for all polymers.**

| Polymer | Starting Material | $T_d$ (°C) | $T_g$ (°C) | $T_{m,1}$ (°C) | $T_{m,2}$ (°C) | $\Delta H_m$ (J $g^{-1}$) |
|---|---|---|---|---|---|---|
| PA6,6 | 6,6 salt | 380 | 60 | 255 | n/a | 102.9 |
| | MA | 394 | 57 | 261 | n/a | 112.8 |
| PA5,6 | 6,5 salt | 364 | 66 | 244 | n/a | 96.7 |
| | CA | 376 | 65 | 253 | n/a | 85.5 |
| PA6,5 | 6,5 salt | 349 | 105 | 235 | 220 | 82.0 |
| | MG | 377 | 77 | 243 | 218 | 91.0 |
| PA5,5 | 5,5 salt | 371 | 43 | 238 | n/a | 92.6 |
| | CG | | 56 | 245 | n/a | 86.9 |
| PA6,4 | 6,4 salt | 309 | | 276 | 237 | 92.1 |
| | MS | 293 | | 276 | 261 | 113.6 |
| PA5,4 | 5,4 salt | 309 | | 273 | n/a | 73.8 |
| | CS | 297 | | 272 | n/a | 76.9 |

**Supplementary methods for enzymatic reactions:**

*Scale-up of enzymatic synthesis optimization*: We first examined the effect on yield of increasing the substrate loading to 25 mM, 50 mM, or 100 mM (i.e. 5- to 20-fold increases) and varying the polyphosphate concentration. In general, the percentage yield decreased while titer increased as the substrate load was increased from 25 mM to 100 mM (Figure S61). Additionally, increasing polyphosphate concentrations led to higher production, particularly at high substrate loadings. To maximize titer while maintaining a relatively high yield, we selected 50mM substrate loadings and then investigated the effects of $MgCl_2$, AMP, different diacid/diamine ratios, and pH on system performance.

Notably, $MgCl_2$ concentration significantly impacted production, with a marked increase observed from 10 mM to 50 mM, and stabilization with further increases to 100 mM (Figure S62). Since polyP could chelate magnesium ions, potentially affecting production, 100 mM of $Mg^{2+}$ was selected for the subsequent large-scale reaction. We also noticed a slight increase in production when AMP concentration was raised from 1 mM to 5 mM (Figure S63). Interestingly, increasing the diamine to diacid ratio from 1:1 to 2:1 enhanced MS production, suggesting that an excess of amine may drive the reaction equilibrium forward (Figure S64). However, further increases did not result in additional gains, potentially due to enzyme inhibition by high amine concentrations. pH had a modest impact on reaction performance, with pH 8 remaining the optimal condition for the overall system (Figure S65).

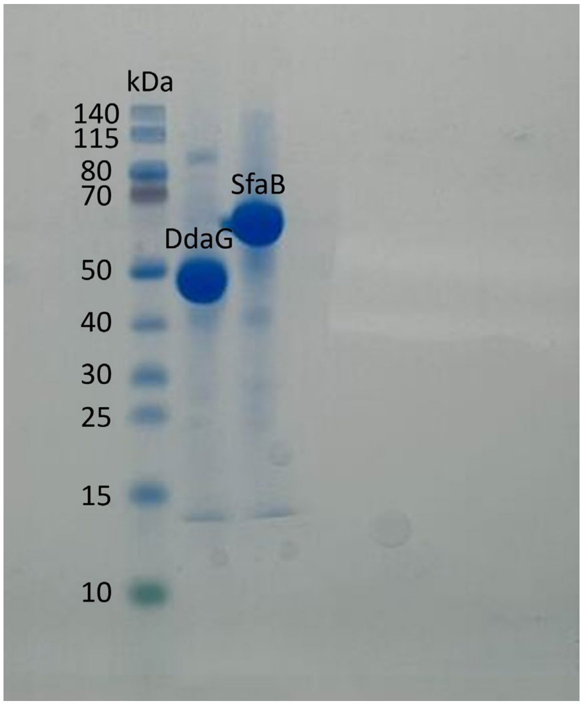


**Figure S53: SDS-PAGE analysis of recombinant *N*-terminal His-tagged proteins purified from *E.coli* BL21(DE3)**. Novex Tris-Glycine gels (4%-12%, Invitrogen) were used. DdaG expected size: 45.58 kDa; SfaB expected size: 65.06 kDa.

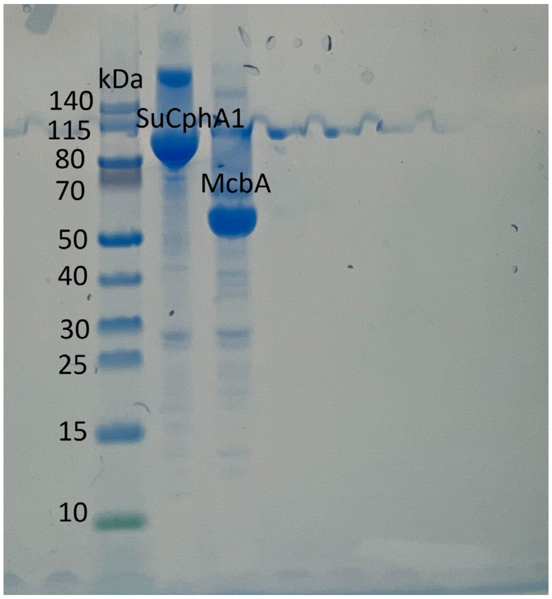


**Figure S54: SDS-PAGE analysis of recombinant *N*-terminal His-tagged proteins purified from *E.coli* BL21(DE3)**. Novex Tris-Glycine gels (4%-12%, Invitrogen) were used. SuCphA1 expected size: 94.85 kDa; McbA expected size: 55.02 kDa.

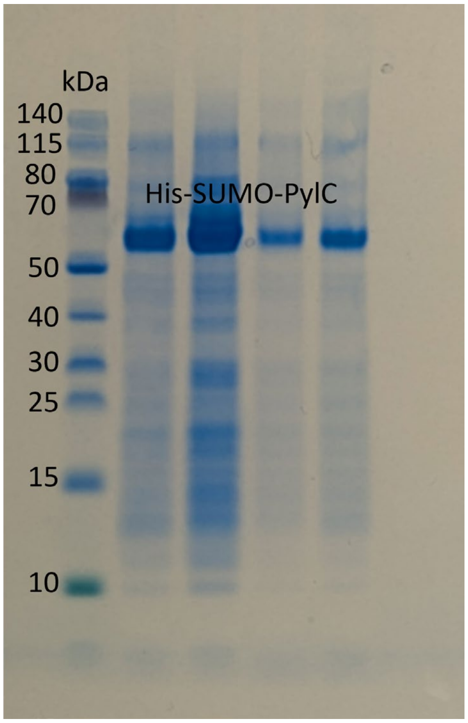


**Figure S55: SDS-PAGE analysis of recombinant *N*-terminal His-tagged proteins purified from *E.coli* BL21(DE3)**. Novex Tris-Glycine gels (4%-12%, Invitrogen) were used. His-SUMO-PylC expected size: 54.50 kDa. Four aliquots of His-SUMO-PylC were included in the gel.

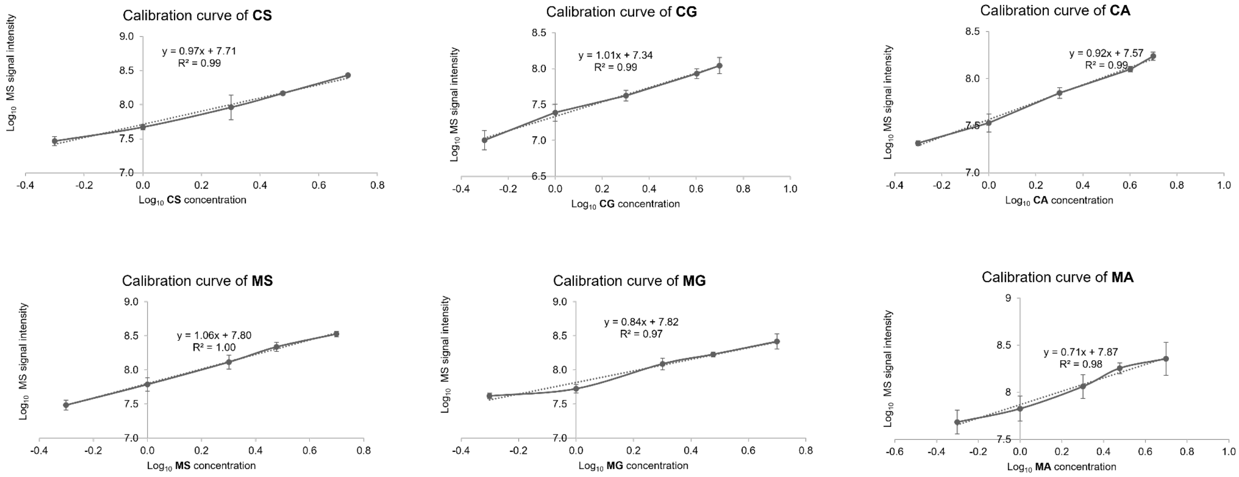


**Figure S56: Calibration curves of CS, MS, CG, MG, CA and MA**. Chemically-synthesized **CS, MS, CG, MG, CA** and **MA** were diluted in reaction buffer without enzymes at relevant concentrations (i.e., 0.5, 1, 2, 3 or 4, and 5 mM) to generate calibration curves for IDOT/OPSI-MS analysis. Sample size is $N = 3$. Error bars are shown as the mean ± standard deviation.

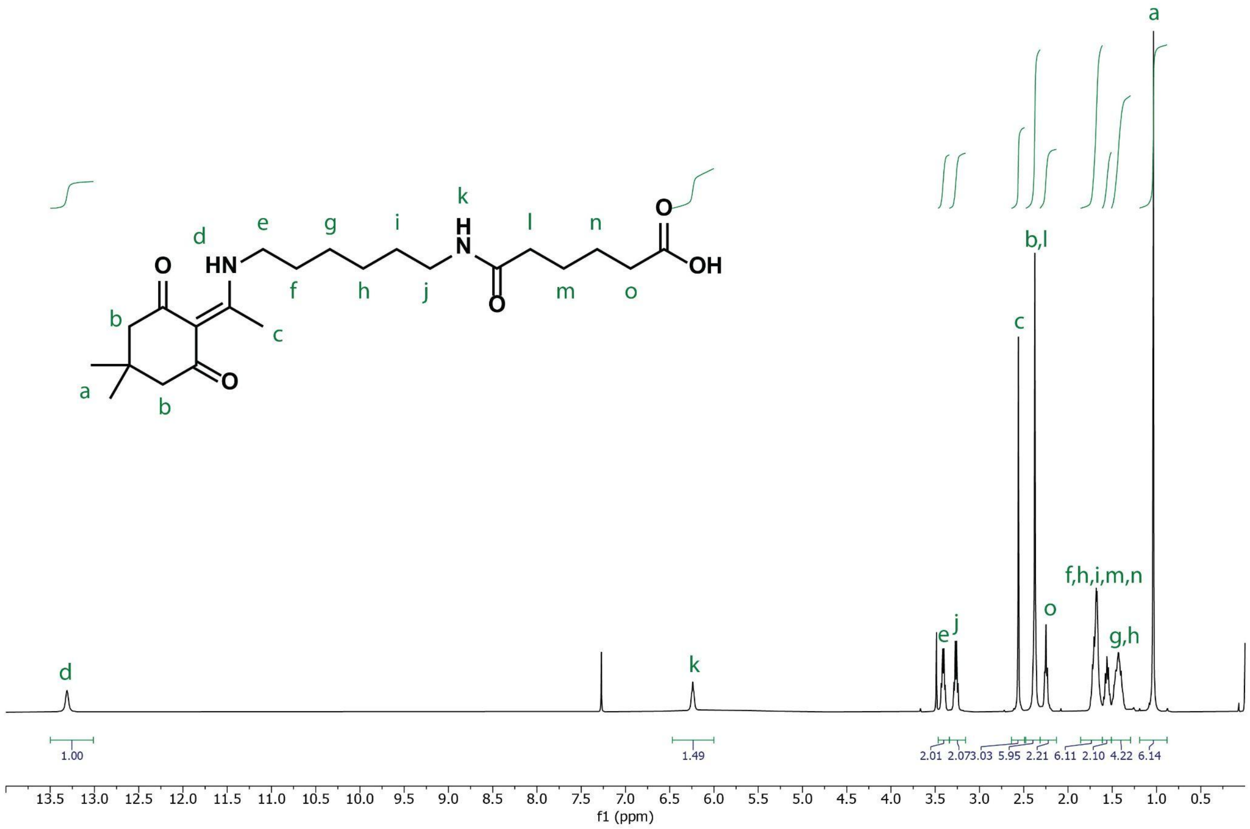


**Figure S57: $^1$H NMR spectrum of dde-functionalized MA in $CDCl_3$.**

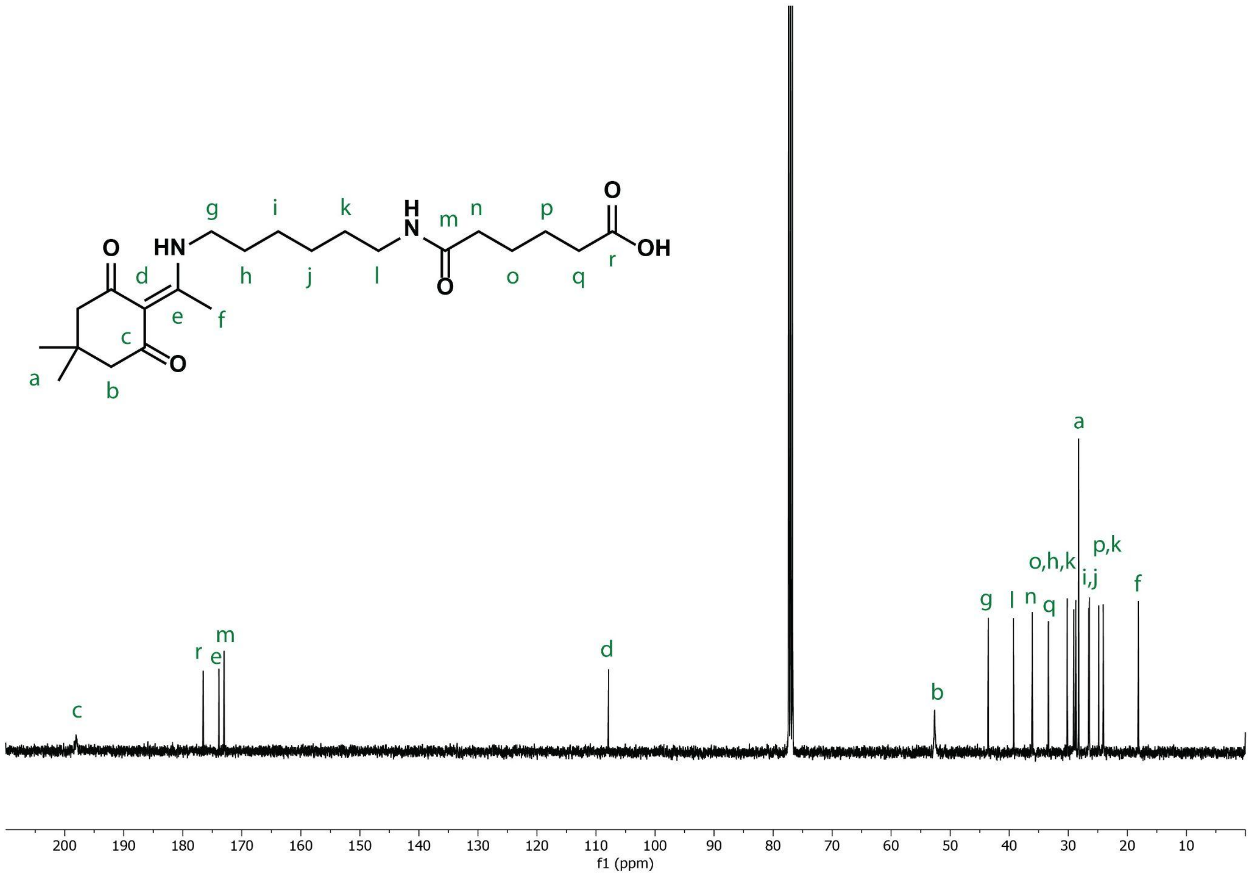


**Figure S58: $^{13}C$ NMR spectrum of dde-functionalized MS in $CDCl_3$.**

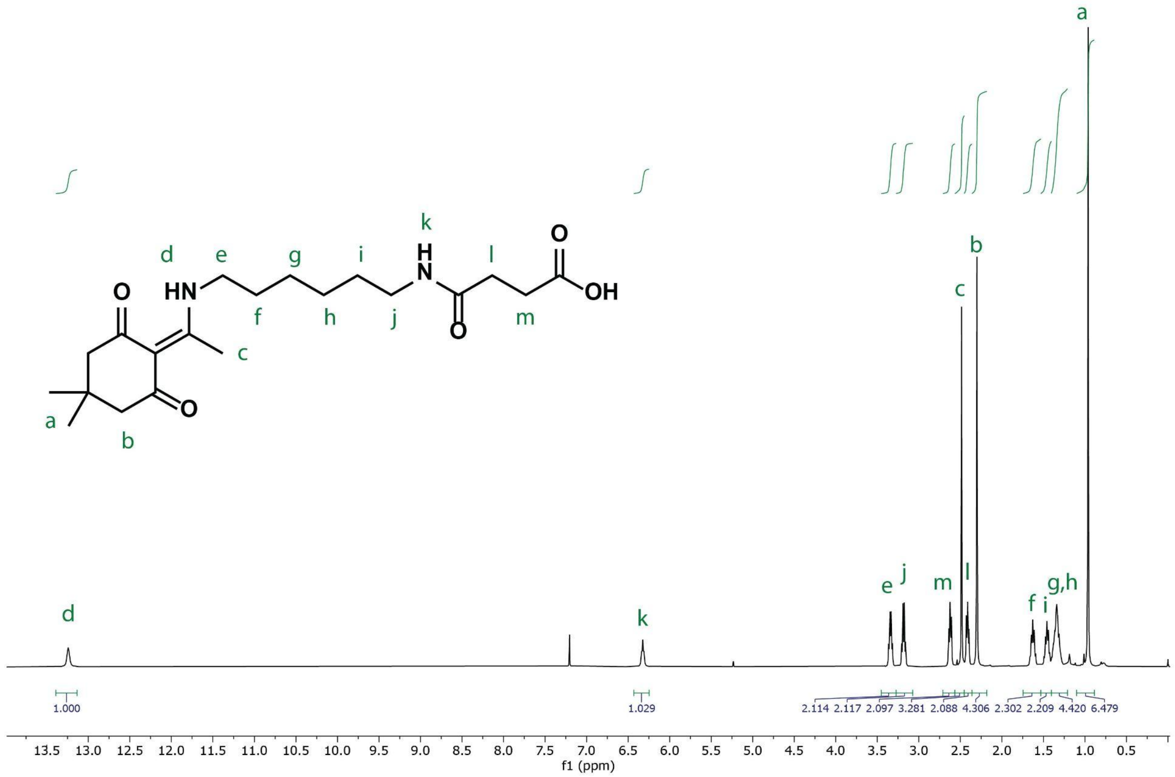


**Figure S59: $^{1}$H NMR spectrum of dde-functionalized MS in $CDCl_3$.**

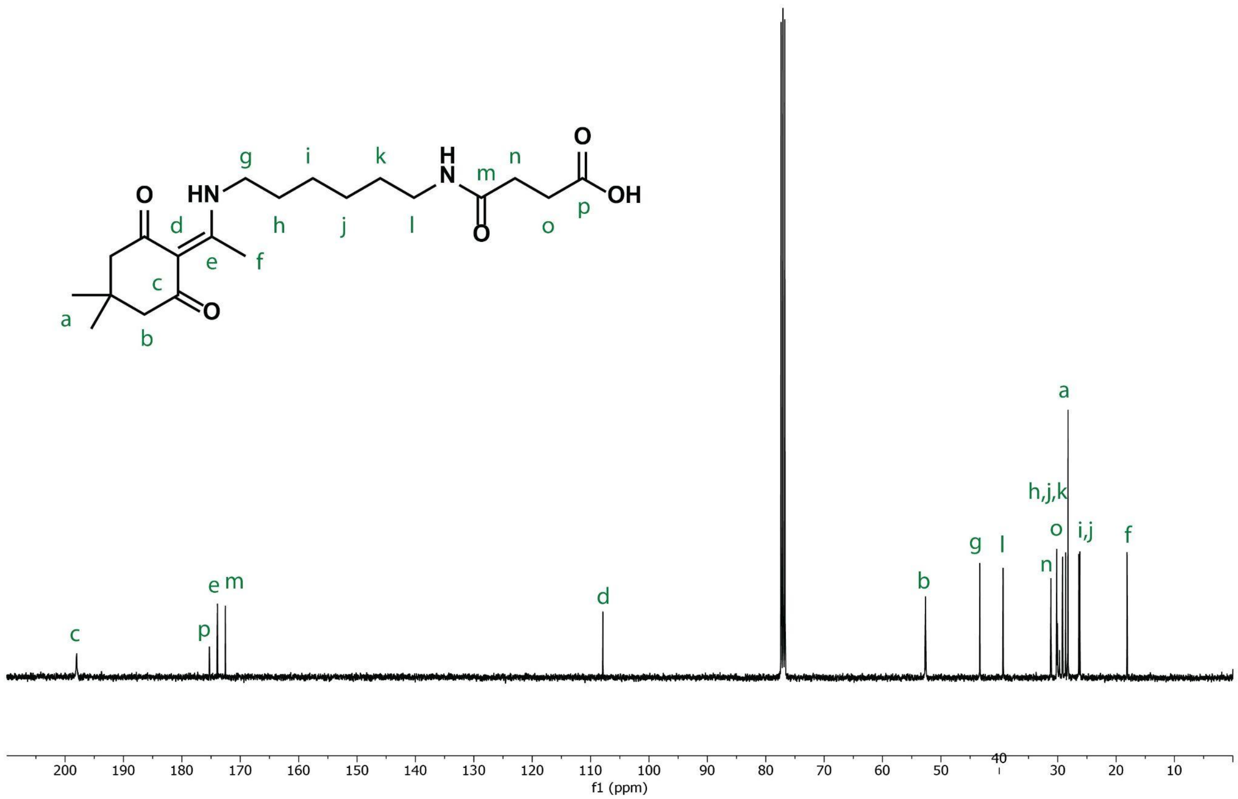


**Figure S60: [13]C NMR spectrum of dde-functionalized MS in CDCl₃.**

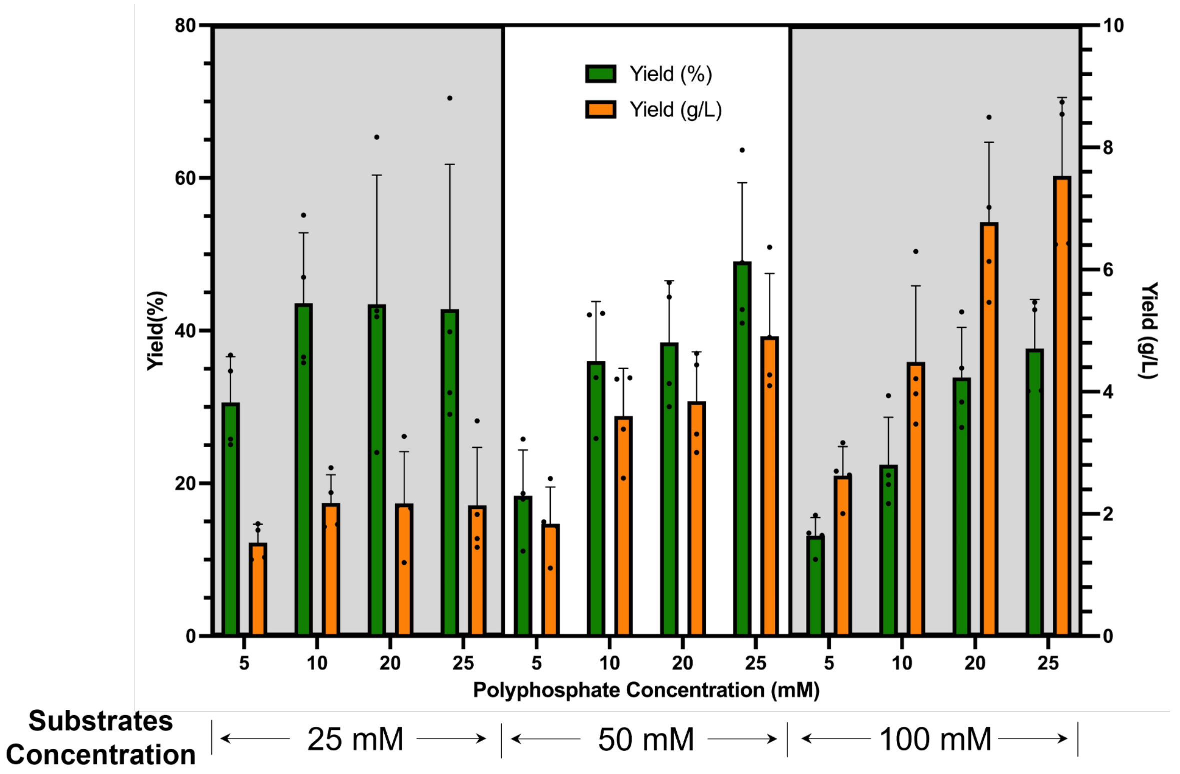


**Figure S61: Effect of substrate and polyphosphate concentration on MS synthesis**. Assays were conducted by incubating varied substrate concentrations and different polyphosphate concentrations with DdaG coupled with ATP-regeneration systems. Reactions were performed in 100 mM HEPES pH 8, 1 mM ATP, 1 mM AMP, 40 mM $MgCl_2$, 5-25 mM SHMP, 25-100 mM disodium succinic acid, 25-100 mM hexamethylene diamine, 10 µM DdaG, 0.2 U ecPPase, and 1 mg/mL PPK12. Products were assayed using OPSI-MS and the yields were calculated by referring to the calibration curve. Sample size is $N$ = 4. Error bars are shown as the mean ± standard deviation.

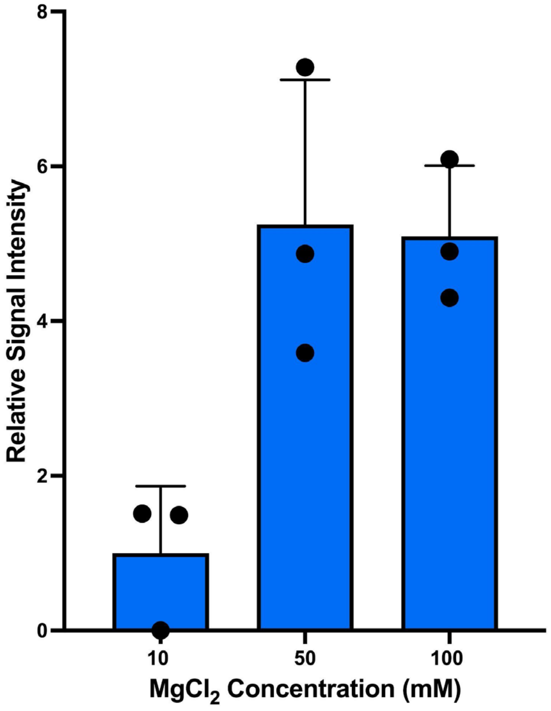


**Figure S62: Effect of $MgCl_2$ concentration on MS synthesis**. Assays were conducted by incubating different $MgCl_2$ concentrations with DdaG coupled with ATP-regeneration systems. Reactions were performed in 100 mM HEPES pH 8, 1 mM ATP, 1 mM AMP, 10-100 mM $MgCl_2$, 25 mM SHMP, 50 mM disodium succinic acid, 50 mM hexamethylenediamine,10 µM DdaG, 0.2 U ecPPase, and 1 mg/mL PPK12. Products were assayed using OPSI-MS and the relative signal intensity were calculated by referring to the mean of the group with 10 mM $MgCl_2$. Sample size is $N = 3$. Error bars are shown as the mean ± standard deviation.

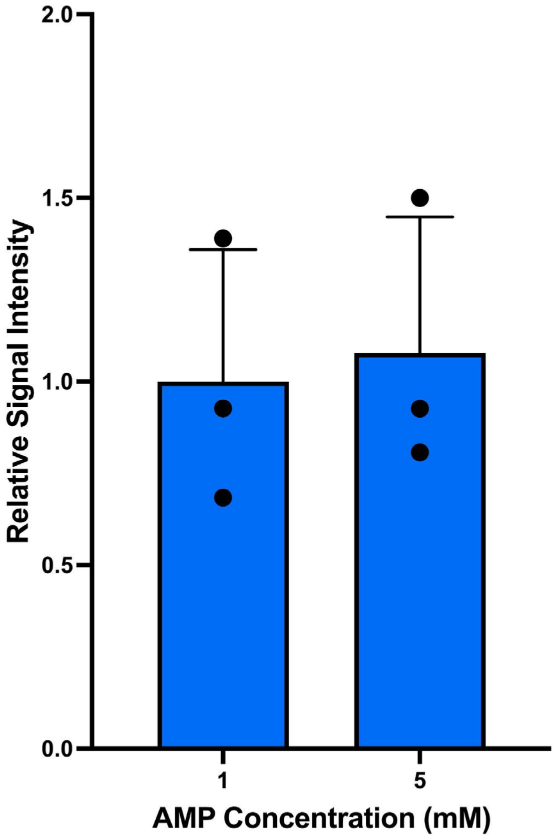


**Figure S63: Investigation of AMP concentration on MS production**. Assays were conducted by incubating different AMP concentrations with DdaG coupled with ATP-regeneration systems. Reactions were performed in 100 mM HEPES pH 8, 1 mM ATP, 1-5 mM AMP, 100 mM $MgCl_2$, 25 mM SHMP, 50 mM disodium succinic acid, 50 mM hexamethylenediamine,10 µM DdaG, 0.2 U ecPPase, and 1 mg/mL PPK12. Products were assayed using OPSI-MS and the relative signal intensity were calculated by referring to the mean of the group with 1 mM AMP. Sample size is *N* = 3. Error bars are shown as the mean ± standard deviation.

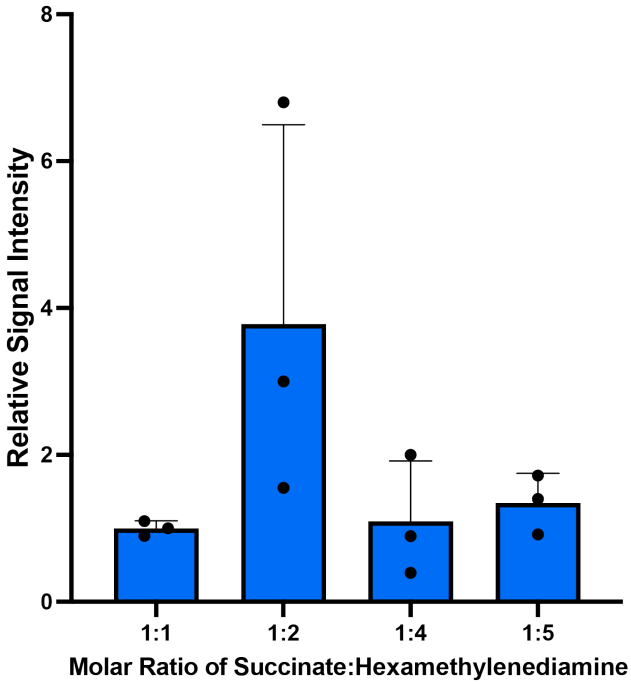


**Figure S64: Investigation of molar ratio of succinate to hexamethylenediamine on MS production**. Assays were conducted by incubating different molar ratio of succinate to hexamethylenediamine with DdaG coupled with ATP-regeneration systems. Reactions were performed in 100 mM HEPES pH 8, 1 mM ATP, 5 mM AMP, 100 mM $MgCl_2$, 25 mM SHMP, 50 mM disodium succinic acid, 50-250 mM hexamethylenediamine,10 µM DdaG, 0.2 U ecPPase, and 1 mg/mL PPK12. Products were assayed using OPSI-MS and the relative signal intensity were calculated by referring to the mean of the group with 1:1 as the molar ratio of succinate to hexamethylenediamine. Sample size is $N$ = 3. Error bars are shown as the mean ± standard deviation.

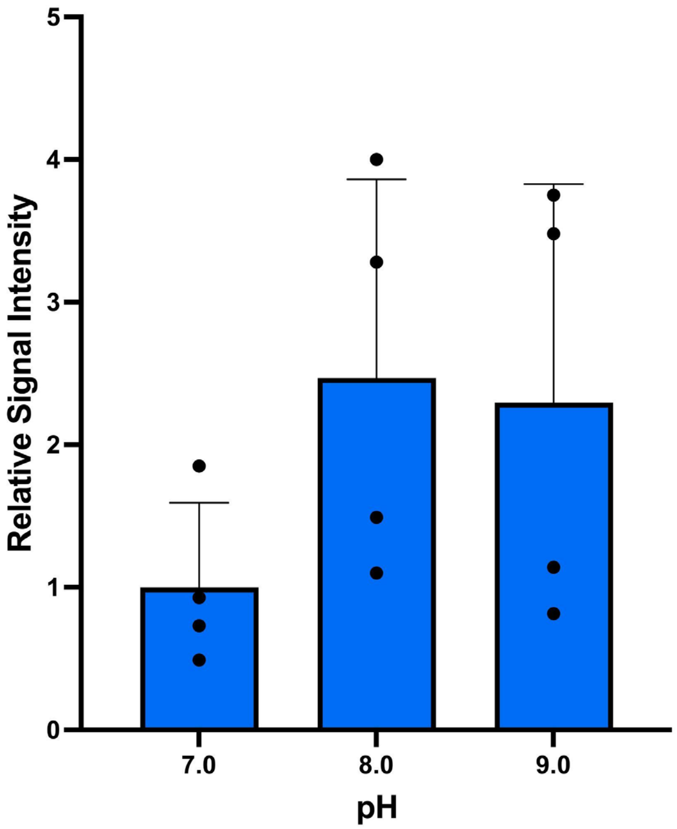


**Figure S65: Investigation of pH effect on MS production**. Assays were conducted by performing reactions in different pH of 100 mM HEPES buffers with DdaG coupled with ATP-regeneration systems. Reactions were performed in 100 mM HEPES pH 7.0-9.0, 1 mM ATP, 5 mM AMP, 100 mM $MgCl_2$, 25 mM SHMP, 50 mM disodium succinic acid, 100 mM hexamethylenediamine,10 µM DdaG, 0.2 U ecPPase, and 1 mg/mL PPK12. Products were assayed using OPSI-MS and the relative signal intensity were calculated by referring to the mean of the group with pH 7.0. Sample size is $N = 4$. Error bars are shown as the mean ± standard deviation.

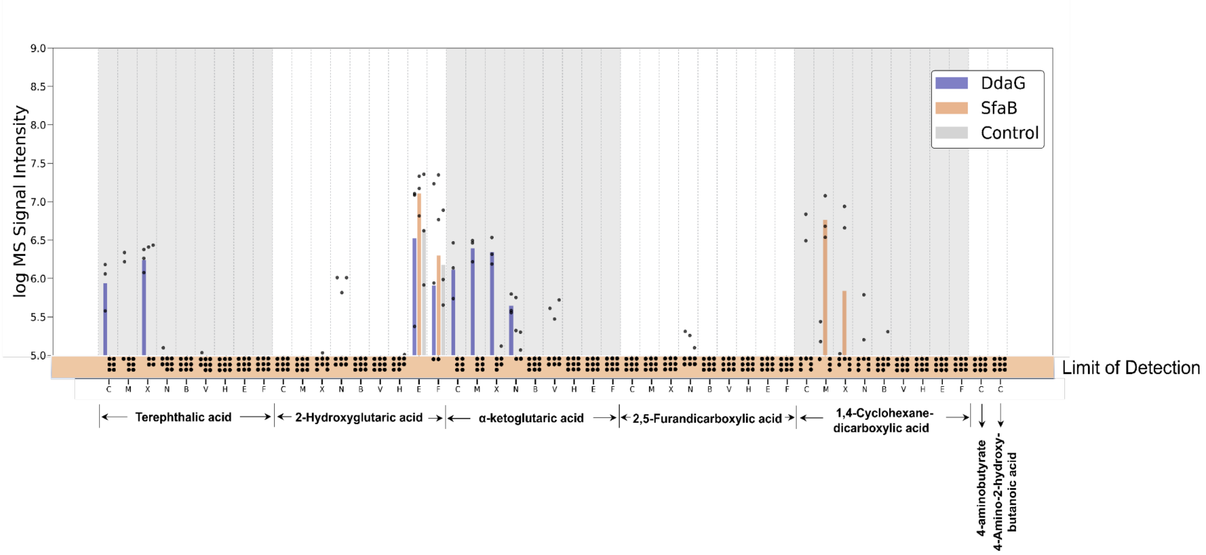


**Figure S66: Amide synthetases generate polymer-relevant diads**. *In vitro* biochemical assays were conducted by incubating 5 mM of different carboxylic acid-containing diacids or ω-amino acids (i.e., terephthalic acid, 2-hydroxyglutaric acid, α-ketoglutaric acid, 2,5-furandicarboxylic acid, 1,4-cyclohexanedicarboxylic acid, 4-aminobutyrate and 4-amino-2-hydroxybutanoic acid) with 5 mM diamine or ω-amino acids in the presence of 10 μM enzymes (i.e., DdaG or SfaB) or no-enzyme control at 30 °C for 16 h with shaking at 200 rpm. Products were assayed using OPSI-MS. The log MS signal intensity was cut off at 5.0 to filter out noise. **C**: cadaverine; **M**: hexamethylenediamine; **X**: *p*-Xylydiamine; **N**: *cis*-1,4-Cyclohexanediamine; **B**: 4-aminobutyrate; **V**: 5-aminovalerate; **H**: 6-aminohexanoic acid; **E**: 4-amino-2-hydroxybutanoic acid; **F**: 4-amino-3-hydroxybutanoic acid. Sample size is $N = 3$.

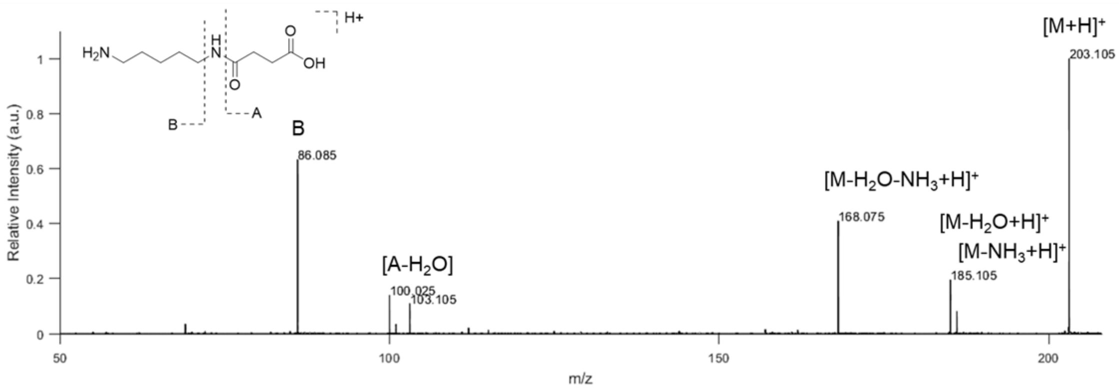


**Figure S67: I.DOT/OPSI-MS[2] of CS.**

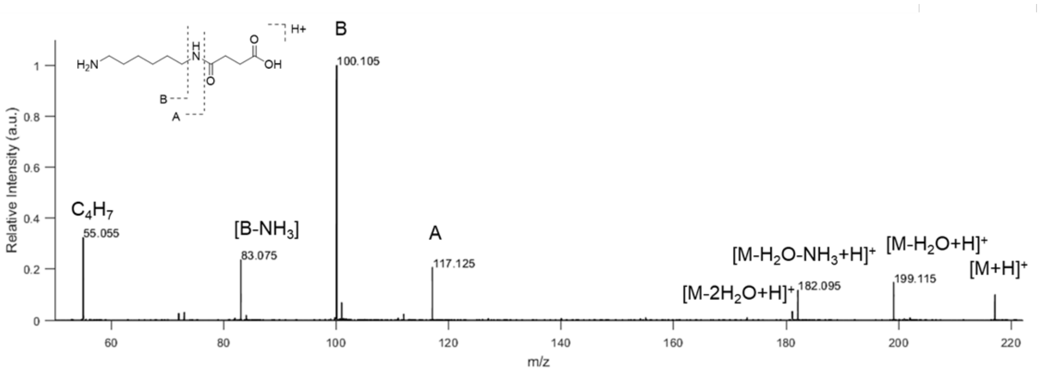


**Figure S68: I.DOT/OPSI-MS$^2$ of MS.**

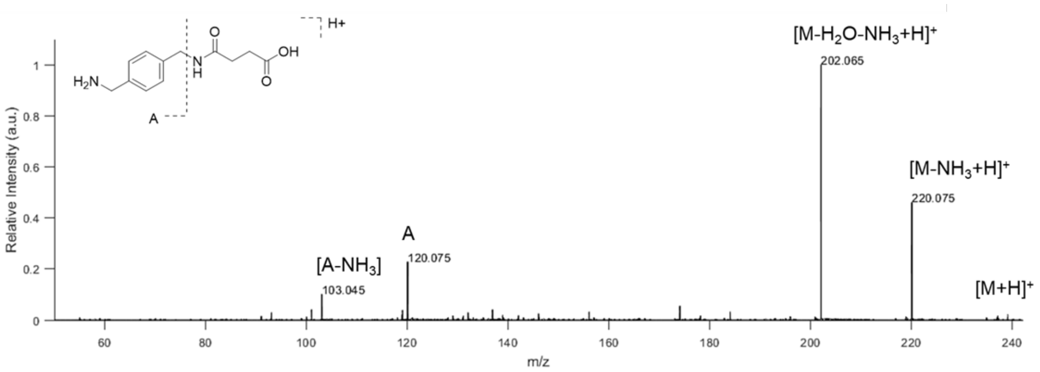


**Figure S69: I.DOT/OPSI-MS$^2$ of XS.**

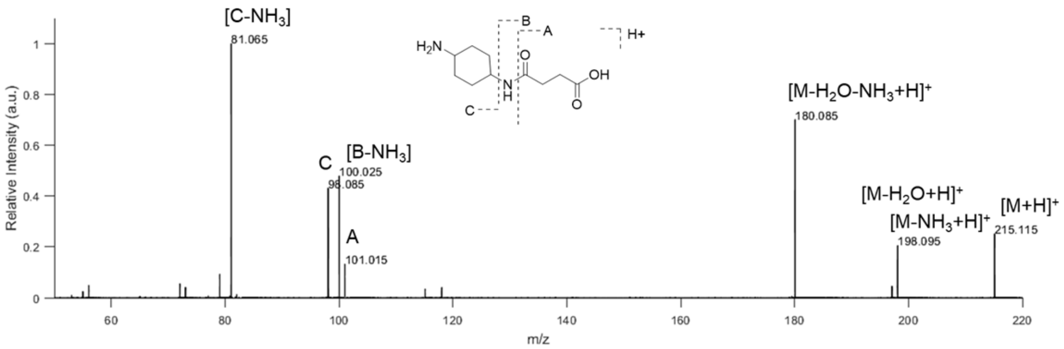


**Figure S70: I.DOT/OPSI-MS$^2$ of NS.**

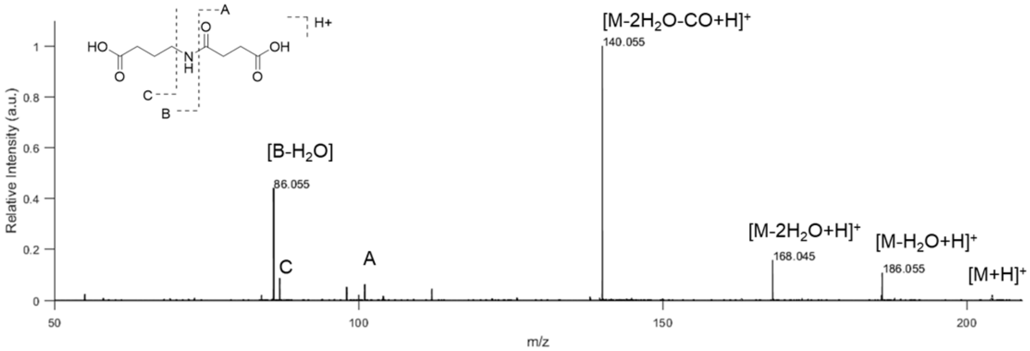


**Figure S71: I.DOT/OPSI-MS[2] of BS.**

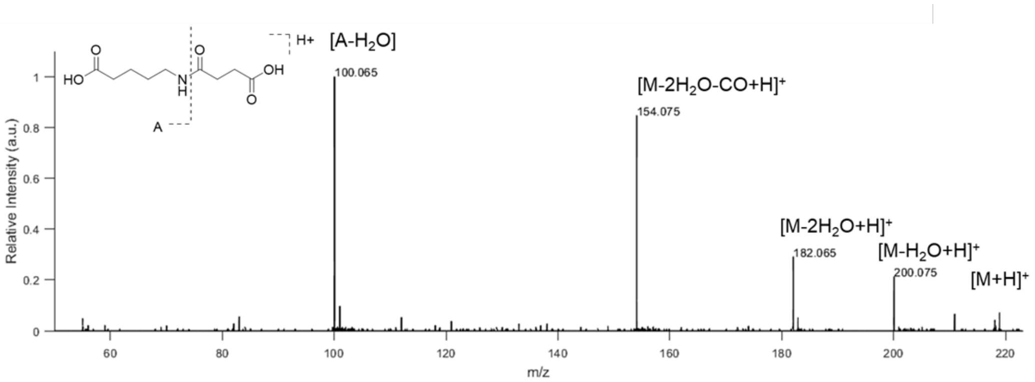


**Figure S72: I.DOT/OPSI-MS[2] of VS.**

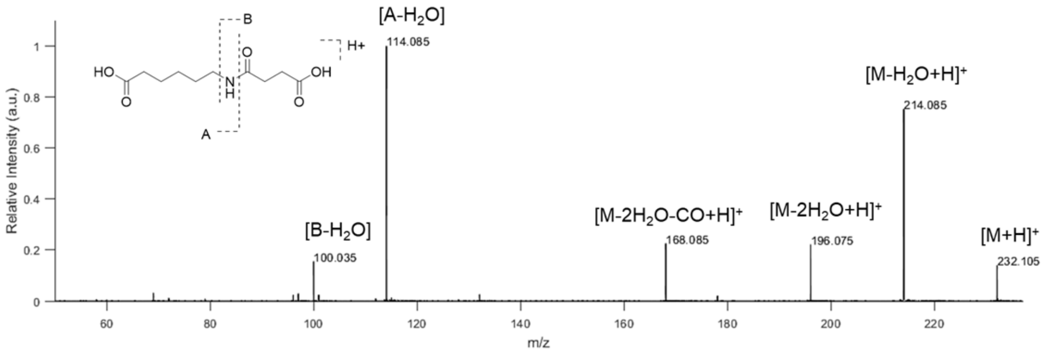


**Figure S73: I.DOT/OPSI-MS$^2$ of HS.**

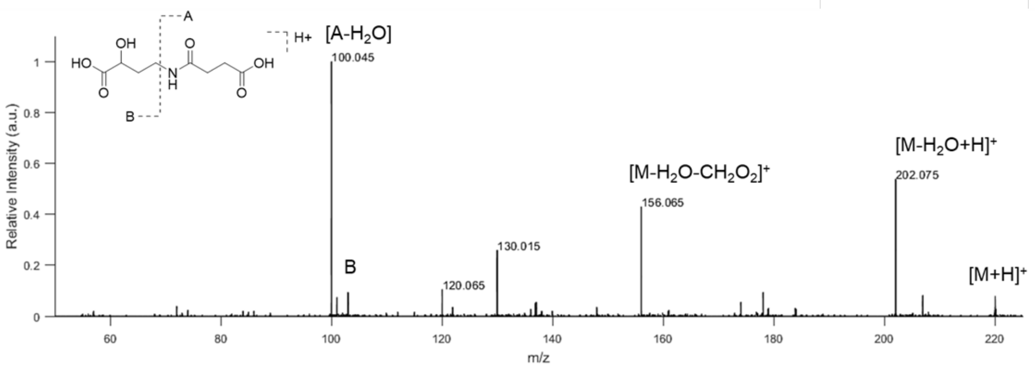


**Figure S74: I.DOT/OPSI-MS$^2$ of ES.**

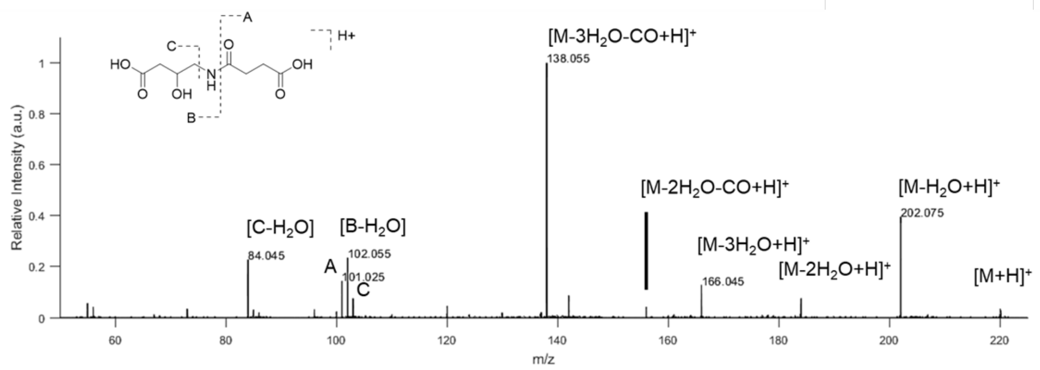


**Figure S75: I.DOT/OPSI-MS$^2$ of FS.**

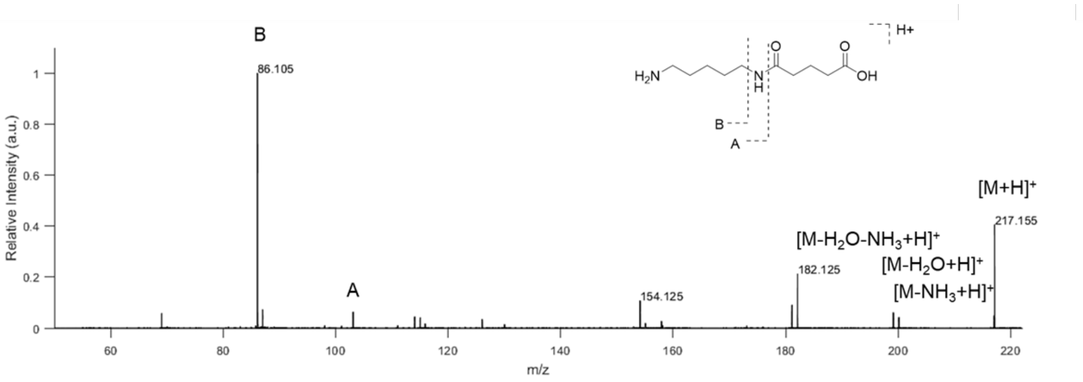


**Figure S76: I.DOT/OPSI-MS$^2$ of CG.**

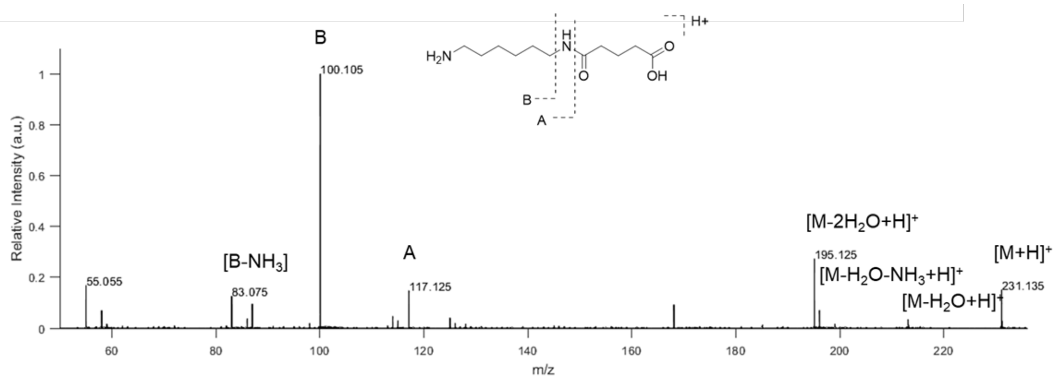


**Figure S77: I.DOT/OPSI-MS[2] of MG.**

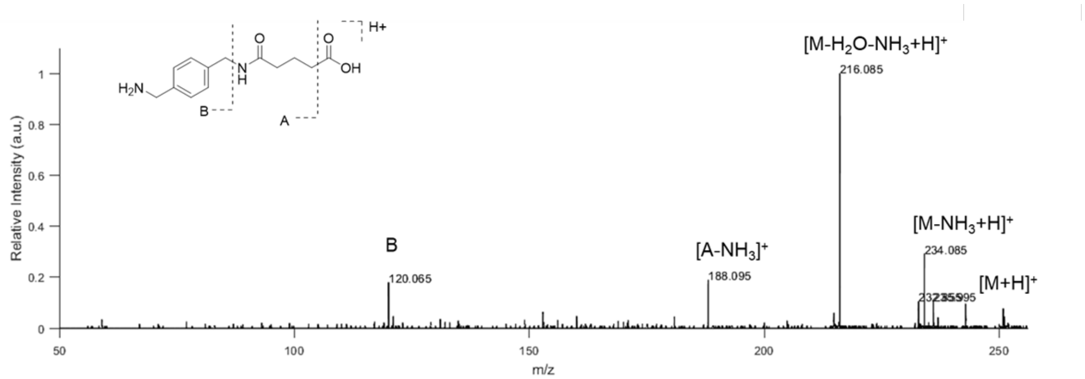


**Figure S78: I.DOT/OPSI-MS$^2$ of XG.**

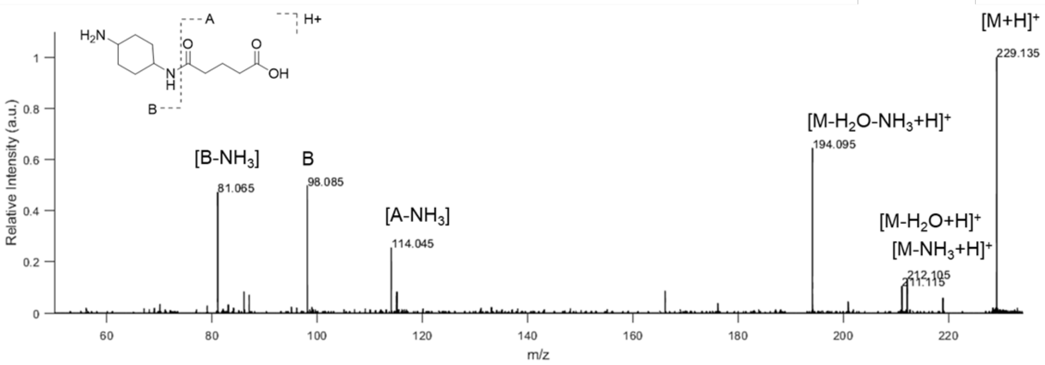


**Figure S79: I.DOT/OPSI-MS$^2$ of NG.**

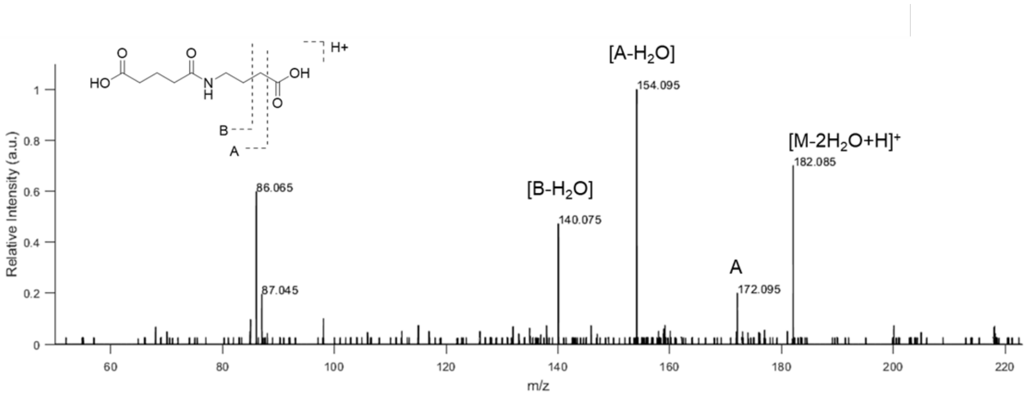


**Figure S80: I.DOT/OPSI-MS$^2$ of BG.**

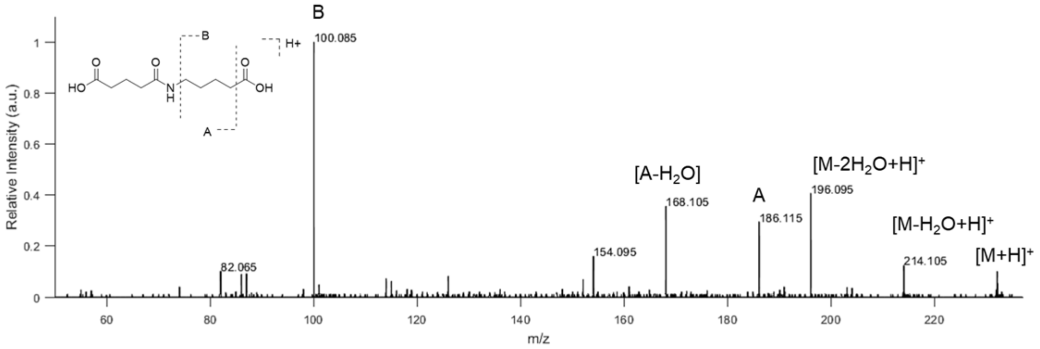


**Figure S81: I.DOT/OPSI-MS$^2$ of VG.**

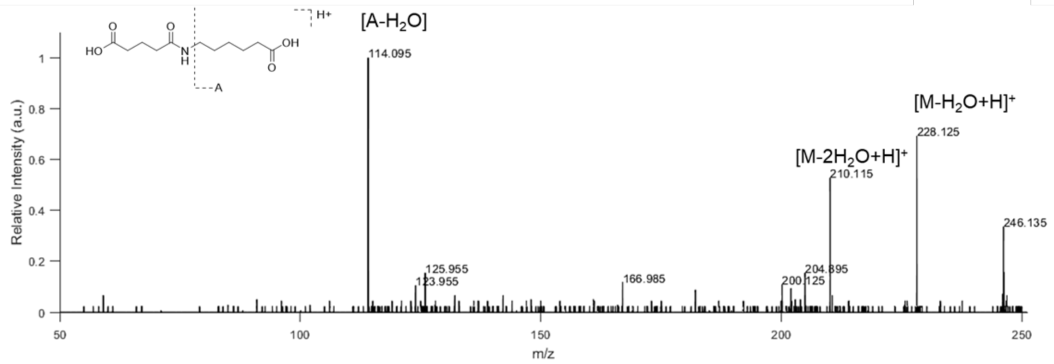


**Figure S82: I.DOT/OPSI-MS$^2$ of HG.**

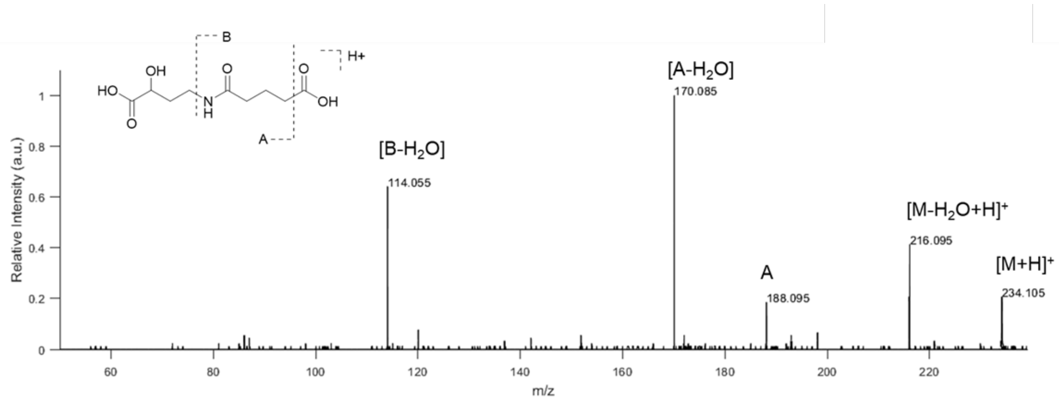


**Figure S83: I.DOT/OPSI-MS$^2$ of EG.**

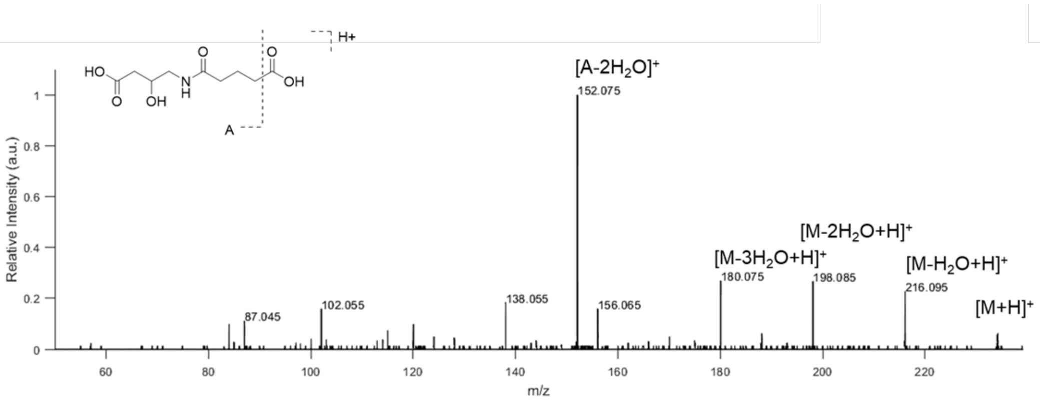


**Figure S84: I.DOT/OPSI-MS$^2$ of FG.**

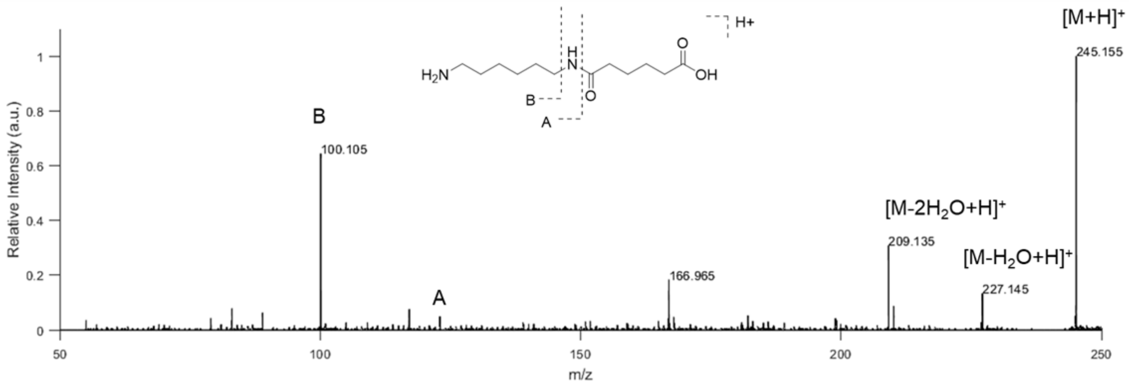


**Figure S85: I.DOT/OPSI-MS$^2$ of MA.**

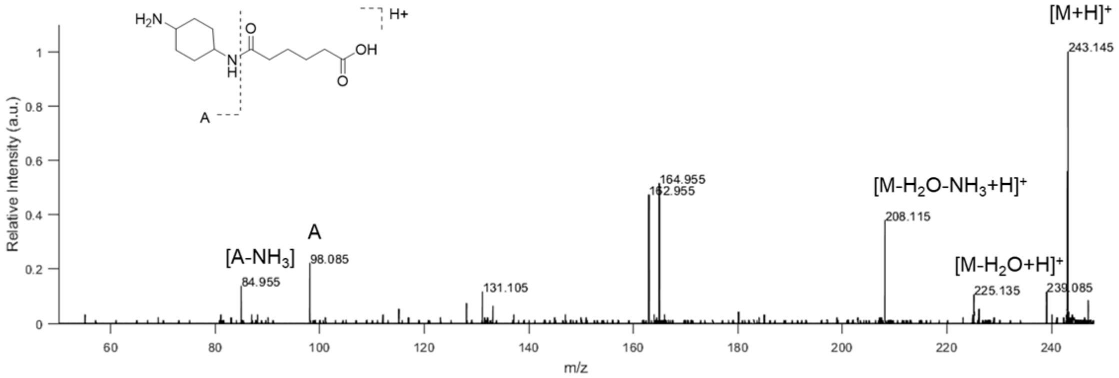


**Figure S86: I.DOT/OPSI-MS$^2$ of NA.**

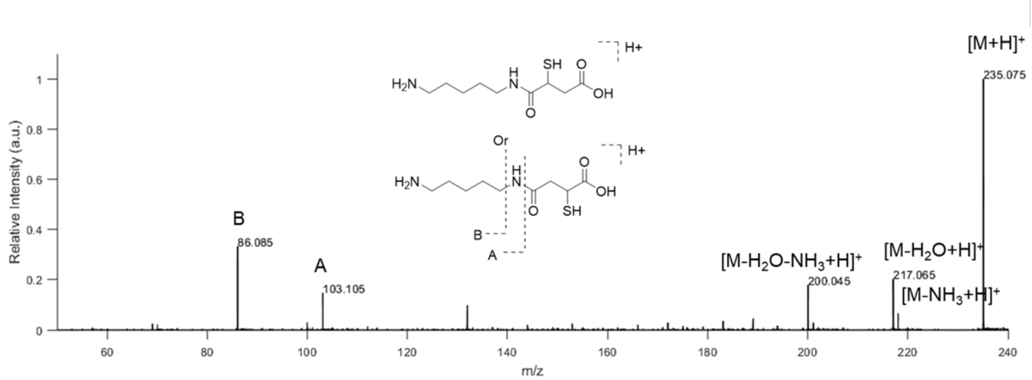


**Figure S87: I.DOT/OPSI-MS[2] of CP.**

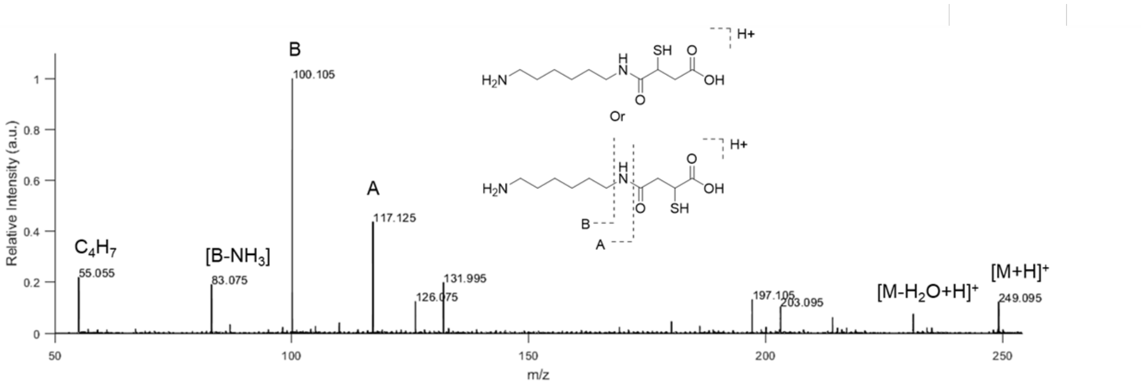


**Figure S88: I.DOT/OPSI-MS$^2$ of MP.**

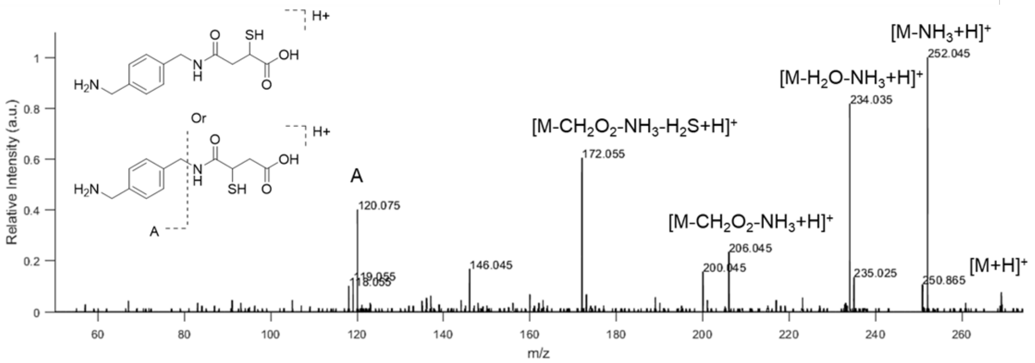


**Figure S89: I.DOT/OPSI-MS$^2$ of XP.**

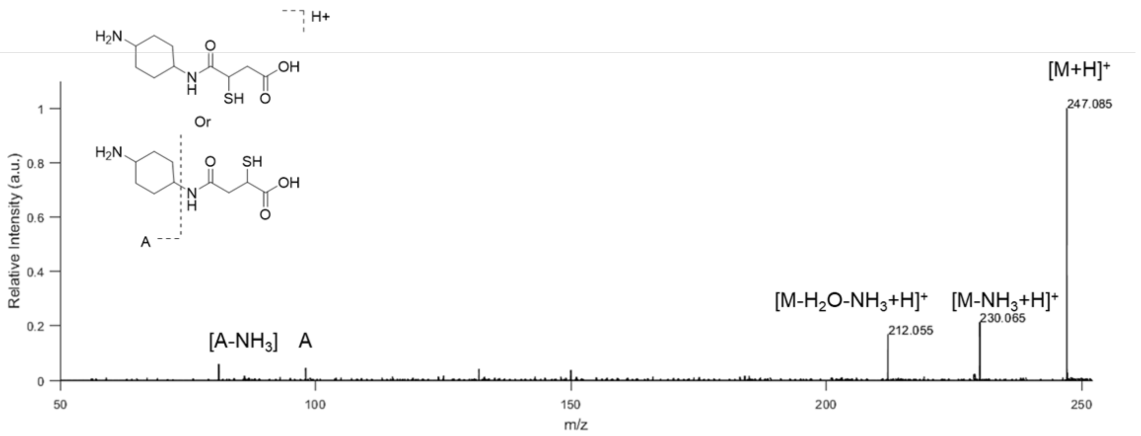


**Figure S90: I.DOT/OPSI-MS[2] of NP.**

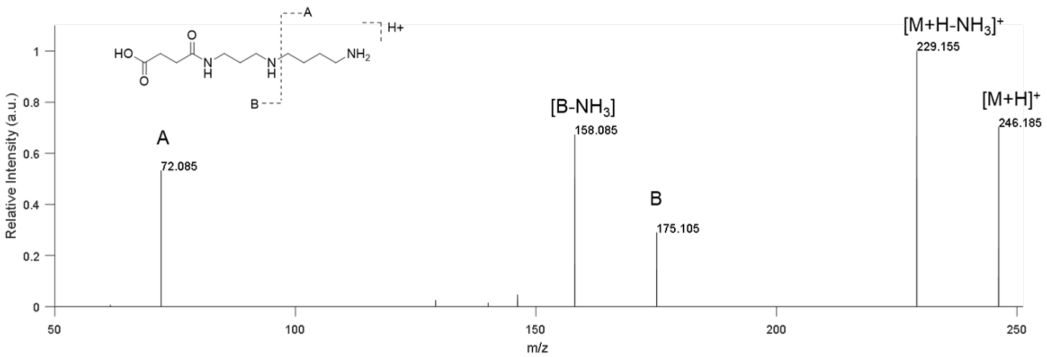


**Figure S91: I.DOT/OPSI-MS$^2$ of ligation product using succinic acid and spermidine with DdaG.** A collision energy of 30 eV was used.

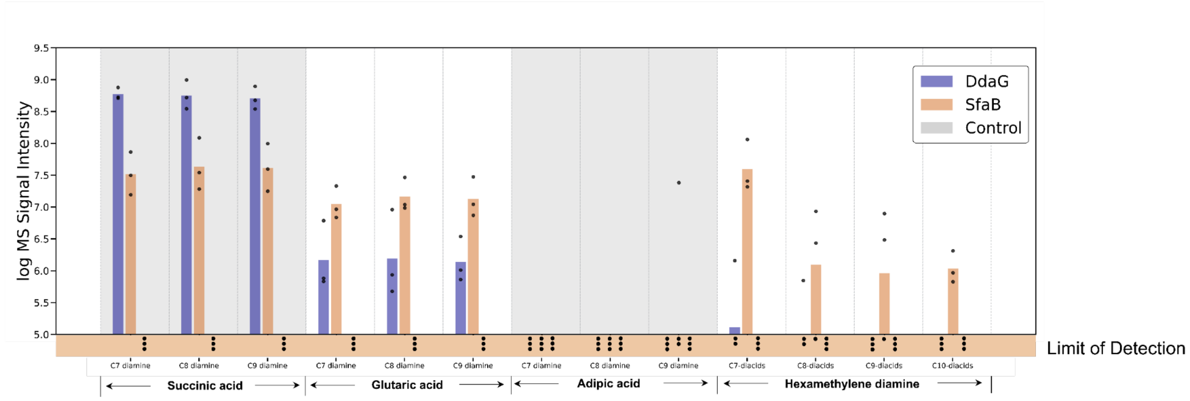


**Figure S92: Investigation of enzyme activities on diacid and diamine chain length** . *In vitro* biochemical assays were conducted by incubating 5 mM of different carbon lengths of linear diacids (i.e., C7-C10 diacids) with 5 mM hexamethylenediamine (**M**) or different carbon lengths of linear diamine (i.e., C7-C9 diamines) with 5 mM succinate (**S**) in the presence of 10 μM enzymes (i.e., DdaG or SfaB) or no-enzyme control at 30 °C for 16 h with shaking at 200 rpm. Products were assayed using OPSI-MS. The log MS signal intensity was cut off at 5.0 to filter out noise. Sample size is $N = 3$.

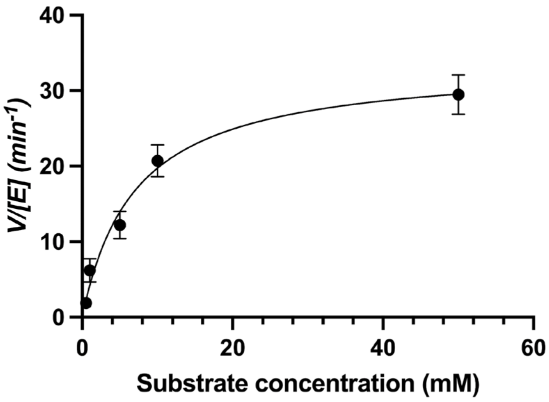


**Figure S93: Plots of *V*/E$_t$ versus [S] according to the Michaelis−Menten equation.** Kinetic study was conducted by incubating different concentrations of Succinate **S** and Hexamethylene diamine **M** with DdaG. Products were assayed using OPSI-MS. Sample size is $N = 3$. Error bars are shown as the mean ± standard deviation.

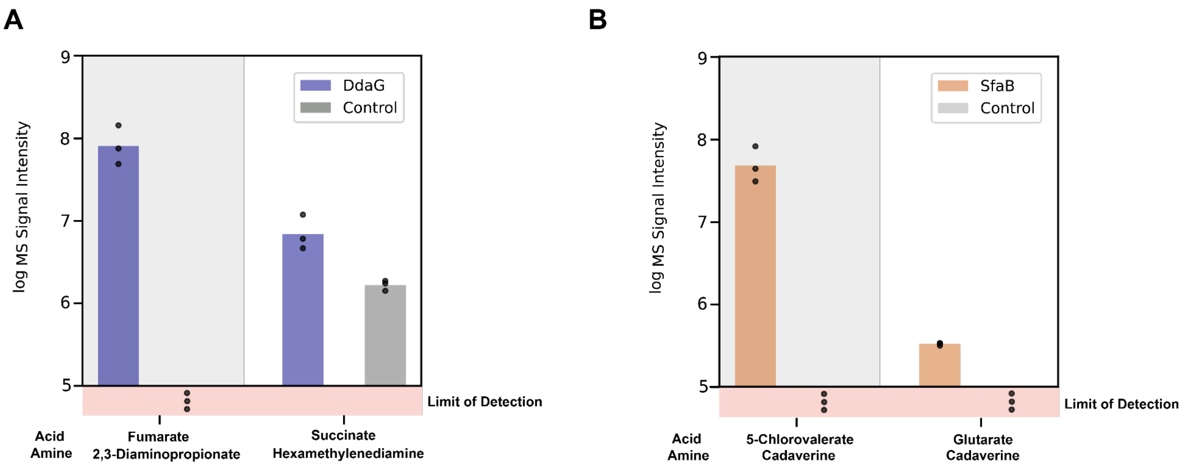


**Figure S94: Activities of amide synthetases using cell-free expression.** A) *In vitro* biochemical assays were conducted by incubating 25 mM of fumarate or succinate with 25 mM 2,3-diaminopropionate or hexamethylenediamine in the presence of cell-free expressed DdaG or no-enzyme control at 30 °C for 24 h with shaking at 200 rpm. B) Similar assays were performed using 25 mM 5-chlorovalerate or glutarate with 25 mM cadaverine and cell-free expressed SfaB or a no-enzyme control. DdaG or SfaB was expressed using NEBExpress® Cell-free *E. coli* Protein Synthesis System at 30 °C for 16 h. Products were assayed using OPSI-MS. The log MS signal intensity was cut off at 5.0 to filter out noise. Sample size is $N = 3$.
